# Human sodium current voltage-dependence at physiological temperature measured by coupling patch-clamp experiment to a mathematical model

**DOI:** 10.1101/2023.06.06.543894

**Authors:** Veronika O. Abrasheva, Sandaara G. Kovalenko, Mihail Slotvitsky, Serafima A. Scherbina, Aleria A. Aitova, Sheida Frolova, Valeria Tsvelaya, Roman A. Syunyaev

## Abstract

Voltage-gated sodium channels are crucial to action potential propagation in excitable tissues. Voltage-clamp measurements of sodium current are very challenging and are usually performed at room temperature due to the high amplitude and fast activation of the current. In this study, we measured sodium current’s voltage dependence in stem-cell-derived cardiomyocytes at physiological temperature. Although apparent activation and inactivation curves measured as the sodium current amplitude dependence on voltage step is within the range reported in previous studies, we demonstrate a systematic error in our measurements that is due to deviation of membrane potential from the command potential of the amplifier. We show how this artifact can be accounted for by the computer simulation of the patch-clamp experiment. This patch-clamp model optimization technique yields a surprising result: −11.5 mV half-activation and −87 mV half-inactivation of the sodium current. Although the half-activation is strikingly different from what was previously believed to be typical for the cardiac sodium current, we show that this estimate explains conduction velocity dependence on extracellular potassium in hyperkalemic conditions.

**Key points:**

- Voltage gated sodium currents play a crucial role in excitable tissues including neurons, cardiac and skeletal muscles.
- Measurement of sodium current is challenging because of its high amplitude and rapid kinetics, especially at physiological temperature.
- We have used the patch-clamp technique to measure human sodium current voltage-dependence in human induced pluripotent stem cell-derived cardiomyocytes.
- The patch-clamp data was processed by optimization of the model accounting for voltage-clamp experiment artifacts, revealing a large difference between apparent parameters of sodium current and the results of the optimization.
- We conclude that actual sodium current activation is extremely depolarized in comparison to previous studies.
- The new sodium current model provides a better understanding of action potential propagation, we demonstrate that it explains propagation in hyperkalemic conditions.

## Introduction

Voltage-gated sodium ion channel is one of the most important channels maintaining the excitability of neurons, muscles, and cardiomyocytes. In the vast majority of excitable tissues, the upstroke of an action potential (AP) is mostly determined by the fast sodium ion current (I_Na_). In the cardiac tissue, the predominant isoform is Nav1.5, encoded by the SCN5A gene (Kaufmann *et al*., 2013; Veerman *et al*., 2015). The role of sodium current makes accurate quantification of its dynamics necessary for an understanding of cardiac function and arrhythmogenesis. The conduction velocity is determined by sodium current (King *et al*., 2013), impaired I_Na_ function may result in arrhythmogenic unidirectional blocks (Starmer, 2003) or, conversely, reduce the probability of unidirectional block (Quan & Rudy, 1990), and mutations in SCN5A gene cause dangerous arrhythmias (Clancy & Kass, 2005).

However, the very properties of I_Na_ that underlie its function make it extremely challenging to measure and characterize. In particular, in the typical voltage-clamp setting, the high amplitude and rapid kinetics of I_Na_ make membrane potential uncontrollable (Nagatomo *et al*., 1998), as a result, most of the published experiments were carried out below physiological temperature (Hanck & Sheets, 1992; Sakakibara *et al*., 1993a; Sheets & Ten Eick, 1994; Nagatomo *et al*., 1998; Wagner *et al*., 2006; Nguyen *et al*., 2022). In general, AP mathematical models extrapolate the data to 37°C using a Q10 estimate for time constants or linear extrapolation for steady-state activation/inactivation curves (Tusscher *et al*., 2004; O’Hara *et al*., 2011). However, some peculiarities of modern mathematical models demonstrate the limitations of this approach: as an example, the widely used O’Hara-Rudy model is prone to conduction block even at mild hyperkalemic conditions (see online commentary to the O’Hara et. al. article (O’Hara *et al*., 2011)). The goal of the current study is to develop the pipeline making the accurate quantification of I_Na_ possible.

Traditional voltage-clamp experiments involve the measurement of the amplitude of current at different membrane voltages. The voltage is stepped from the holding potential, and the current transient is recorded. As a general rule, Hodgin-Huxley gating variables are regarded as “well-separated” by the experimental protocol, i.e., the time constants of the kinetics of activating and inactivating gates are rendered to differ by order of magnitude (Beaumont *et al*., 1993). A number of publications discussed that this “traditional” technique may result in large errors and proposed improved methods of parameter estimation based upon the minimization of objective function accounting for both activation and inactivation simultaneously (Willms *et al*., 1999; Lee *et al*., 2006). Due to the limitations of the voltage-step-based protocols, other techniques were proposed. For example, the so-called “Nonequilibrium Response Spectroscopy” was proposed by Millonas and Hanck, which is essentially a rapidly fluctuating large amplitude voltage clamp recorded using low-resistance pipettes (Millonas & Hanck, 1998). This technique was further developed lately with several studies proposing non-standard dynamic rapid-frequency voltage-clamp (Groenendaal *et al*., 2015; Clerx *et al*., 2019). and current-clamp (Groenendaal *et al*., 2015) protocols allowing a researcher to reduce the uncertainty of the model parameters. Moreover, recently (Lei *et al*., 2022), the technique of designing an optimal experimental protocol to calibrate cardiac electrophysiology models was proposed. However, the limitation of these non-standard protocols lies partly in the biological variability of the object under investigation: for example, vastly used human induced pluripotent stem cells derived cardiomyocytes (hiPSC-CM) represent a mixture of nodal-like, atrial-like and ventricular-like phenotypes (Ma *et al*., 2011; Karakikes *et al*., 2015; Argenziano *et al*., 2018; Burnett *et al*., 2021). This heterogeneity of cell culture requires “trained eye” quality control to make sure that the ionic current under investigation is indeed being expressed in the particular cell. While the dynamics of ionic currents during the longer voltage steps are, in general, well known, the high-frequency protocols might mask the experimental artifacts that would be apparent otherwise.

Another source of high variability in patch-clamp experiments is the limitation of the technique itself: while membrane potential is supposed to follow the command potential of the amplifier, there is actually a substantial difference both because of membrane capacitor charging time and voltage drop on series resistance (Moore *et al*., 1984; Strickholm, 1995a, 1995b; Sherman *et al*., 1999). Several techniques have been proposed over the years to overcome these undesirable artifacts, some of which were implemented in patch-clamp amplifiers. Most often, “supercharging” is used to accelerate membrane charging time by applying a large initial potential, thus boosting the command signal (Armstrong & Chow, 1987; Strickholm, 1995b). The series resistance voltage drop can be compensated by feeding back a portion of the recorded current (Strickholm, 1995a). These techniques, however, require careful manual adjustment by the patch-clamp operator since overcompensation results in current oscillation and seal disruption (Montnach *et al*., 2021). These problems came to the attention of the electrophysiological community once again lately (Lei *et al*., 2020; Montnach *et al*., 2021) with the emergence of high-throughput automated patch-clamp systems (Seibertz *et al*., 2022). In a study by Lei et al. (Lei *et al*., 2020), a novel technique of interpreting patch clamp data was used to fit the model of the hERG1a channel. In order to eliminate experimental variability, the authors constructed a model that described not only the ion current dynamics but the entire voltage-clamp experiment. We use a similar technique in our study to quantify fast sodium current at physiological temperature while using more traditional stepped potential protocols.

## Materials and methods

### IPSC differentiation

All studies were approved by the Moscow Institute of Physics and Technology Life Science Center Provisional Animal Care and Research Procedures Committee, Protocol \#A2-2012-09-02 and conducted in accordance with the Declaration of Helsinki. The cell line m34Sk3 is provided by the E. Meshalkin Novosibirsk Scientific Research Institute of Circulation Pathology, handling approved by the Institute of Circulation Pathology Ethics Committee (#27, March 21, 2013). Pluripotent stem cells line was generated from cells donated by the patient without reported cardiovascular diseases. Informed consent was obtained from the donor.

Fibroblast derivation, reprogramming to the pluripotent state, characterization of induced pluripotent stem cells (iPSC) line m34Sk3, direct differentiation of iPSCs into ventricular-like cardiomyocytes were previously described in the previous publications (Podgurskaya *et al*., 2019; Slotvitsky *et al*., 2020). iPSCs were cultivated for several passages under feeder-free conditions in the Essential 8™ Medium on Geltrex LDEVFree hESC-Qualified Reduced Growth Factor Basement Membrane Matrix (Thermo Fisher Scientific). 3–4 days before differentiation, iPSCs were plated on 24-well plates. Directed differentiation of iPSCs into cardiomyocytes was triggered at 80–90% cell density by adding the RPMI 1640 medium (Lonza) containing B27 supplement minus insulin (Thermo Fisher Scientific) and 8 μM CHIR99021 (SigmaAldrich) for 48h. Other differentiation steps were carried out according to the Gi-Wi differentiation protocol as described in (Lian *et al*., 2013).

Starting from day 9 of differentiation, the first cell contractions were observed in presence of RPMI 1640 medium (Lonza) with B27 supplement (Thermo Fisher Scientific). The flow cytometry data for cardiac markers cardiac troponin T (cTnT) and immunostainings for ventricle-type β-myosin regulatory light chain (MLC2), β-myosin heavy chain (MHC) and cardiomyocyte transcription factor NKX2.5 are reported previously (Podgurskaya *et al*., 2019). An increase in cx43 expression during directed differentiation was verified in (Slotvitsky *et al*., 2020, 2022), which corresponds to the literature data on the maturation of ventricular-like iPSC-derived cardiomyocytes (Tan & Ye, 2018). The efficiency of directed iPSC differentiation into cardiomyocytes estimated by the cTnT expression was 47% (Dementyeva *et al*., 2019). The differentiation yield for Gi-Wi protocol on m34Sk3 cell line proved to be stable during cell maturation (up to day 58) and has been characterized previously in (Slotvitsky *et al*., 2020).

### Patch-clamp samples preparation

To obtain samples for patch-clamp experiments, cells were disaggregated using TrypLE Express (1X) (Sigma) to a unicellular state according to the protocol described in (Burridge *et al*., 2014) at 25-30 days of differentiation. Cells were incubated for 5 minutes in 700ml of a disaggregating solution at physiological temperature in a CO_2_ incubator. A cell suspension was washed off from the original substrate with RPMI 1640 medium with 1 × B27 Supplement with 10 μM Y27632 (StemRD, USA). The suspension is centrifuged for 5 minutes at 200 g, all cells are precipitated. Then the supernatant (containing the medium and the disaggregating solution) was completely drained, the cell suspension was again mixed in RPMI 1640 medium with 1 × B27 Supplement with 10μM Y27632, after which all cells were plated on a new substrate. For the patch clamp studies, 13 mm glasses were sterilized and coated with Human Fibronectin (Imtek). The differentiated cells were seeded in a concentration of 10 000 cells/cm2 to obtain samples with single cells.

### Immunohistochemical staining

Samples were fixed for 10 min in 4% paraformaldehyde (Sigma-Aldrich, 158127), permeabilized for 10 min in 0.4% Triton-X100. Cells were further incubated for 30 min in blocking buffer (1% bovine serum albumin in phosphate-buffered saline, PBS), overnight at 4°C with primary antibodies and for 1 h at room temperature with secondary antibodies. Cells were washed twice for 15 min in PBS. Sarcomeric α-actinin mouse (Abcam, ab9465, working dilution — 1:100) was used as the primary antibody. Alexa Fluor 568 goat anti-mouse IgG (HþL) highly cross adsorbed (Thermo Fisher Scientific, A11031, working dilution — 1:400) was used as the secondary antibody. Nuclei were stained with DAPI (Vector Laboratories, Inc., Canada). Samples were analyzed and processed on a Zeiss LSM 710 confocal mi-croscope with Zen black 3.0 software (Carl Zeiss, Germany).

### Patch-clamp experiments

Whole-cell perforated patch-clamp recordings were performed at a physiological temperature (37°C). Amphotericin B (Sigma Aldrich) was used as a perforating agent dissolved in DMSO at a concentration of 0.24 mg/ml. The extracellular solution used for recording fast sodium current I_Na_ contained (in mM): NaCl 20, CsCl 120, CaCl2 1.8, MgCl2 1, D-glucose 10, HEPES 10, Nisoldipine 0.002, E-4031 0.0005, Ivabradine 0.003 (pH = 7.4 CsOH). The pipette was filled with the intracellular solution containing (mM): NaCl 10, CsCl 135, CaCl2 2, MgATP 5, EGTA 5, and HEPES 10 (pH = 7.2 CsOH).

The experiments were made using a patch-clamp setup consisting of the following main elements: digital converter Digidata 1440A (Axon Instruments, Inc., USA), Axopatch 200B amplifier (Axon Instruments, Inc., USA), MP-285 micromanipulator (Sutter Instrument), Olympus IX71 inverted microscope, HumBug noise isolator (A-M-Systems), anti-vibration platform (AVTT75). The temperature of the extracellular solution was maintained at 37° C in the chamber using an automatic temperature controller TC-324C (Warner Instruments). For the pipettes manufacture we used micropipette puller P-97 (Sutter Instrument), billets of borosilicate glass (BF150-86-10, Sutter Instrument), micro-diaper (Micro forge, MF-900, Narishige).

Pipettes were made from borosilicate glass with tip resistance averaging 3 MΩ. The pipette offset was adjusted to zero just before the formation of the gigaohm contact (GΩ). After forming the gigaohm resistance, the capacitive components were compensated using amplifier settings. The end of the membrane perforation with amphotericin B was determined by a change in the capacitive currents. The approximate perforation time by antibiotic was about 5 minutes. After the cell capacitance was compensated, the voltage-gated sodium channels were detected using the established stimulation protocol.

Current recordings were obtained by using standard activation and inactivation protocols (Fig. 1A,B). The activation protocol for I_Na_ was determined using a stimulating step protocol from −80 to +15 mV from a holding potential of −80 mV with a duration of 100 ms in 5 mV increments. The inactivation protocol for I_Na_ was carried out by applying a 50-ms pulse of −20 mV following a 500-ms stimulating prepulse from −140 to 0 mV in 5 mV increments (holding potential of −120 mV).

**Fig. 1.**
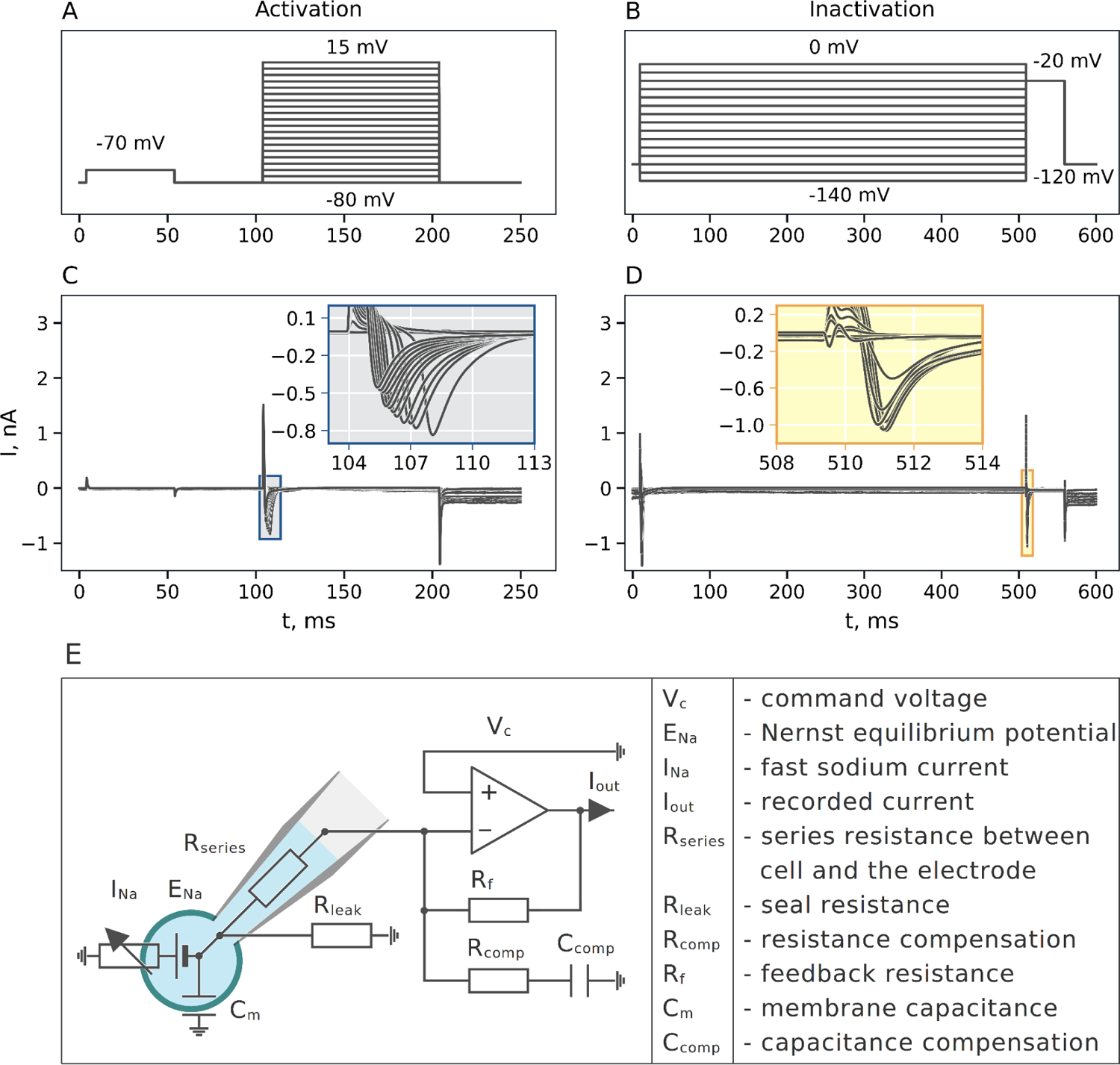
The voltage-clamp experiment: (**A**, **B**) Voltage-clamp protocol used to measure the activation and the inactivation. (**C, D**) Representative recordings of fast sodium current during an activation and inactivation voltage protocol. (**E**) The equivalent electrical circuit of the patch-clamp experiment model.

To ensure adequate voltage control in experiments, the peak sodium current was reduced by using the low extracellular sodium concentration ([Na^+^] *=* 20 mM), which results in *-18 mV* sodium reversal potential. The same technique was previously used by others (Ma *et al*., 2011). In addition, recordings of I_Na_ were made after about 10 minutes after perforation to minimize the effects of inadequate perforation. The membrane capacitance measured using the pCLAMP10.2 software ranged from 10 to 56 pF.

### Optical mapping

Optical mapping was carried out on the installation consisting of a high-speed EM-CCD video camera (Andor i-Xon 3, Andor Technologies), a mercury lamp (Olympus U-RFL-T) and optical microscope (Olympus MVX10 MacroView) with a bandpass filter cube (Olympus U-M49002XL). Periodic stimulation of samples was carried out with an impulse generator (Vellemann, PCGU-1000), a platinum point electrode (stimuli) and a reference circular electrode (reference electrode).

Optical mapping of human iPSC-derived cardiomyocytes occurred approximately at 50 days after the initiation of direct differentiation. The potential-sensitive dye FluoVolt™ Membrane Potential Kit (Thermofisher, USA, F10488) was used for the action potential recordings according to protocol described in (Slotvitsky *et al*., 2020). Fluo-4 AM, a calcium-sensitive dye, was used to measure the conduction velocity (CV) due to its higher signal amplitude and stability. The sample was incubated in a sterile medium at 37°C with the fluorescent calcium-dependent Fluo-4 AM (Invitrogen, USA) dye in concentration of 4 μg/ml for 30 min. Then the dye solution was exchanged with a sterile Tyrode’s solution. All experiments were carried out at a temperature of 37 °C. For the initiation of excitation waves, a looped 500-μm diameter, platinum wire reference electrode was placed onto the well. A point platinum wire electrode was placed ∼0.3 mm above the sample close to its edge. Monolayers were stimulated by 6 V square pulses of 20 ms duration and 1000 ms cycle length, CV was measured 1. 2 ± 0. 1 mm from the stimulating electrode.

### Model of experimental setup

In this study we use two different techniques for approximating the sodium current voltage-dependence. Firstly, we use a traditional technique of measuring current amplitude dependence on voltage step termed *apparent* activation/inactivation curves below. Sodium current is described via Hodgkin-Huxley formalism with three identical activating and two inactivating gates (Hodgkin & Huxley, 1952; Tusscher *et al*., 2004; O’Hara *et al*., 2011):

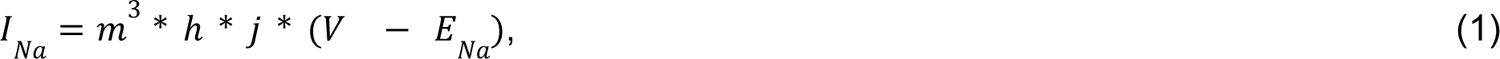

where m is the activating gating variable, h – fast inactivating gate, j – slow inactivating gate, while *j*_∞_ = ℎ_∞_.

Standard Hodgkin-Huxley differential equations described the gating variables dynamics.

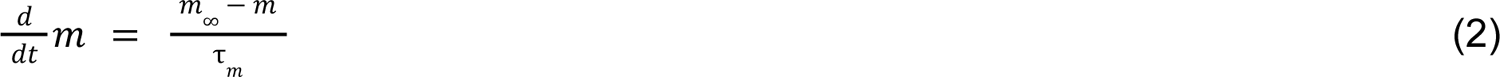

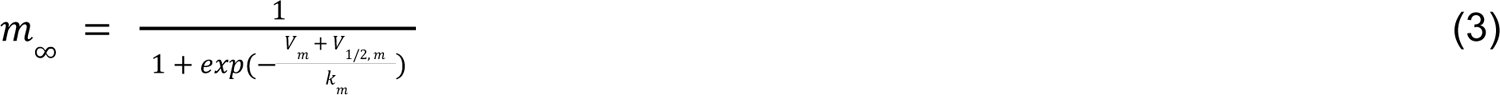

The steady-state model parameters for the *apparent* activation/inactivation were thus found by simple least-square fitting of the normalized current/voltage relationship to the following equations: equation (4) was used for the activation traces, equation (5) – for the inactivation traces. The Levenberg-Marquardt algorithm was used for the optimization technique.

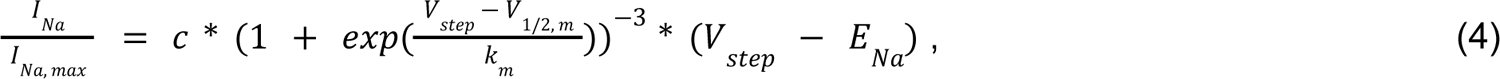

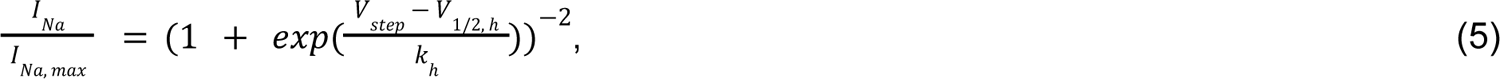

Secondly, to account for experimental artifacts, we recover parameters of sodium current via the optimization of the experimental setup model. The equivalent electrical scheme of the experimental setup that we used is shown in Fig. 1E. The full list of equations is given in the Supplementary Material (equations S1-S21). In this study, we mostly follow the set of differential equations developed by Lei et al. (Lei *et al*., 2020) apart from the compensation: while feedback current was used in the aforementioned study (Strickholm, 1995a) supercharging (Strickholm, 1995b) was used in our experiments.

### Genetic algorithms

The patch-clamp model parameters were optimized to match the patch-clamp current traces by custom Genetic Algorithm modification (Smirnov et al., 2020). During simulations, command potential dependence on time reproduced corresponding activation or inactivation experiments, Fig. 1A,B. The initial generation of organisms (N=512) was exposed to selection, mutation, and crossover for 600 generations to minimize the cost function (see below). A full list of model parameters and their boundaries is provided in the Table S1, full list of equations is provided in the Supplementary Material, equations S1-S21. Note that Nernst potential, E_Na_, was considered as a free model parameter since we used reduced 20 mM extracellular Na^+^ concentration. In this case, for example, a 5 mM intracellular concentration change would result in 20 mV Nersnt potential change, and unsettled [Na^+^]_i_ concentrations might thus have a significant effect on experimental recordings. The set of differential equations was integrated using the CVODE solver (Hindmarsh *et al*., 2005; Gardner *et al*., 2022) with 1e-13 relative error tolerance and two values of absolute error tolerance: 1e-8 for *m*, ℎ, *j* gating variables and 1e-12 for the remaining variables.

The cost function was calculated as:

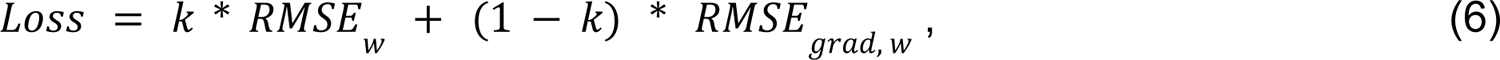

where *RMSE_w_*is the weighted root-mean-square error of the simulated output current, *i.e.;*

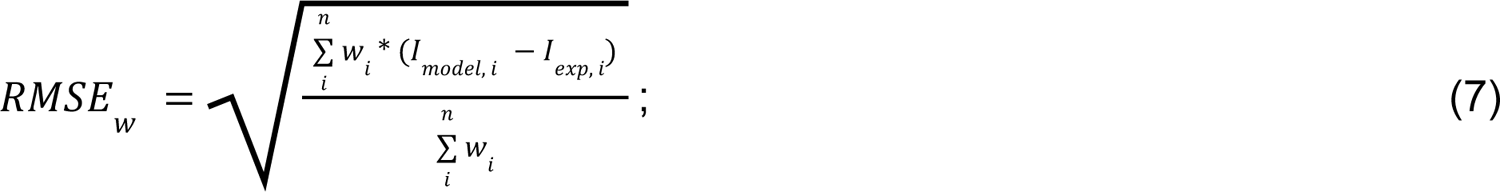

*RMSE _grad,w_*is the weighted root-mean-square error of derivative:

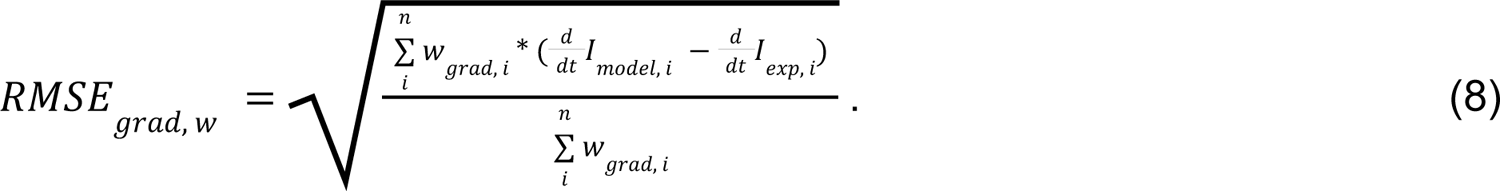

The weights (*w_i_*) were manually adjusted to increase the weight of sodium current transient and reduce the weight of the high-amplitude pipette charging transient (∼3nA). The reasons for using the derivative of the signal in the cost function are discussed below (see Results section).

### Uncertainty quantification

We used Markov Chain Monte-Carlo (MCMC) technique (William H. Press *et al*., 2007; Johnstone *et al*., 2016) in order to estimate parameter uncertainty. MCMC algorithm is a Bayesian technique used for sampling from posterior distributions of model parameters, *i.e.* for finding the probabilities of particular parameter estimates given the experimental data:

*P*(θ|*D*), where θ is the vector of model parameters, *D* is an experimental recording. In order to sample from posterior distribution one has to estimate the likelihood of the data: *P*(*D*|θ).

We used the Gaussian noise model for this estimate: 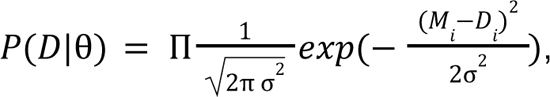 where *M_i_* corresponds to model predictions of the patch-clamp trace, *D* – to the experimental measurements. In our case, the derivative of the signal was closer to the actual sodium current than the current trace than the recorded signal itself (see “Patch-clamp model optimization” subsection of the “Results” section), thus the derivative of the signal was used for the *M_i_* and *D_i_* in the equation above. The noise variance - σ^2^, was considered as model hyperparameter and was sampled *via* Metropolis-within-Gibbs technique (William H. Press *et al*., 2007) from the inverse gamma distribution, which is the conjugate prior of the normal distribution (Smith, 2013): *P(σ^2^|θ,D*) = InvGamma(n/2, Σ(M_i_ − D_i_)^2^), where n is the number of data points. Delayed Rejection Adaptive Metropolis (DRAM) technique was used to sample from model parameters θ (Haario *et al*., 2006). MCMC sampling was implemented with a custom code based on pymc3 software (Salvatier *et al*., 2016). Probability density functions were estimated from the samples using kernels density estimation (Wand & Jones, 1994) with a Gaussian kernel function and 2 mV bandwidth. The highest posterior density credible intervals (CI) are reported throughout the text (Hespanhol *et al*., 2019).

Due to the sodium current’s high amplitude and short characteristic activation time, the model trace was very sensitive to parameters. For example, a 0.01% change in half-activation from the genetic algorithm solution resulted in the likelihood decrease by the order of 27, which in turn resulted in a very long Markov chain mixing time. To alleviate this issue, we have downsampled the signal (1:5 downsampling in inactivation and 1:10 in activation traces), and every second voltage step was omitted from the recording. 120 chains with 10000 iterations were used for sampling, the first 3000 iterations were eliminated as a burn-in period. The prior distributions for model parameters are given in the Table S2.

The model parameters θ can be divided into two parts: sodium current parameters, θ_M_, such as and *V_1/2_,m*; and *k_m_*; and experimental setup parameters, θ_E_, such as membrane capacitance and access resistance. While we suppose the former to be the same in each experiment, the latter depends on the particular cardiomyocyte. Ultimately, we are interested in the set of model parameters that explains every experiment available: *P*(θ_M_ | *D*1, *D*2, *D*3… *D*N), according to Bayes theorem:

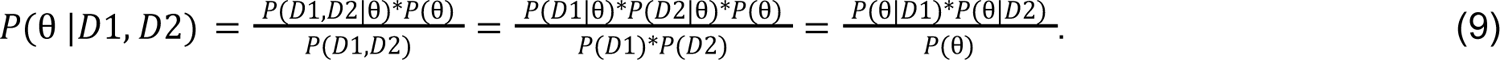

Moreover if θ corresponds to the *D*1 experiment θ – to the D2 experiment, we can write:

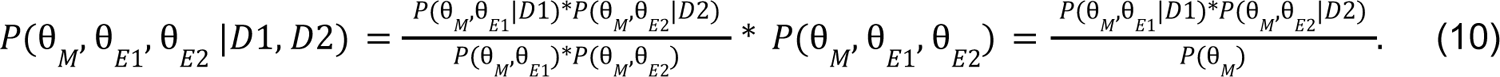

In the simplest case of noninformative prior of θ_E1_, i.e. when *P*(θ_M_) corresponds to a uniform distribution, by integrating out and θ_E1_ and θ_E2_, we can conclude that marginal posterior distribution of model parameters for a set of experiments can be found by simple multiplication of the marginal distributions for each experiment. Thus, to decrease computational time, instead of sampling model parameters using simultaneously all experimental recordings available, we sampled them separately. Furthermore, for the sake of simplicity, we treated unidentifiable parameters the same way we treated experimental setup parameters: these were not supposed to be the same in every experiment. In particular, only and *V_1/2,m_* and *k_m_* were regarded to be identifiable in the activation experiments, and *V_1/2,h_* and *k_h_* – in the inactivation experiments (see “Parameters uncertainty” subsection below on parameters identifiability).

### Simulation of hyperkalemic conditions

Action potential propagation in hyperkalemic conditions was simulated with monodomain equations coupled with single-cell electrophysiology described by the O’Hara-Rudy model (O’Hara *et al*., 2011) for ventricular-like tissue or Paci2020 model (Paci *et al*., 2020) for IPSC-CM tissue:

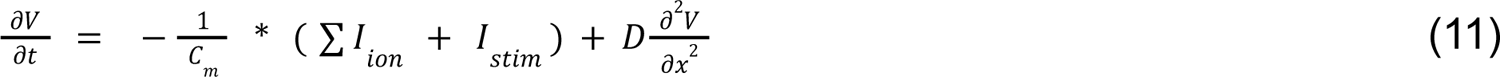

In the case of the O’Hara-Rudy model, an explicit four-point stencil scheme was used to solve partial differential equations. Paci2020 model equations were solved with OpenCARP software (Plank *et al*., 2021) using the Crank-Nikolson method for PDE, and explicit Euler for ODE. The Rush-Larsen technique (Rush & Larsen, 1978) was used for gating variables. The spatial grid step was set to 0.01 mm, and the temporal step was 0.005 ms. The diffusion coefficient, *D*, was set to adjust conduction velocity (CV) to 5.2 cm/s, corresponding to our measurements in the monolayers of IPSC-CM (non-hyperkalemic conditions, *i.e.,* extracellular potassium concentration [K+] = 5.4 mM). The length of the 1D string was set to 2 mm. The single-cell models were pre-paced for 1000 s to reach stationarity conditions. CV in hyperkalemic conditions was measured after 40 beats in the center of the 1D string. Further increase in the length of simulated tissue or the number of paced beats did not affect the CV. In the simulations with I_Na_ modified according to our measurements, the time parameters were lifted from corresponding O’Hara-Rudy and Paci2020 models. Since maximum diastolic potential was above −60 mV in severe hyperkalemic conditions, the following modification was introduced to the calcium-dependent calcium inactivation of ICaL in the Paci2020 model:

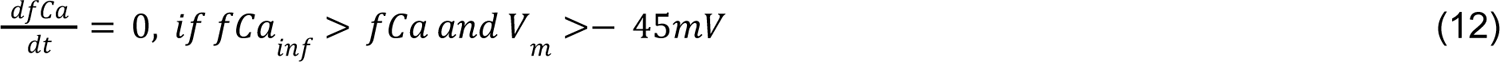

The same conditional statement was used in the original model, but the threshold potential for resetting *fCa* was −60 mV.

## Results

### Stem-cells derived myocytes phenotype characterization

Fig. 2A shows representative imaging of a cardiomyocyte’s cytoskeleton stained with alpha-actinin and nuclei with counterstain by DAPI. Alpha-actinin staining demonstrates clear striations within cells with a longitudinal size of approximately 55 ± 14 μm. The optical mapping demonstrated ventricular-like AP with a distinct plateau phase (Fig. 2B). The AP duration at 80% repolarization (APD80) was equal to 364 ± 21 ms (n=5), which corresponded both to native myocytes isolated from human ventricles (372 ± 16 ms (Kang *et al*., 2017)) and hIPSC-derived ventricular type myocytes (332 ± 38 ms (Shaheen *et al*., 2018)).

**Fig. 2.**
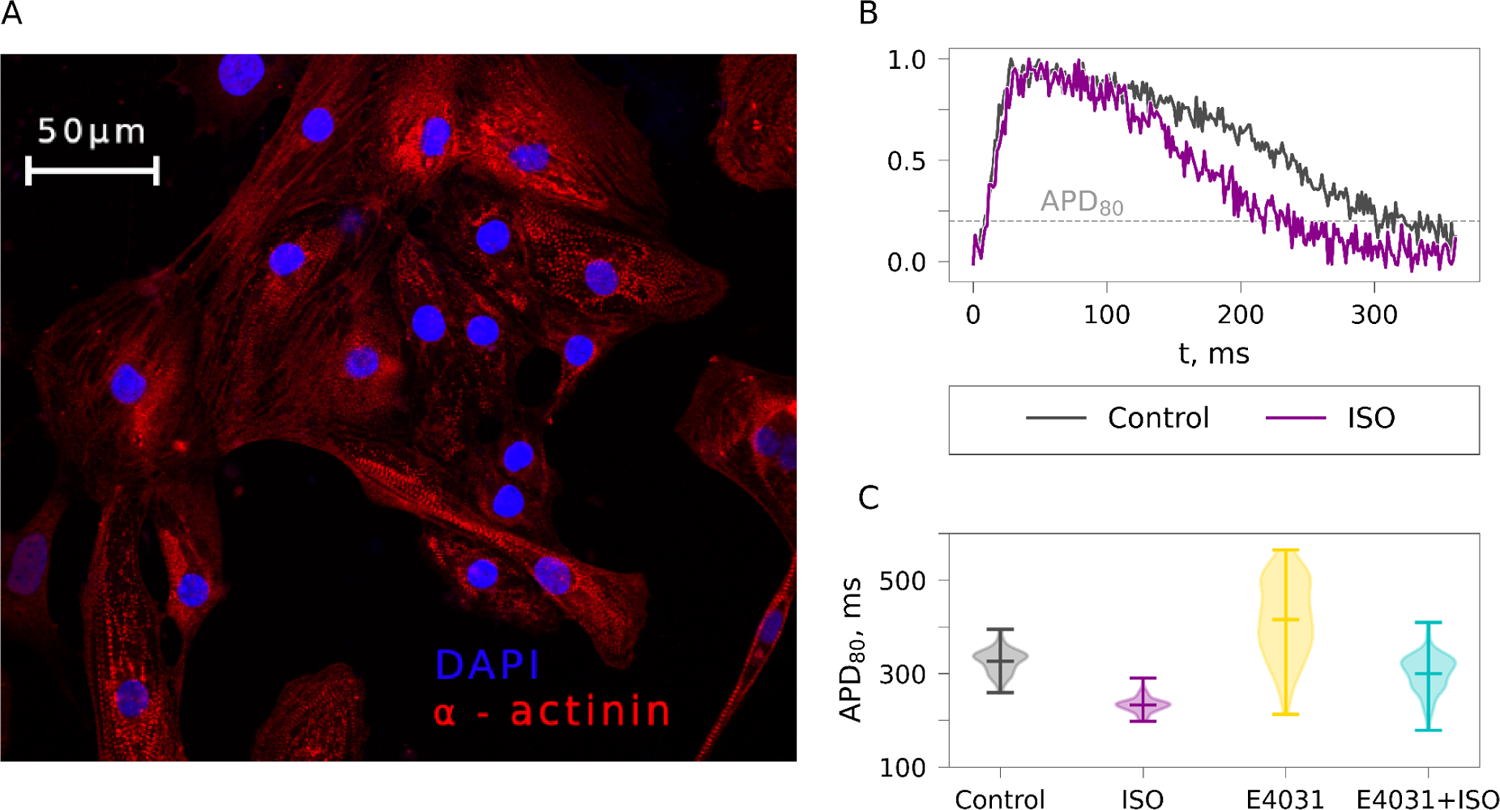
IPSC-derived myocytes characteristics. **(A)** Immunohistochemical staining of iPSC-CMs: α-actinin (red) and DAPI (blue). **(B)** Optical action potential in control (black) and isoprenaline (violet). **(C)** APD_80_ in control/isoprenaline/IKr block by E4031.

The maturity of iPSC-CMs as well as heterogeneity within a single monolayer (Fig. 2C) and across monolayers (Fig. S1) was further addressed by testing the effect of beta-adrenergic stimulation and selective I_Kr_ channel block. The 1 μm isoprenaline (ISO) superfusion caused a shortening of AP to 27% (from 364 ± 21 to 267 ± 15 ms, n=5) (Fig. 2B and Fig. S1), while a somewhat smaller 14% shortening was previously reported in wedge preparations at 100 nm isoprenaline (Kang *et al*., 2017). Fig. 2C shows the distribution of APD80 within a monolayer of iPSC-CMs: control (APD_80_ = 315 ± 30 ms, black), after 1 μM ISO superfusion (APD_80_ = 234 ± 18 ms, purple). The block of the IKr by 0.5 mkM E-4031 was very heterogeneous, prolonging APD_80_ to 416 ± 82 ms (Fig. 2C, yellow). Similarly, 70 ms APD_90_ prolongation by E-4031 was reported previously in isolated native ventricular cells (Jost *et al*., 2005), while much larger 70% APD_80_ prolongation was reported in human ventricular wedges (Holzem *et al*., 2016). Application of isoprenaline over E-4031 reversed APD prolongation (APD_80_ = 277 ± 51 ms, blue) indirectly confirming the expression of isoprenaline-sensitive I_Ks_ (Volders *et al*., 2003; Jost *et al*., 2005; Szentandrássy *et al*., 2012; Banyasz *et al*., 2014; Kang *et al*., 2017). We can thus conclude that despite the heterogeneity of iPSC-CM within monolayer (Fig. 2C), the general phenotype is similar to native ventricular myocytes.

The isolated IPSC-CMs membrane capacitance measured by Axopatch200B amplifier was 26.0 ± 14.0 pF (n = 58), which is much smaller than the capacitance of adult human ventricular cardiomyocytes of 194 ± 13 pF (Sakakibara *et al*., 1993a). The small cell size may indicate partial immaturity of cardiomyocytes and makes the patch-clamp experiments technically challenging (Wilson *et al*., 2011).

### Voltage-clamp experiments

First, we have measured the peak current/voltage relationship for activation (Fig. 1 A,C) and inactivation (Fig. 1 B,D). The resulting *apparent* activation/inactivation curves were found by fitting the normalized peak current to the sodium current model (Fig. 3A,B, black). The half-activation voltage and slope factor were equal to −45.6 ± 4.9 mV and 4.4 ± 3.4 mV respectively (n = 13), which is similar to Sakakibara et. al. (Sakakibara *et al*., 1993a) measurements in native ventricular myocytes (−42.3 ± 1.7 mV half-activation) but shifted in comparison to measurements reported in (Valdivia *et al*., 2005; Barajas-Martínez *et al*., 2013) (see also Table 1). The half-inactivation voltage was equal to −69.7 ± 7.3 mV, the inactivation curve slope was 11.6 ± 6.5 mV (n = 13). The half-inactivation measurements in native ventricular myocytes are within the range of −90 – −100 mV (Sakakibara *et al*., 1993a; Valdivia *et al*., 2005; Barajas-Martínez *et al*., 2013), but measurements in iPSC-CM are typically close to our measurements (Satin *et al*., 2004; Ma *et al*., 2011; Terrenoire *et al*., 2013). Half-activation and half-inactivation are compared to published data in Table 1.

**Fig. 3.**
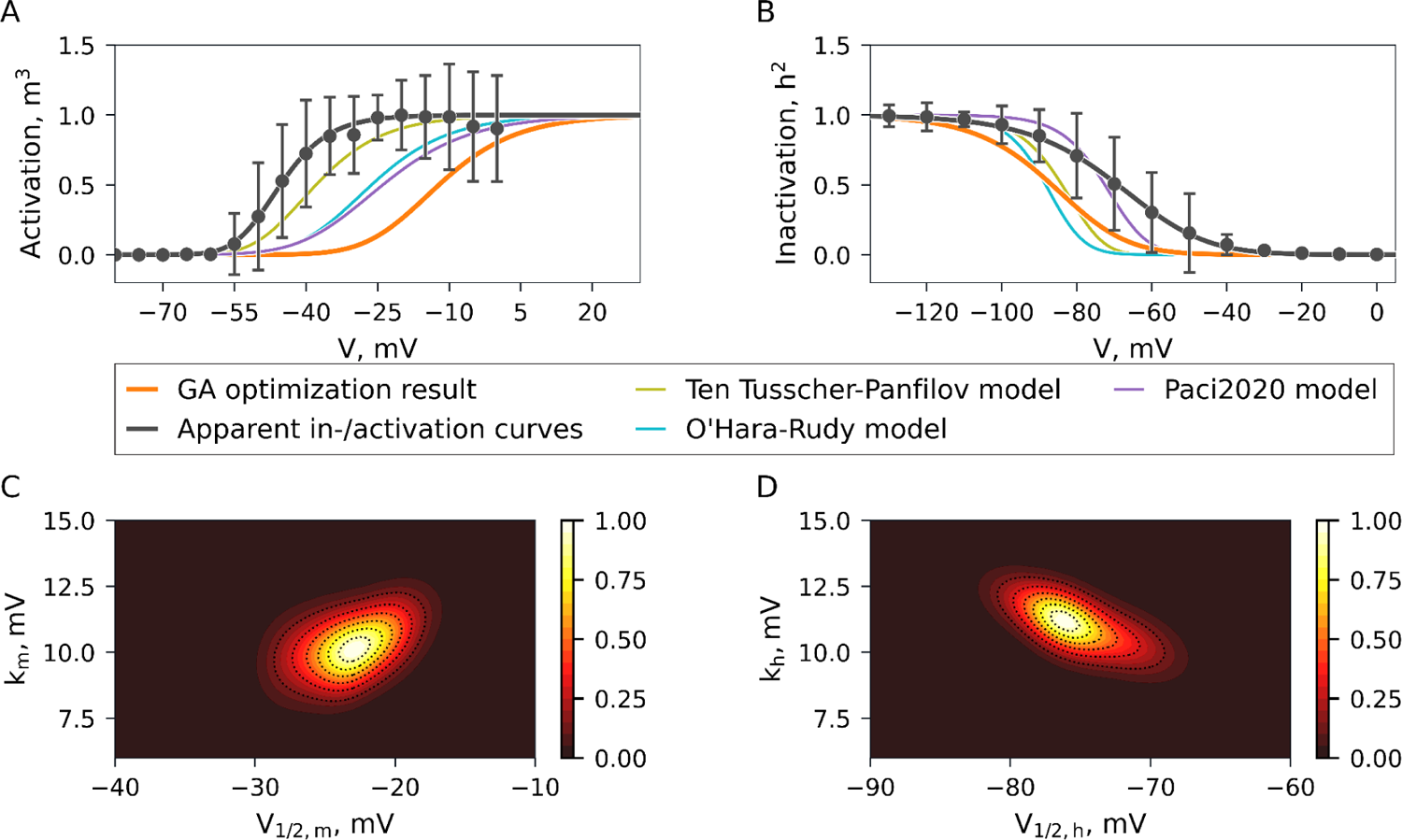
Activation\inactivation model I_Na_ parameters and their uncertainty. **(A, B)** Activation and inactivation parameters measured in this study (black – *apparent* values, orange – patch-clamp model optimization) compared to (Tusscher *et al*., 2004) model (green); (O’Hara *et al*., 2011) model (blue) and (Paci *et al*., 2020) model. **(C, D)** MCMC posterior distributions of I_Na_ parameters.

**Table 1.** Comparison of INa voltage-dependence in literature.

|  | *Half activation, mV* | *Half inactivation, mV* | Temperature | Condition |
| --- | --- | --- | --- | --- |
| This study, iPSC-CM |  |  |  |  |
| Apparent | $-45.6 \pm 4.9$ (n = 13) | $-69.7 \pm 7.3$ (n = 13) | 37± 1°C | |
| Model optimization | -12 ± 12 (n = 13) | -87± 12 (n = 13) | 37± 1°C | |
| MCMC, MLE [95%CI] | -10 [-16.5, -3.0] | -86 [-91.5, -77.0] |  |  |
| iPS-CM |  |  |  |  |
| (Ma *et al.*, 2011) | $-34.1$ (n = 5) | $-72.1$ (n=5) | 35–37°C | |
| (Terrenoire *et al.*, 2013) | $-25.0 \pm 0.3$ (n = 3) | $-70.3 \pm 1.7$ (n = 3) | 23–25°C | |
| (Terrenoire *et al.*, 2013) | $-25.0 \pm 0.3$ (n = 3) | $-68.9 \pm 0.9$ (n = 6) | 23–25°C | |
| (Selga *et al.*, 2018) |  |  |  |  |
| Monolayer-based differentiation | $-32.96 \pm 0.79$ (n = 11) | $-48.80 \pm 0.79$ (n = 8) | Room temperature | |
| EB-based differentiation | $-44.15 \pm 0.37$ (n = 3) | $-61.48 \pm 0.5$ (n = 3) | Room temperature | |
| Embryonic stem cell derived cardiomyocytes |  |  |  |  |
| (Satin *et al.*, 2004) | $-30$ (n = 21) | $-72.6 \pm 0.7$ (n = 19) | 26–29°C | |
| (Jonsson *et al.*, 2012) | $-34$ (n = 7) | $-78$ (n = 7) | 20°C | |
| HEK |  |  |  |  |
| (Nagatomo *et al.*, 1998) | $-43.9 \pm 1.0$ (n = 6) | $-39.6 \pm 2.2$ (n = 8) | 23°C | |
| (Nagatomo *et al.*, 1998) | $-50 \pm 1.1$ (n = 17) | $-85.8 \pm 2.7$ (n = 7) | 33°C | |
| (Nguyen *et al.*, 2008) | $-47.8 \pm 0.5$ (n=10) | $-89.4 \pm 0.8$ (n=10) | ~22°C | WT |
| Native cardiomyocytes |  |  |  |  |
| (Sakakibara *et al.*, 1992) | $-38.9 \pm 0.9$ (n = 46) | $-95.8 \pm 0.9$ (n = 46) | 17 ± 1°C | mixed |
| (Sakakibara *et al.*, 1993b) | $-42.3 \pm 1.7$ (n = 12) | $-99.8 \pm 2.1$ (n = 12) | 17 ± 1°C | Congenital heart disease (CHD) |
| (Sakakibara *et al.*, 1993b) | $-43.8 \pm 0.2$ (n = 10) | $-94.5 \pm 2.3$ (n = 10) | 17 ± 1°C | Cardiomyopathy (CM) |
| (Valdivia *et al.*, 2005) | $-51 \pm 1.0$ (n = 11) | $-102 \pm 16$ (n = 3) | Room temperature | Non-failing human ventricular tissue |
| (Valdivia *et al.*, 2005) | $-50 \pm 1.1$ (n = 17) | $-88 \pm 1.9$ (n = 10) | Room temperature | Heart failure |
| (Barajas-Martínez *et al.*, 2013) | $\sim -52$ | $-90.1 \pm 0.9$ (n = 3) | Room temperature | Left ventricular hypertrophy |

### Patch-clamp model optimization

Secondly, experimental setup model optimization was used to recover the sodium current parameters. The necessity to simulate a voltage-clamp experiment is motivated by the fact that although ideally membrane voltage is supposed to be equal to command potential in a path-clamp setting, it is in fact subject to several undesirable artifacts.

1. Due to finite membrane capacitance, membrane potential changes exponentially with characteristic time *R* * *C_m_* in response to step command potential change.
2. Due to the high amplitude of sodium current the voltage on the cell membrane differs from the voltage on the amplifier because of the voltage drop on series resistance (*R_series_* * *I_Na_*) (Montnach *et al*., 2021).
3. The output current (*I_out_*) is not equal to the actual sodium current (*I_Na_*) due to the leak current through the patch-clamp seal (*I_leak_* = *V_m_/R_leak_*

These artifacts are usually partially compensated during the experiment, which is also accounted for by the experimental setup model (Fig. 1E). We based the patch-clamp setup model on the set of equations given in (Lei *et al*., 2020) (see Methods for details and the Supplementary Material, equations S1-S11 for the full list of equations).

The representative result of model optimization shown in Fig. 4 demonstrates the deviation between actual I_Na_ and recorded current, I_out_. As shown in Fig. 4A, after the voltage step to −10 mV the cell membrane depolarizes to the threshold potential over the first 5 ms, the subsequent sodium current activation results in the depolarization of the membrane above the command potential, and, after sodium current inactivation transmembrane voltage returns to the command potential of −10 mV. As shown in Fig. 4B the recorded current, I_out_, in this case, would deviate from the actual sodium current: in particular, the capacitive current, I_C_, and the I_Na_ negate each other, reducing the amplitude by the order of 10 and changing the dynamics of the output current. On the other hand, the case of 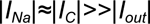 implies that 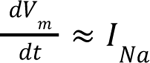 and 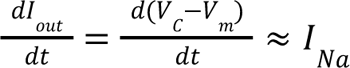
and

**Fig. 4.**
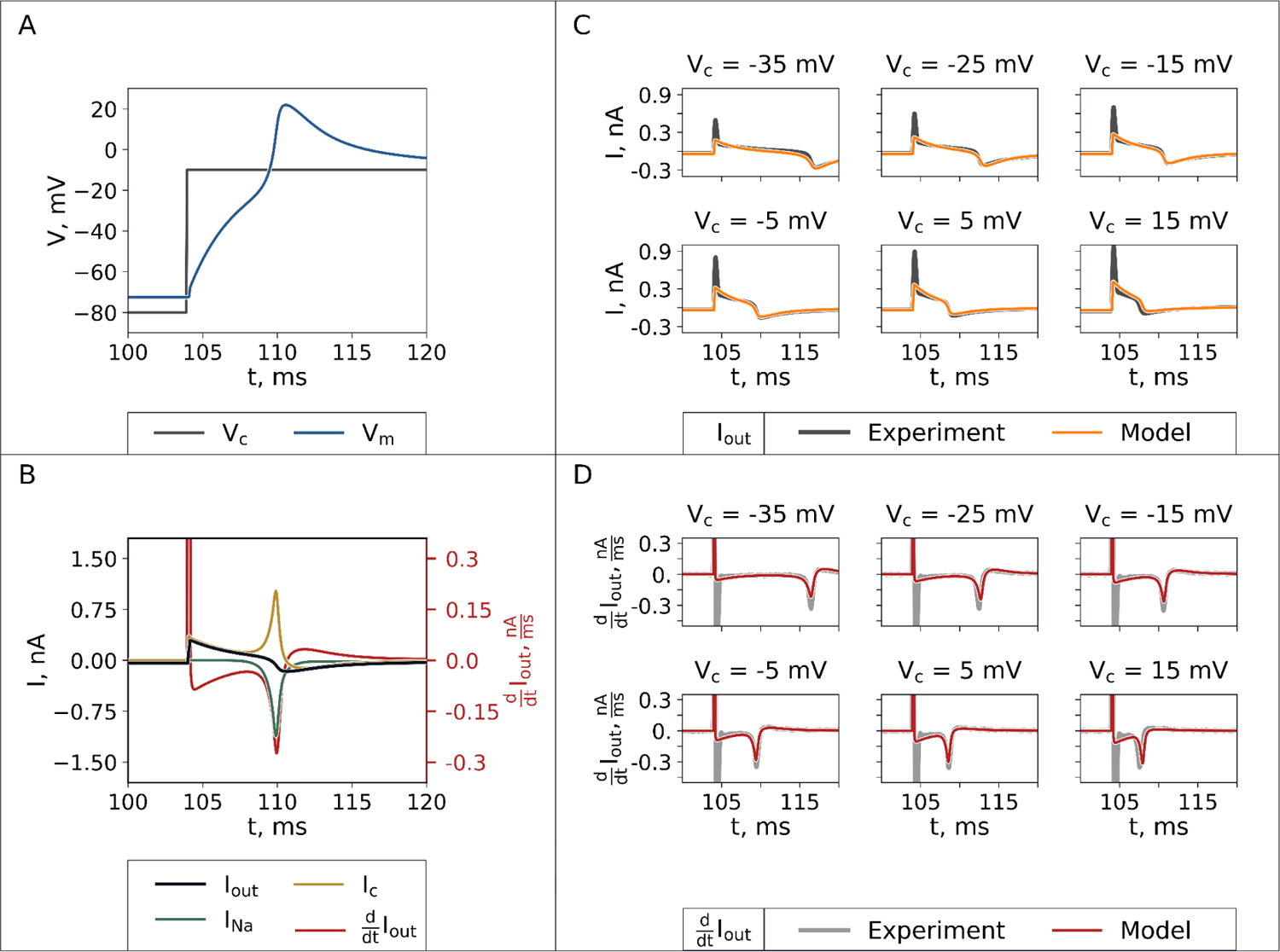
The model accounts for voltage-clamp experimental artifacts. **(A)**The comparison between command potential (*V*_*c*_) and computed cell membrane potential (*V*_*m*_). (**B**) Sodium current, I_Na_, and capacitive current, I_c_, in comparison with output current, I_out_, and its derivative. (**C, D**) The representative example of model optimization by a genetic algorithm: the model reproduces the experimental trace and its derivative.

Indeed, as shown in Fig. 4B, output current deviates from the sodium current, but the derivative of the output signal closely reproduces the actual ionic current dynamics. Therefore, in order to improve the convergence of the genetic algorithm, we used a combination of the signal and its derivative RMSE as the cost function during the optimization (see also “Materials and methods” section).

As shown in Fig. 4C,D, and in the Figures S7-S32 both the signal and its derivative were accurately reproduced by the optimized model, however, the amplitude of the latter was in general underestimated by the model (on average by 10% of the peak current). The RMSE of the activation experiments was 34 ± 14 pA for I_out_, 21 ± 7 pA/ms for dI/dt (for comparison the RMSE is 12 pA and 11 pA/ms in Fig. 4C,D correspondingly). The RMSE of the inactivation experiments was 55 ± 23 pA for I_out_, 16 ± 8 pA/ms for dI/dt.

The membrane capacitance parameter, C_m_, estimated by the optimization algorithm coincided with the measurement by Axopatch amplifier (n=26): 20±10 pF vs 27.0±15.0 pF respectively. In general access resistance was very large in our experiments: 120 ± 70 MOhm. In the particular case shown in Fig. 4 uncontrollable membrane voltage is due to the fact that access resistance and sodium current conductivity were both very large, 185 MOm and 400 mS/μF correspondingly. Although the former could be reduced by longer antibiotic perforation time, up to 30 minutes is required in order to achieve a seal of few MOhm (Kyrozis & Reichling, 1995). We have found long perforation times to be impractical at physiological temperature because of the unstable seal.

As shown in Fig. 3A the activation parameters we found as the result of optimization are strikingly different from the *apparent* activation curve. V_1/2,m_ was equal to −25.5 ± 7.8 mV, k_m_ – to 9.9 ± 3.1 mV; for comparison, the same parameters obtained from *apparent* activation by least-squares fit were −53.7 ± 5.2 mV and 6.2 ± 5.3 mV correspondingly. Note that we report V_1/2,m_ model parameter in this subsection, but half-activation in the “Voltage-clamp experiments” subsection above, Fig. 3 and Table 1; the latter is more common in experimental studies. Because of the third power of the gating variable, half-activation corresponds to ∼80% activation of the corresponding gate. In Fig. 3A we also show sodium current formulation lifted from the O’Hara-Rudy (O’Hara *et al*., 2011) and Ten Tusscher-Panfilov (Tusscher *et al*., 2004) ventricular models and iPSC-CM Paci2020 model (Paci *et al*., 2020) for comparison. Ventricular models rely on experiments at a sub-physiological temperature (Sakakibara *et al*., 1993a; Nagatomo *et al*., 1998) and have V_1/2,m_ of −39.57 mV and −56.86 mV, correspondingly. Paci2020 I_Na_ model is based on the earlier (Koivumäki *et al*., 2018) model, which, in turn, is based on (Ma *et al*., 2011) experiments and uses V_1/2,m_ of −39 mV.

The uncontrollable membrane potential has the opposite, depolarizing effect on the inactivation curve (Fig. 3B). Full-model optimization resulted in *V_1/2,h_=-76.0 ± 9.0* and *k_h_ = 12.0 ± 3.0 mV* against apparent values of *-56.6 ± 9.7 mV* for and *14.1 ± 7.6 mV* correspondingly. As one can see in Fig. 3B and Table 1 the optimization shifted inactivation to the values typical for ventricular cells, while the depolarized *apparent* inactivation curve is typical for iPSC-CM. For example, *V_1/2,h_=-66.5 mV* is used in the Paci2020 model.

### Parameters uncertainty

The difference between *apparent* activation/inactivation curves and the result of optimization implies a systematic error in experimental recordings: as described above, partial activation of sodium current depolarizes the membrane in comparison to command potential and results in further activation. However, two interrelated questions remain: firstly, the optimization algorithm itself might introduce systematic error; secondly, the uncertainty and identifiability of model parameters are unclear.

First, we tested the optimization algorithm on synthetic traces generated by the patch-clamp model (a set of parameters for synthetic data is given in the Table S1) with 4 pA Gaussian noise corresponding to experimental noise at the −80 voltage step. Genetic algorithm output parameters are shown in the Fig. S2,S3. We can conclude that steady-state activation and inactivation parameters as well as several important experimental artifact parameters, such as access resistance and membrane capacitance, are identifiable by genetic algorithm. In particular, *V_1/2,m_* was equal to −25±7 mV (n=39) in comparison to the input parameter of *-29 mV*; and k_m_=11±2 mV in comparison to 9 mV. Inactivation traces yielded similar results: V_1/2,h_=-79±9 (n=38) in comparison to −77 mV input value, k_h_=12.3*±*1.4 mV in comparison to 11 mV. On the other hand, ionic channel kinetics parameters, *i.e.* activation/inactivation time constants, were poorly identifiable from the experimental protocol. For example, τ_m_ at −40 mV was equal to 0.6*±*0.4 μs in comparison to the input value of 0.7 μs; the peak τ_h_ was 0.14 *±*0.12 ms in comparison to the true value was 0.08 ms.

Actual posterior distributions of model parameters given the experimental data were estimated *via* the MCMC technique as described above in the corresponding Materials and Methods subsection (“Uncertainty quantification”). The example of MCMC sampling for a particular experiment is shown in the Fig. S4; the recording itself is shown in Fig. S9. The signal was strongly downsampled in order to decrease mixing time and the likelihood function accounted for the trace derivative, instead of the recorded current itself (see Fig. 4B and discussion above in “Patch-clamp model optimization subsection”), but standard deviations of MCMC parameter samples were similar to synthetic genetic algorithm runs above: in the particular case shown in Fig. S4, *V_1/2,m_=23±7*, *k_m_=9±3 mV*. Finally, posterior distributions accounting for all experimental data available estimated using the eqn (10) above are shown in Fig. 3C (activation traces) and Fig. 3D (inactivation traces). The maximum likelihood estimator (MLE) and 95% credible intervals (CI) for steady-state activation/inactivation parameters are reported in Table 2.

**Table 2.** Sodium current parameters uncertainty as estimated by Markov-chain Monte-Carlo technique.

| Parameter | MLE | 95% CI |
| --- | --- | --- |
| $V_{1/2,m}$ | -23 mV | [ -28.6, -17.9 ] |
| $k_m$ | 10 mV | [ 5.6, 14.8 ] |
| $V_{1/2,h}$ | -76 mV | [ -80.9, -68.7 ] |
| $k_h$ | 11.25 mV | [ 9.3, 12.9 ] |

### AP propagation in hyperkalemic conditions

Previously, biphasic dependence of CV on [K^+^]_o_ was reported in guinea pigs (Weiss *et al*., 2017; King *et al*., 2021). At lower potassium concentrations the difference between resting membrane potential and activation threshold is larger, resulting in decreased CV. On the other hand, in hyperkalemic conditions, the CV is low because of the partial inactivation of the sodium channel. The measurement of the CV dependence on extracellular potassium concentration can thus indirectly verify or disprove sodium current voltage-dependence measurements.

Our measurements in the iPSC-CM monolayers (Fig. 5C) demonstrated 5.2 ± 1.8 cm/s CV in normokalemic conditions ([K^+^]_o_=5.4 mM). This value is low in comparison to ∼60 cm/s typical to human hearts (Glukhov *et al*., 2012; Aras *et al*., 2018), but similar to measurements in the monolayers (Zhu *et al*., 2017) and is explained by the absence of organized fibers in iPSC-CM monolayers. We did not observe any change of CV in hypokalemic conditions: at the 2.7 mM [K^+^]_o_ the CV was equal to *4.9 ± 2.2 cm/s*. Increased potassium concentration resulted in a gradual decrease of the CV to 0.9 ± 0.8 cm/s at [K^+^]_o_ = 14 mM and it was impossible to pace the tissue at [K^+^]_o_>14 mM.

**Fig. 5.**
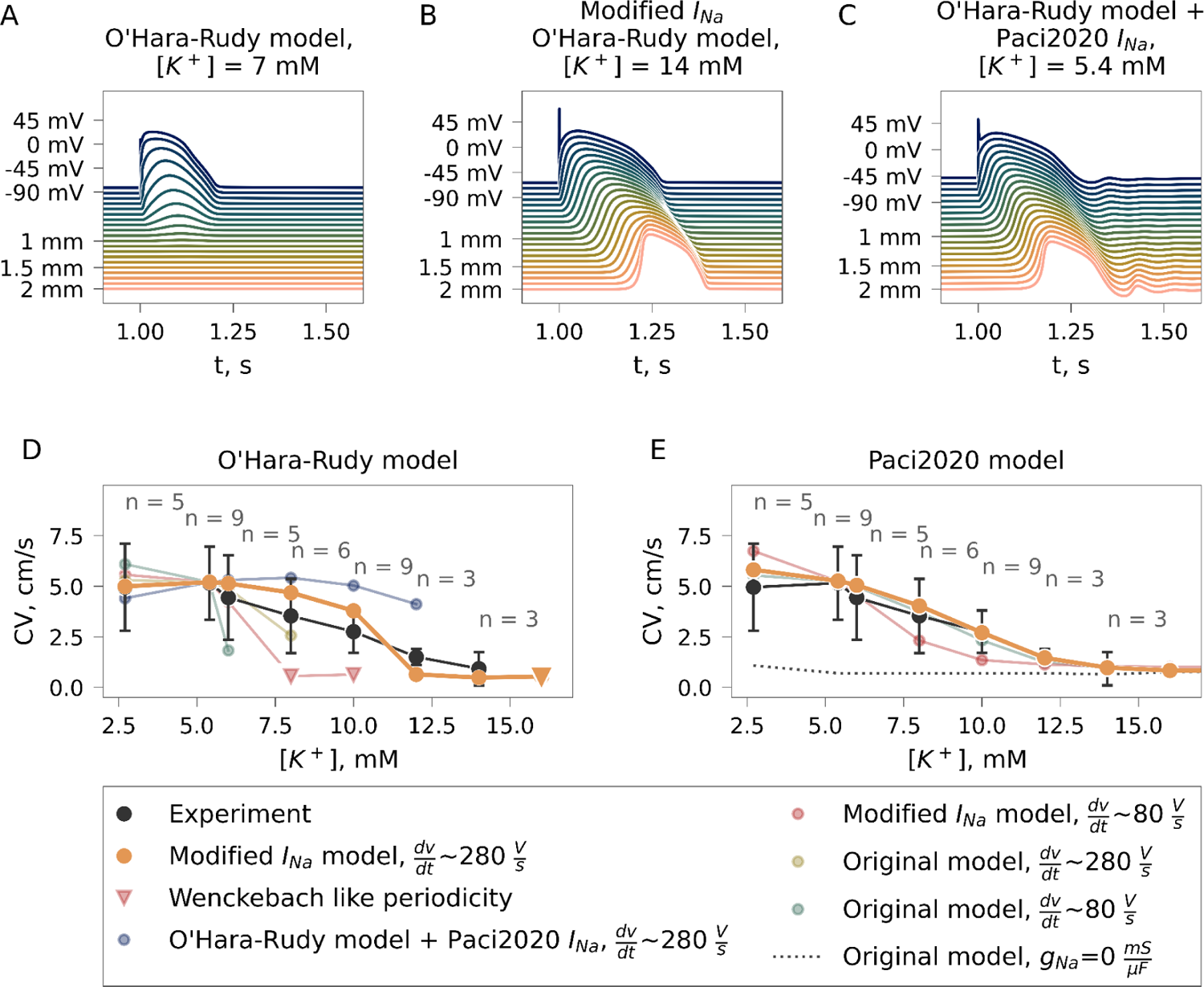
Action potential propagation in hyperkalemic conditions. **(A-C)** AP propagation in ventricular tissue simulated with three different I_Na_ formulations, – **(A)** O’Hara-Rudy (O’Hara *et al*., 2011) at [K^+^]_o_ = 7.0 mM; **(B)** this study at [K^+^]_o_ = 14.0 mM; **(C)** Paci2020 (Paci *et al*., 2020) I_Na_ model at [K^+^]_o_ = 5.4 mM. **(D, E)** Experimental CV dependence on [K^+^]_o_ compared to AP propagation simulated with **(D)** O’Hara-Rudy and **(E)** Paci2020 models. The black line corresponds to experimental data; orange – this study, *dV/dt=280 V/s;* red – this study, *dV/dt=80 V/s;* cyan – original model formulation (O’Hara-rudy in **(D)**, Paci2020 in **(E)**), *dV/dt=280 V/s;* green – original model formulation, *dV/dt=80 V/s*; violet – O’Hara-Rudy model combined with Paci2020 I_Na_ equations.

An increase of extracellular potassium by 1mM in the original O’Hara-Rudy model (O’Hara *et al*., 2011) completely abolishes AP propagation (Fig. 5A). The CV depends both on gap junction conductivity and sodium current amplitude (Tse & Yeo, 2015), therefore propagation failure in slightly hyperkalemic conditions may be caused by reduced I_Na_. Therefore, we have tested how increased gNa in the original O’Hara-Rudy model from 75 mS/μF to 210 mS/μF would affect the CV dependence. While the upstroke, dV/dt_MAX_, increased from 87 to 286 V/s in normokalemic conditions, the critical [K^+^]_o_ concentration changed only slightly, from 6 to 8 mM (Fig. 5D, green line).

Typical measurements of dV/dt_MAX_ in ventricular tissue are between 200 and 300 V/s (Drouin *et al*., 1995; O’Hara *et al*., 2011; Lemoine *et al*., 2017) (note, however, that actual upstroke velocity may be even higher (Windisch *et al*., 1995)). A much wider range is reported in iPSC-CM studies (Ma *et al*., 2011; Davis *et al*., 2012; Feaster *et al*., 2015; Herron *et al*., 2016; Lemoine *et al*., 2017): as low as 28+/-5 V/s (Ma *et al*., 2011) and as high as 219+/-15 V/s (Lemoine *et al*., 2017). Because of that, we performed two series of simulations with the O’Hara-Rudy model modified according to our measurements of I_Na_ voltage-dependence: with the sodium current maximal conductivity corresponding to higher dV/dt_MAX_ of 288 V/s and lower dV/dt_MAX_ of 82 V/s corresponding to the propagating AP in the original O’Hara-Rudy model. The case of large gNa resulted in CV dependence similar to experimental data (Fig. 5D, orange line). The CV decreased from 5.2 cm/s normokalemic conditions to 0.6 cm/s at 12 mM [K^+^]_o_, and, similarly to experimental data, the critical [K^+^]_o_ concentration was 14 mM (Fig. 5B). While it was still possible to pace the tissue at slightly higher potassium concentration (16 mM), we observed Wenckebach-like periodicity with 1 AP propagation to 8 stimuli (1:8). The simulations with lower gNa did not agree well with experimental data: CV decreased abruptly to 0.6 cm/s (Fig. 5D, red line) at 8mM [K^+^]_o_, with Wenckebach-like periodicity at the 8 and 10 mM [K^+^]_o_.

Stem-cell-derived myocytes exhibit spontaneous activity sustained by both hyperpolarization-activated current (I_f_) and Na^+^/Ca^2+^ exchanger (I_NCX_) (Kim *et al*., 2015; Koivumäki *et al*., 2018; Paci *et al*., 2020). The maximal diastolic potential is thus significantly depolarized by these currents in comparison to K^+^ Nernst potential, which, in turn, affects CV dependence on extracellular potassium concentration. In order to account for these factors, we have performed simulations of iPSC-CM tissue with the Paci2020 model (Paci *et al*., 2020). The original Paci2020 model paced in 1D-tissue resulted in *77 V/s dV/dt_MAX_,* which is similar to the O’Hara-Rudy model (*86 V/s*) and reproduced experimental CV dependence on [K^+^]_o_ (Fig. 5E). However a complete block of I_Na_ still resulted in propagating AP (dashed line in Fig. 5E) and, consequently, the tissue could be paced even at very high potassium concentrations (>*30 mM*).

The Paci2020 model with I_Na_ activation/inactivation modified according to our measurements in this study was not spontaneously active. The increase in I_f_ conductivity restored spontaneous activity, but for the sake of simplicity, we kept all parameters unrelated to I_Na_ unmodified in tissue simulations. Similarly to the O’Hara-Rudy model above, we have performed two series of simulations: we used g_Na_=77.4 mS/τF, corresponding to dV/dt_MAX_ of 270 V/s; and g_Na_=35.5 mS/τF, corresponding to dV/dt_MAX_ of 80 V/s. In the former case (Fig. 5E) CV dependence reproduced the experimental measurements. However, as mentioned above, contrary to experimental data AP propagated even at very high extracellular potassium. In the case of low g_Na_, we observed a somewhat more abrupt drop of CV in hyperkalemia: an increase of [K^+^]_o_ from *5.4* to *8 mM* resulted in the reduction of CV from 5.2 to *2.3 cm/s*. Both our and Paci2020 I_Na_ formulations thus seem to reproduce the experimental data, but our model suggests that the upstroke of the AP is more rapid when compared to measurements in iPSC-CMs. However, it should also be noted that the upstroke depends on I_f_ conductivity and the increase of funny current by a factor of 5, not only restored spontaneous activity but also reduced *dV/dt_MAX_*in stimulated tissue from 270 V/s to the value 100 V/s more typical to iPSC-CM measurements.

Paci2020 I_Na_ model is based on the earlier (Koivumäki *et al*., 2018) model, which, in turn, is based on (Ma *et al*., 2011) experiments. Although the half-activation measured by Ma et. al. is much less extreme than the estimate we propose in the current study, it still seems to agree with experimental CV dependence in hyperkalemic conditions. To further compare these two possible I_Na_ descriptions we hypothesized that the voltage dependence of the sodium channel voltage dependence in iPSC-CM is actually the same as in the ventricular tissue. We thus simulated the ventricular tissue with the O’Hara-Rudy model modified according to Paci2020 I_Na_ equations. Due to the fact that the I_Na_ formulation used in the Paci2020 model has a relatively large window current very close to the resting potential (Fig. S5), it was impossible to increase the depolarization rate above 150 V/s by a simple increase of I_Na_: it resulted in unphysiological oscillations and a resting potential above −50 mV, especially at hypokalemic conditions (Fig. 5C). Simultaneous increase of I_Na_ and I_K1_ conductivities abolished the oscillations, restored the resting potential of −87 mV and resulted in dV/dt_max_ above 200 V/s typical to ventricular tissue (Drouin *et al*., 1995; O’Hara *et al*., 2011; Lemoine *et al*., 2017). However due to the proximity of the window current to the resting potential we observed only a mild dependence of CV on [K^+^]_o_: at 12 mM concentration CV decreased to 4 cm/s (Fig. 5D, blue line). Furthermore, it was impossible to pace the tissue at 14 mM potassium concentration, since the resting potential at this concentration coincided with the maximal window current (∼ −60 mV).

## Discussion

Emerging practical applications of cardiac models both in drug-safety studies and patient-specific simulations (Dutta *et al*., 2017a; Boyle *et al*., 2019; Tomek *et al*., 2019; Fassina *et al*., 2023) require an accurate description of ionic channel dynamics. The fast kinetics and high amplitude of sodium current may result in uncontrollable membrane potential during patch-clamp recordings, a challenge that has been a persistent issue for the electrophysiology community for decades. Table 1 demonstrates that human sodium current activation and inactivation parameters in literature are quite contradicting. For example measurements of half activation range from −25 to −53 mV. These discrepancies can be explained by several factors.

Firstly, the sodium channel is assembled from the pore-forming alpha-subunit SCN5A and auxiliary beta-subunits, the latter interacting with the alpha-subunit and modulating its voltage dependence (Balser, 1999). Moreover, alternatively spliced variants of beta-subunits are known to affect voltage dependence (Martinez-Moreno *et al*., 2020). Inherited disorders including dilated cardiomyopathy are known to shift sodium current activation and inactivation (George, 2005; Nguyen *et al*., 2008). These facts imply that recordings from model objects, such as iPSC-CM and HEK cells, and diseased hearts may not correspond to native cardiac sodium channels and may account for some discrepancies in the published data. However, although the half-activation range in myocytes extracted from human ventricles is somewhat more narrow, half-activation still ranges from −38 to −52 mV (Table 1).

Secondly, both kinetics and steady-state voltage dependence of ionic channels is known to change with temperature (Nagatomo *et al*., 1998). While traditionally room temperature measurements were extrapolated to physiological temperature (Tusscher *et al*., 2004; O’Hara *et al*., 2011) this technique results in imprecise predictions: for example, Q10 estimates for sodium channel activation kinetics in literature are quite contradicting, the measurements range from 1.4 to 2.7 (Nagatomo *et al*., 1998; Wang *et al*., 2002). Moreover, there is no analog of Q10 for steady-state activation/inactivation and linear extrapolation to the physiological temperature is generally used (Tusscher *et al*., 2004; O’Hara *et al*., 2011).

Lastly, the discrepancy in patch-clamp recordings may be attributed to the experimental artifacts. This question was previously discussed in the study of HERG channels by Clerx et.al. (Lei *et al*., 2020): what is the source of variability in the patch-clamp recordings? Their conclusion was that accounting for cell-specific conductances and experimental artifacts with no actual difference in channel dynamics can explain the discrepancy. Our study confirms these conclusions.

As shown in Fig. 5 and Table 1 the apparent activation in our experiments was −52.9 ± 5.0 mV which corresponds to measurements in native ventricular myocytes (Valdivia *et al*., 2005; Barajas-Martínez *et al*., 2013). The inactivation of −70.6 ± 6.4 mV is somewhat depolarized in comparison with ventricular myocytes but agrees with stem-cell-derived myocytes data reported by other authors (Satin *et al*., 2004; Ma *et al*., 2011; Terrenoire *et al*., 2013). However, several features of our recordings suggested uncontrollable membrane potential. Firstly, there was a high discrepancy between our recordings: apparent half-activation ranged from −58.6 to −35.6 mV. Secondly, the activation curve was often very steep as exemplified in the Fig. S6: I_Na_ activated abruptly at some voltage step and the slope of the activation curve below 1 mV was common in our measurements.

Indeed, we were not able to replicate the recorded current traces with the model when I_Na_ was simulated with *apparent* activation/inactivation parameters. On the other hand, as we show in Fig. 4 optimization of a model accounting for the voltage difference between the amplifier and cell membrane allowed us to replicate the recording at every voltage step with high precision. The model parameters derived from optimization imply that even a minor sodium current activation results in membrane depolarization which explains the steepness of the activation curve in individual experiments. Apart from the high amplitude of sodium current, two factors are responsible for this undesirable artifact: low membrane capacitance of iPSC-CM (20 ± 10 pF) and high access resistance (120 ± 70 MOhm). While the latter is too high, the dependence of access resistance on the perforation time was previously measured (Kyrozis & Reichling, 1995) and ∼100 MOhm access resistance was reported after 20-30 minutes of antibiotic application, even longer perforation was required in order to achieve 10-20 MOhm. Thus access resistance might be improved by membrane rupture with prolonged perforation, electroporation, or negative pressure, however, these techniques resulted in an unstable seal at physiological temperature in our recordings, unstable seal at higher temperatures was previously reported, for example, in (Nagatomo *et al*., 1998; Wang *et al*., 2002). Similarly to our study, Windisch et. al (Windisch *et al*., 1995) reported that AP upstroke velocity could be underestimated by ∼30% in the current-clamp recordings. They demonstrate the crucial influence of cell access resistance in current-clamp setups, which sometimes exceed 50 MOhm and may unpredictably increase during experiments despite low pipette resistance. We may assume that these changes in access resistance may actually stand behind the time-dependent hyperpolarizing shifts in sodium current activation that was shown previously (Hanck & Sheets, 1992).

The nature of the artifact discussed above results in the hyperpolarizing shift of the measured *apparent* activation curve and depolarizing shift of the inactivation curve; the difference between the apparent activation/inactivation curve and the result of model optimization was very large: 30 mV/-19 mV correspondingly. The error is systematic, thus increasing the number of experiments would not eliminate the shift. The nature of this systematic error in the apparent activation curve points out a problem in the patch-clamp measurements in general: while this error can be reduced, it still plagues the experimental data. For instance, that implies that *apparent* sodium current activation measured as the recorded current amplitude dependence on the voltage step is always hyperpolarized to some degree. This might explain why our estimate of the half-activation, which is −10 mV (95% CI is [−16.5 mV, −3.0 mV]), is depolarized in comparison with the literature quoted in Table 1.

As discussed above, voltage-dependence of ionic current in iPSC-CM might differ from native myocytes, however, abnormalities of mathematical models and, in particular, their behavior in hyperkalemic conditions, imply errors in sodium current measurements in native myocytes as well. Increased extracellular potassium concentration depolarizes the resting potential partially inactivating the sodium current. This, in turn, decreases the conduction velocity or, at higher concentrations, results in tissue inexcitability (Weiss *et al*., 2017). Classical mathematical models of ventricular tissue demonstrate propagation failure even at a mild increase of extracellular potassium concentration: in the case of the O’Hara-Rudy model (O’Hara *et al*., 2011) propagation failure occurs at extracellular potassium concentration above 6 mM (Fig. 3); in case of Grandi-Pasqualini-Bers model (Grandi *et al*., 2010), which uses sodium current based on formulation from earlier TenTusscher-Panfilov model (Tusscher *et al*., 2004) this happens at a concentration above 8 mM (Dutta *et al*., 2017b). More recently several research groups (Dutta *et al*., 2017b; Tomek *et al*., 2019; Bartolucci *et al*., 2020) introduced changes to I_K1_ and I_Na_ formulations in order to circumvent this limitation, however, these modifications were not quantitatively verified by experimental measurements in hyperkalemic conditions.

Thus, in order to validate our measurements of sodium current-voltage dependence, we have optically mapped monolayers of IPSC-CM. As we demonstrate above in Fig. 5, CV measurements provide evidence in favor of our patch-clamp data analysis: both CV changes and critical [K^+^]_o_ are in agreement with the simulation results of the O’Hara-Rudy model with modified sodium current. The functional immaturity of iPSC-CMs is known to result in a depolarized resting potential in comparison to native cardiomyocytes (Ma *et al*., 2011; Davis *et al*., 2012; Prè *et al*., 2014; Chen *et al*., 2017; Goversen *et al*., 2018) (however, also note a study by Horváth et. al. discussing that, in part, the technical issues may underlie reduced resting potential (Horváth *et al*., 2018)). Simulations with Paci2020 (Paci *et al*., 2020) model accounting for this immaturity were inconclusive (Fig. 5E): firstly, even severe hyperkalemia did not result in propagation failure; secondly, both original Paci2020 equations and the model proposed in the current study seem to reproduce CV dependence on [K^+^]_o_. The I_Na_ equations in the Paci2020 (Paci *et al*., 2020) and the earlier Koivumaki2018 (Koivumäki *et al*., 2018) iPSC models are based on measurements by Ma et. al. (Ma *et al*., 2011) (see Table 1). We have tested if voltage-dependence reported by Ma. *et.al.* can explain propagation in ventricular tissue (Fig. 5C,D, blue line). These measurements imply the prominent sodium window current ranging from −80 to −40 mV, close to the ventricular resting potential (Fig. S5). Tetrodotoxin-sensitive window current was reported in canine purkinje fibers as early as 1979 (Attwell *et al*., 1979), but the window range from −60 to −10 mV is reported in this study, which corresponds to our model (Fig. S5). The proximity between resting potential and sodium window current results in partial depolarization and oscillations of membrane potential that are due to I_Na_ activation (Fig. 5C). Moreover, the CV reduction in hyperkalemic conditions is less prominent than expected in ventricles (Fig. 5D, blue) (Weiss *et al*., 2017; King *et al*., 2021). As discussed above, innate immaturity of iPSC-CMs may imply that the sodium channel is different from the mature ventricular phenotype. However, the differences reported in previous studies can actually be explained by technical limitations of patch-clamp measurements, while it is possible to describe propagation both in iPSC-CM tissue and ventricular tissue using the same I_Na_ voltage-dependence.

In conclusion, in this study, we have measured human sodium current-voltage dependence at the physiological temperature. We have shown that the optimization of the voltage-clamp experiment model makes it possible to circumvent the common problem of uncontrollable membrane voltage in the patch-clamp experiment. Although our estimate of −10 mV half-activation is surprisingly depolarized, we demonstrate that these measurements are in agreement with action potential propagation in hyperkalemic conditions. However, experiments on native ventricular myocytes are required in order to refine our measurements of the fast sodium current-voltage dependence. Another limitation of our study is the fact that time constants were unidentifiable from standard voltage-step protocols, thus assessing sodium channel kinetics at physiological temperature requires optimized voltage-clamp protocols (Groenendaal *et al*., 2015; Lei *et al*., 2017, 2022).

## Abbreviations

CV: conduction velocity

AP: action potential

iPSC-CM: induced pluripotent stem-cell-derived cardiomyocytes

APD_80_: Action potential duration

MLE: Maximum likelihood estimator

CI: 95% credible intervals

## Additional information section

Data availability statement: Raw patch-clamp recordings are available at https://zenodo.org/record/7997728. Source code is available at https://github.com/humanphysiologylab/PCoptim.

Competing interests: Authors declare no competing interests.

## Author contributions

Conceptualization: RS, VT, SF Software: VA, RS

Investigation: VA, SR, SF, MS, VT, RS, SS, AA Visualization: VA

Supervision: RS

Resources: SR, SF, MS, VT, RS Writing—original draft: VA, RS Writing—review & editing: VA, SR, MS, VT, RS

## Supplementary Materials for

### Supplementary text

#### Model equations

##### Voltage-clamp experiment model

In this study we mostly follow the set of differential equations developed in the study by Lei et.al. (Lei *et al*., 2020). The only difference is the compensation: feedback current (Strickholm, 1995a) was used in the Lei et.al. study, but supercharging (Strickholm, 1995b) was used instead in our experiments. Supercharging technique decreases cell membrane charging time by increasing the actual voltage step, *V_cp_*, above the command potential, *V_c_*:

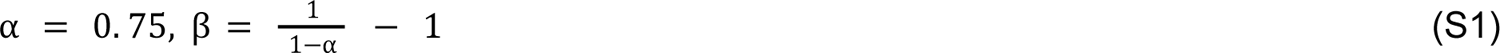

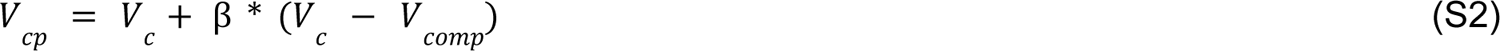

The *V* is gradually reduced with the time constant, τ, which is manually set by changing compensation resistance and capacitance during the experiment (*R* and *C* parameters):

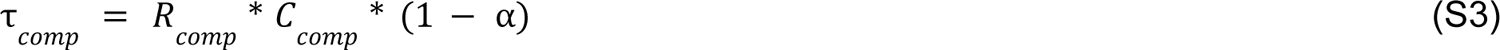

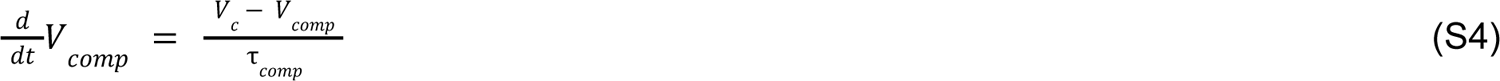

Cell membrane charging transient is removed from the recording by transferring the current, *I*, to the capacitance set in parallel with the feedback resistor, *C* (Axon Instruments Inc, 1997–1999):

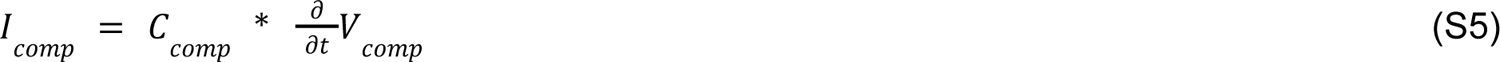

The membrane potential, *V*_m_, is determined by the sum of the current injected from the amplifier, sodium current, *I _Na_*, and the leak current,

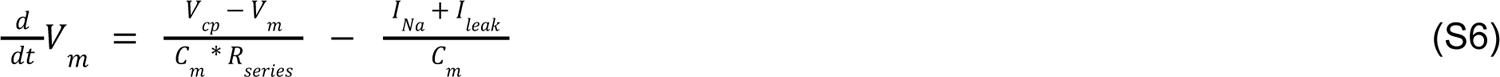

The input current, *I_leak_*, is contaminated by cell membrane capacitive current, *I _in_*, leak current, *I* _m_, while the membrane charging transient is partially compensated by

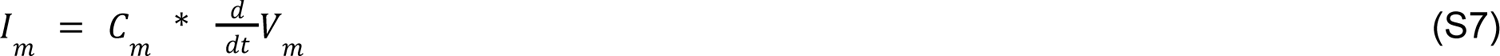

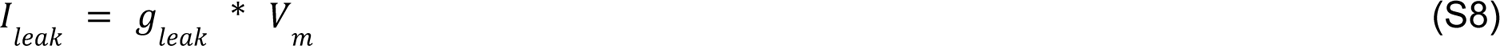

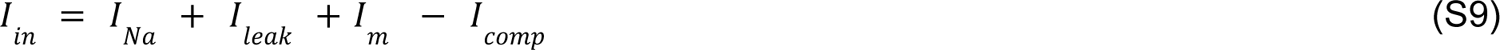

Finally the recorded current is the input current low-pass filtered through the feedback resistance, *R_f_*, set in parallel with the capacitance, *C_f_*:

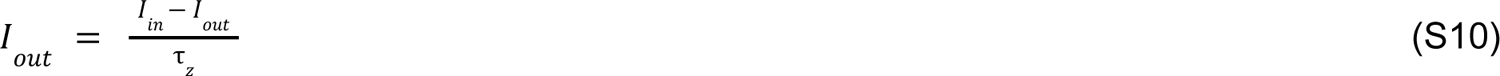

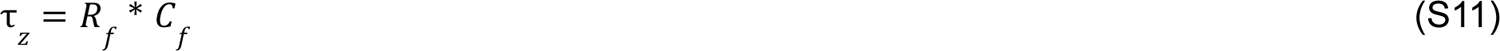

## Sodium current model

Sodium current is described via Hodgkin-Huxley formalism with three identical activating and two inactivating gates. In this study we used the equations from the O’Hara-Rudy model (O’Hara *et al*., 2011):

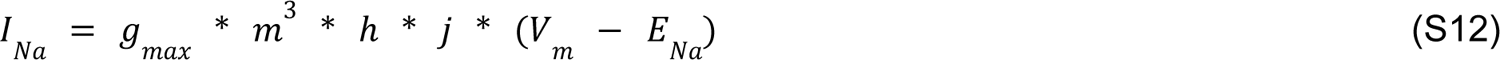

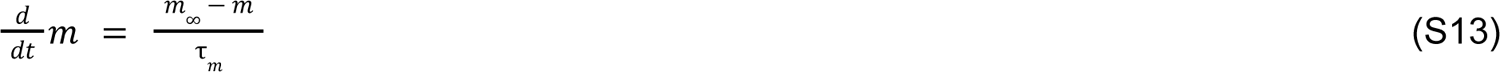

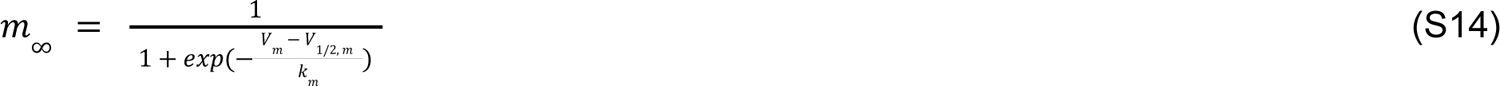

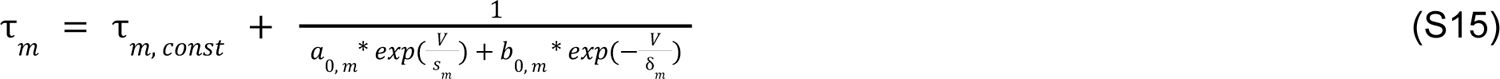

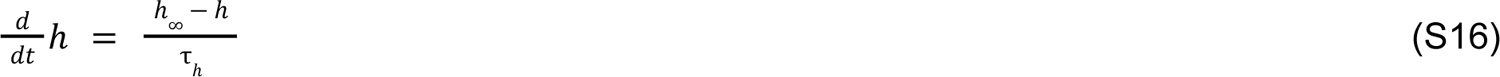

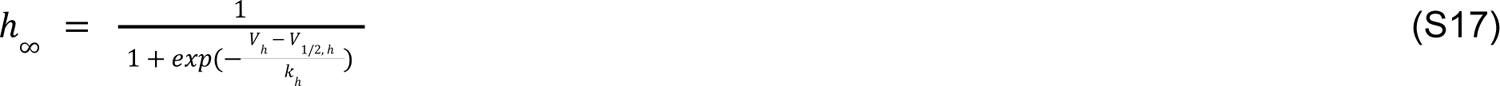

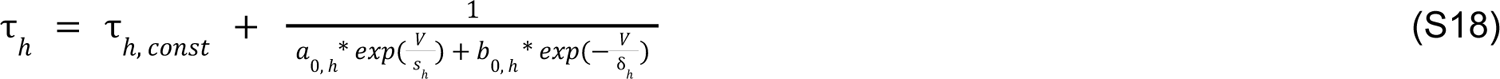

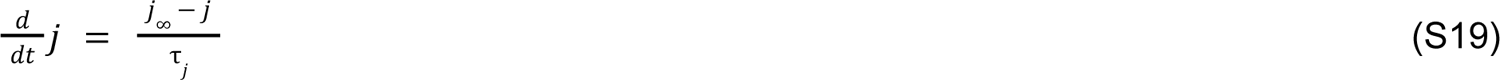

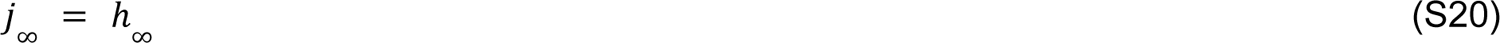

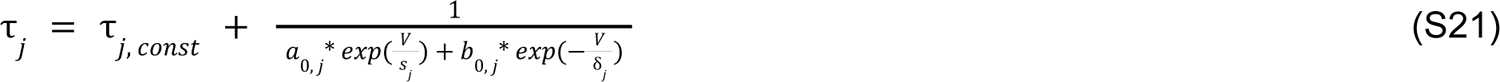

---

**Table S1.** Genetic algorithm free parameters, their bounds and values used for synthetic trace in Figs S2-S5 below.

| Parameter | Upper bound | Lower bound | Synthetic trace | Parameter | Upper bound | Lower bound | Synthetic trace |
| --- | --- | --- | --- | --- | --- | --- | --- |
| $a_{0,m}$ | 1652.906 | 165290.6 | 16529.06 | $\tau_{j,const}, ms$ | 0.132 | 13.2 | 4.6336 |
| $b_{0,m}$ | 38.67 | 3867 | 531.66 | $C_m, pF$ | 2.47 | 124.0 | 23.6 |
| $\delta_m, mV$ | 2.2823 | 228.23 | 42.695 | $R_{series}, M\Omega$ | 2.495 | 2494.68 | 105.025 |
| $s_m, mV$ | 1.28321 | 128.321 | 8.18 | $g_{Na}, nS$ | 419.62 | 839240 | 4416.57 |
| $\tau_{m,const}, ms$ | 1.87E-3 | 7.48E-2 | 3.337E-2 | $g_{leak}, nS$ | 0.04812 | 4.81244 | 1.13424 |
| $a_{0,h}$ | 0.626 | 62.6 | 3.4276 | $V_{1/2,h}, mV$ | -30 | -100 | 29.001 |
| $b_{0,h}$ | 5813.1407 | 581314.07 | 16889.72 | $V_{1/2,m}, mV$ | -10 | -60 | 77.0228 |
| $\delta_h, mV$ | 1.027583 | 102.7583 | 2.459711 | $k_h, mV$ | 6 | 15 | 9.00 |
| $s_h, mV$ | 2.162 | 216.2 | 21.62 | $k_m, mV$ | 6 | 15 | 11.285 |
| $\tau_{h,const}, ms$ | 0.02 | 2 | 0.02 | $C_{comp}, pF$ | 0.1 | 100. | 1.0 |
| $a_{0,j}$ | 0.06462 | 6.462 | 2.571 | $R_{comp}, M\Omega$ | 10 | 1000 | 40.829 |
| $b_{0,j}$ | 1135.831 | 113583.1 | 7581.26 | $\alpha$ | 0.72 | 0.78 | 0.7352 |
| $\delta_j, mV$ | 0.7445 | 74.45 | 5.66088 | $E_{Na}, mV$ | 16 | 80 | 34.9186 |
| $s_j, mV$ | 6.9417 | 694.17 | 69.417 | $\tau_z, ms$ | 0.04 | 0.600204 | 0.17222 |

**Table S2.** Bounds and prior distributions used for Markov chain Monte-Carlo sampling.

|  | Lower bound<br>(LB) | Upper bound<br>(UB) | Prior normal distribution<br>for activation trace |  | Prior normal distribution<br>for inactivation trace |  |
| --- | --- | --- | --- | --- | --- | --- |
| | | | $\mu$ | $\sigma$ | $\mu$ | $\sigma$ |
| $V_{1/2,m}$ , mV | -10 | -60 | -25 | 70 | -25 | 3 |
| $k_m$ , mV | 3 | 20 | 11.5 | 17 | 10 | 2.5 |
| $V_{1/2,h}$ , mV | -30 | -100 | -76 | 9 | -65 | 70 |
| $k_h$ , mV | 3 | 20 | 11 | 3 | 11.5 | 17 |
| $E_{Na}$ , mV | 1 | 150 | 18 | 7 | 18 | 7 |
| Remaining<br>parameters | See Table<br>S1 | See Table<br>S1 | (UB+LB)/2 | UB-LB | (UB+LB)/2 | UB-LB |

**Fig S1.**
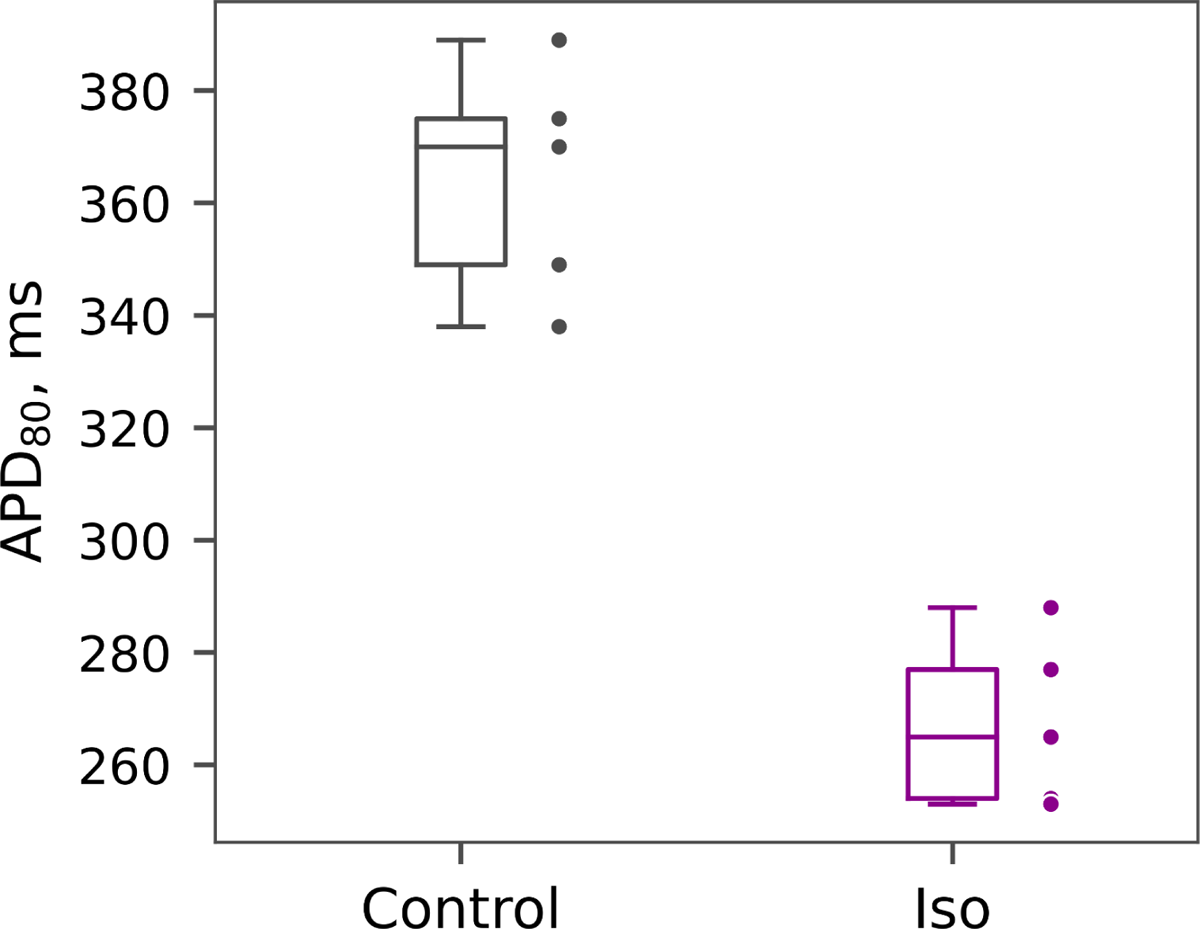
Beta-adrenergic effect on the action potential duration across different monolayers (n=5). 1 μm isoprenaline reduced APD80 from 364 ± 21 to 267 ± 15 ms.

**Fig S2.**
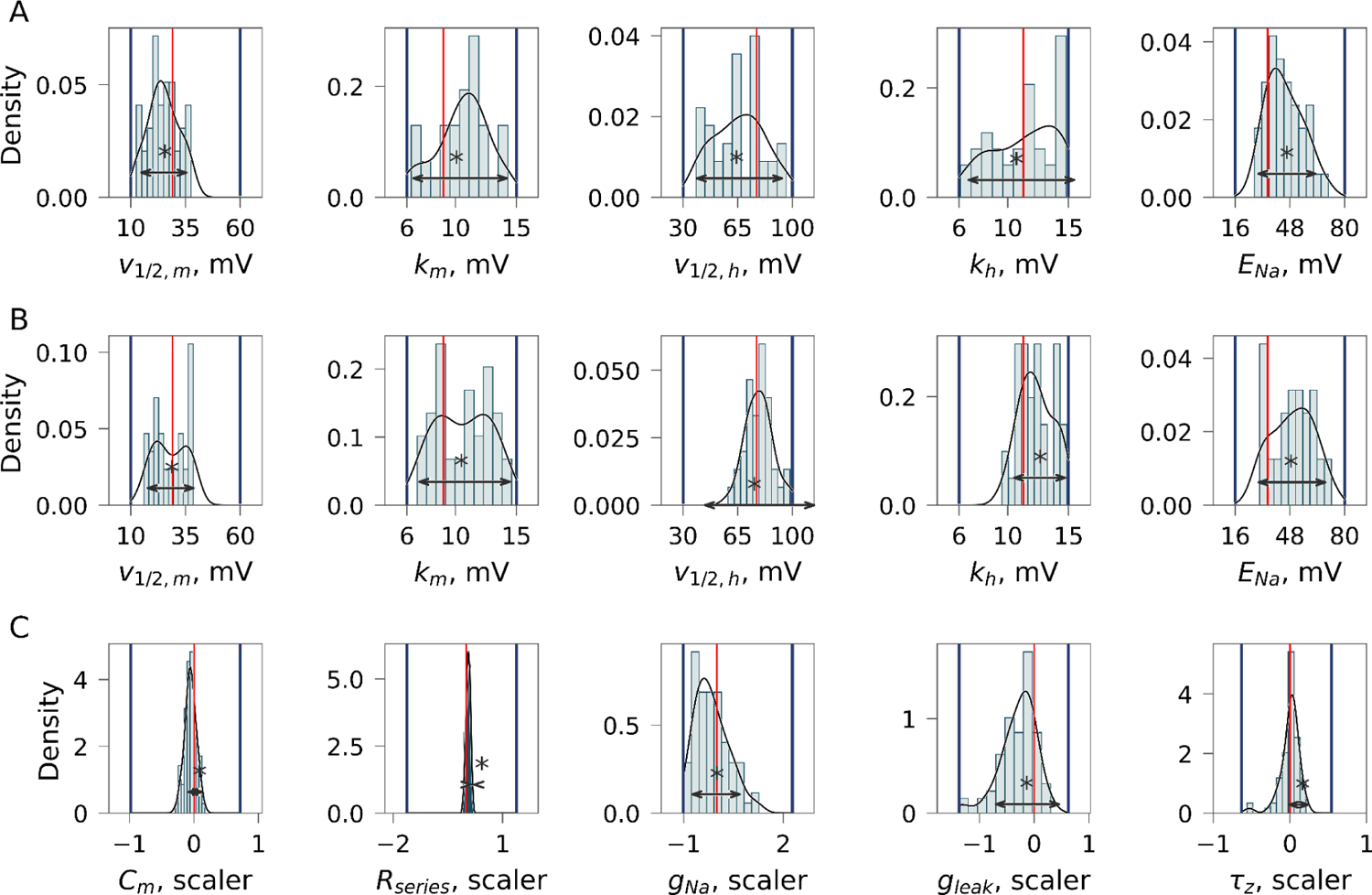
Histogram of parameters identified by genetic algorithm (GA) from synthetic activation trace(panel A, n = 39), inactivation trace(panel B, n = 38) and both of them combined (panel C, n =77): blue lines show GA parameter bounds, red line indicate input model values, 95%CI is shown by arrows below asterisk.

**Fig S3.**
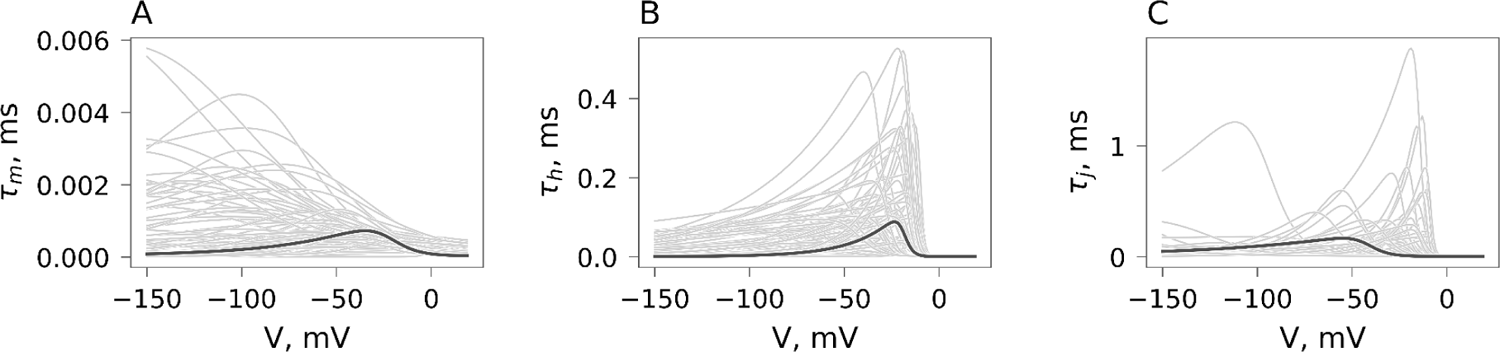
I_Na_ kinetics identifiability. Activation (A), fast inactivation (B) and slow inactivation (C) times identified by genetic algorithm (gray) compared to the input model (black).

**Fig S4.**
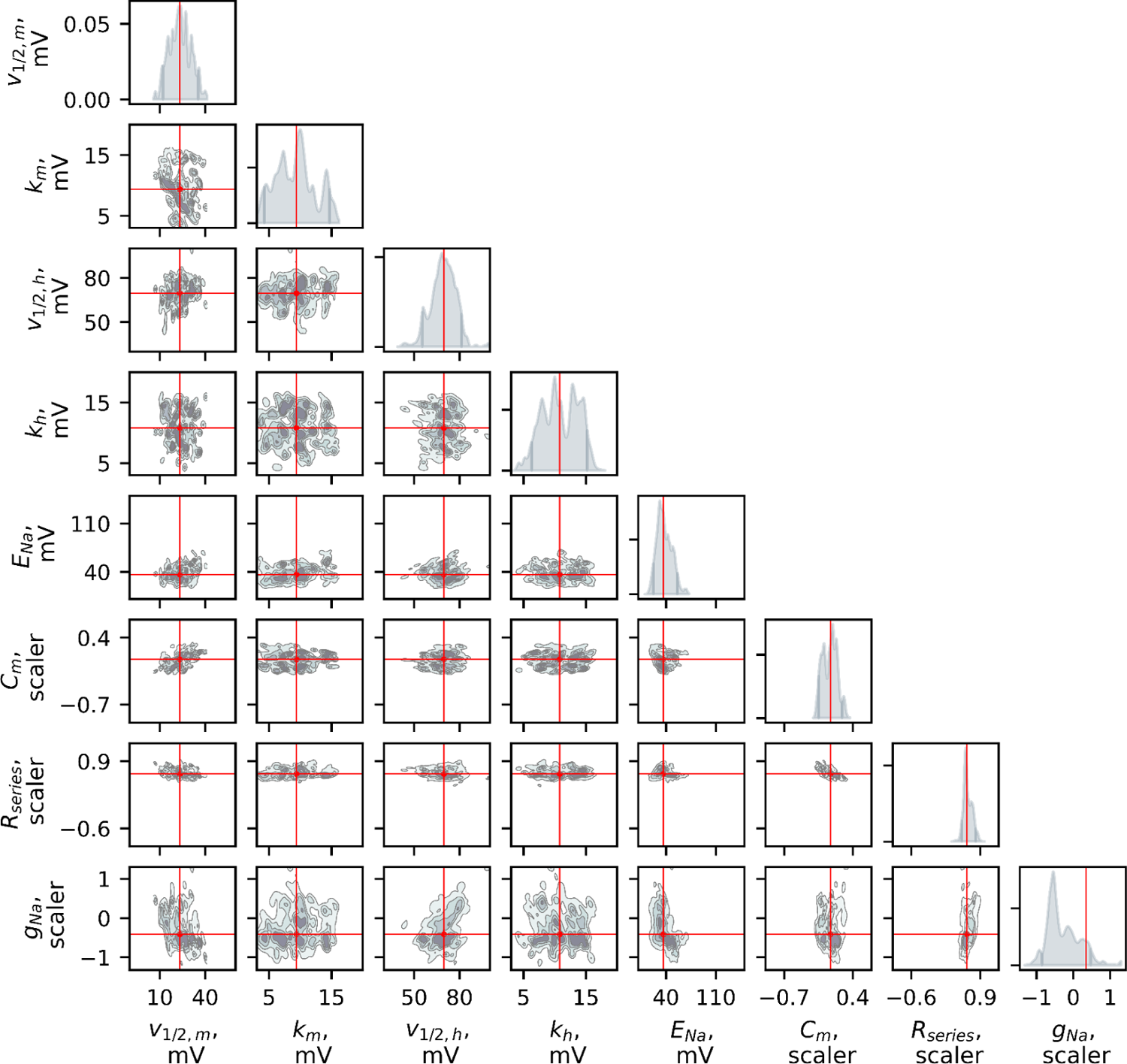
Example of Markov chain Monte-Carlo sampling of the posterior distributions. Model parameters were sampled for the activation trace shown in Fig S9.

**Fig S5.**
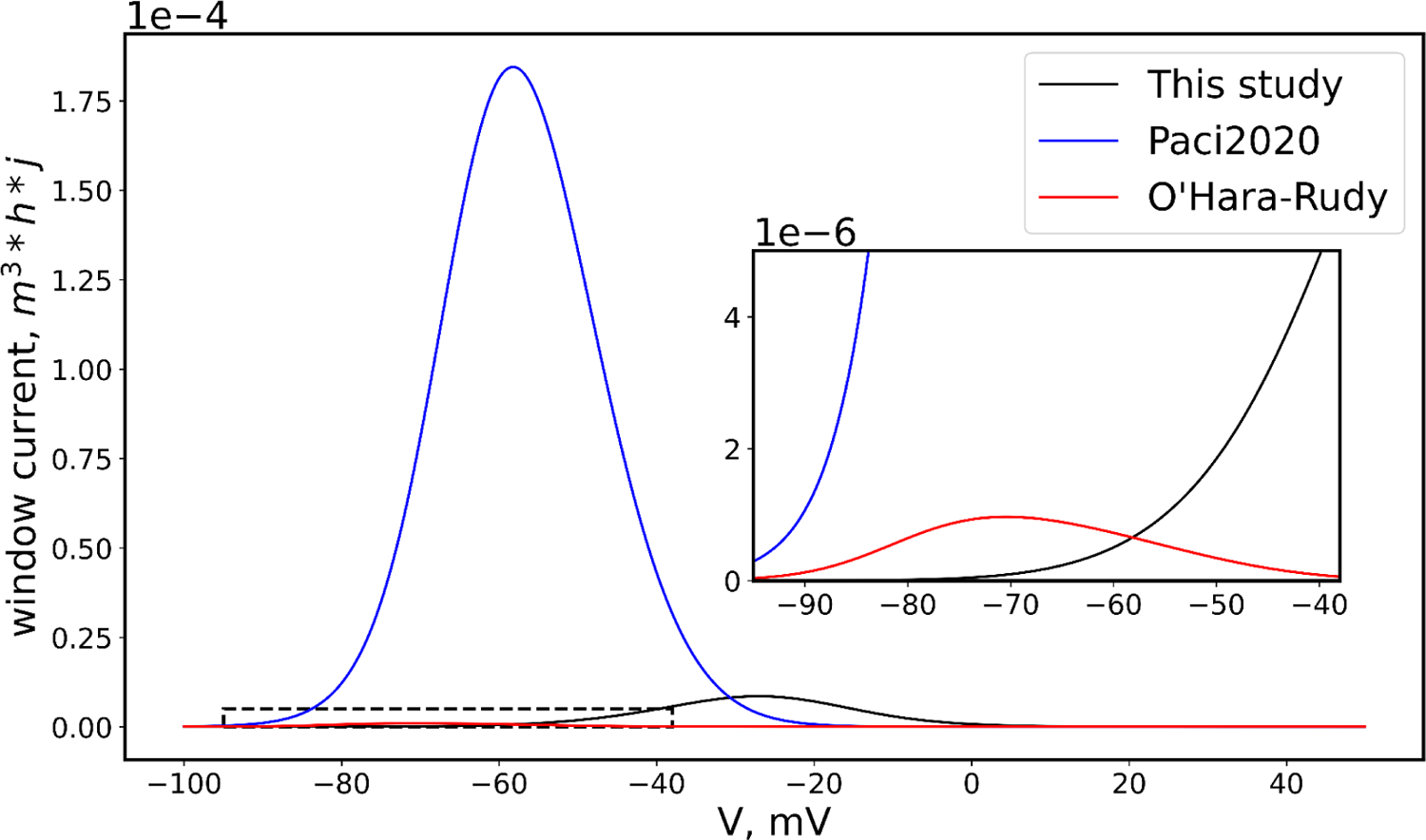
The comparison of I_Na_ window current for the model developed in this study, O’Hara-Rudy model (O’Hara *et al*., 2011) and Paci2020 model (Paci *et al*., 2020). The range marked by the dashed square is shown in inset.

**Fig S6.**
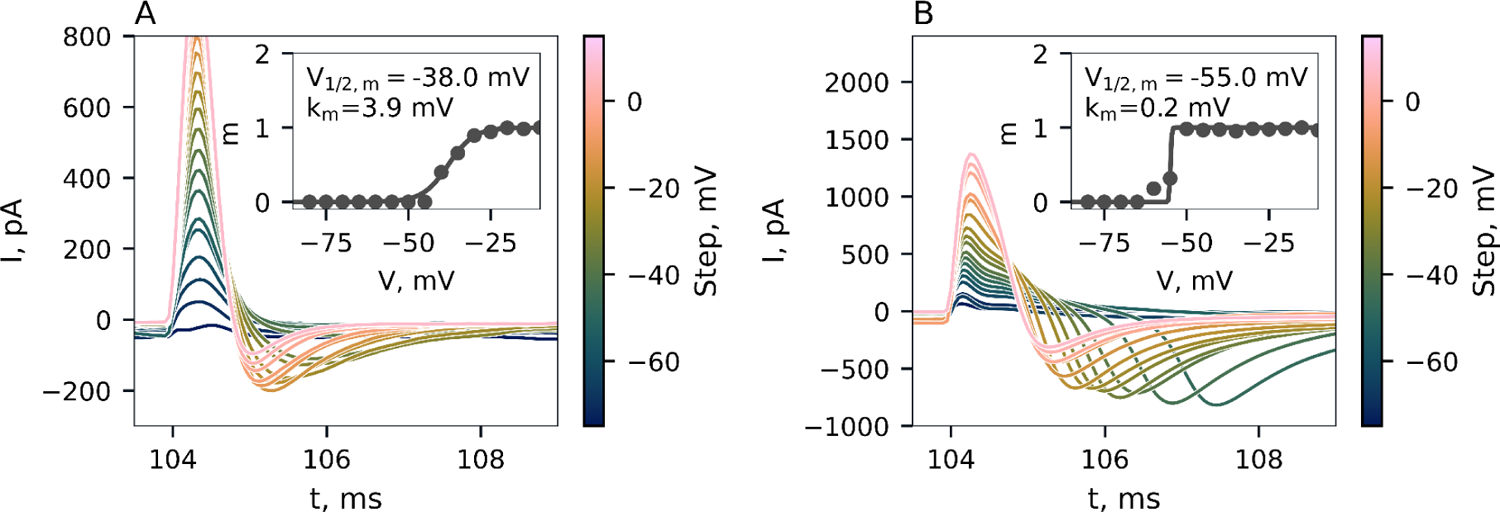
Representative examples of experimental activation curve measurements. Voltage step is shown by the color of the corresponding trace. Least square curve fit of the *apparent* activation curve is shown in insets. Note the steep slope (low k_m_) of the curve.

**Fig S7.**
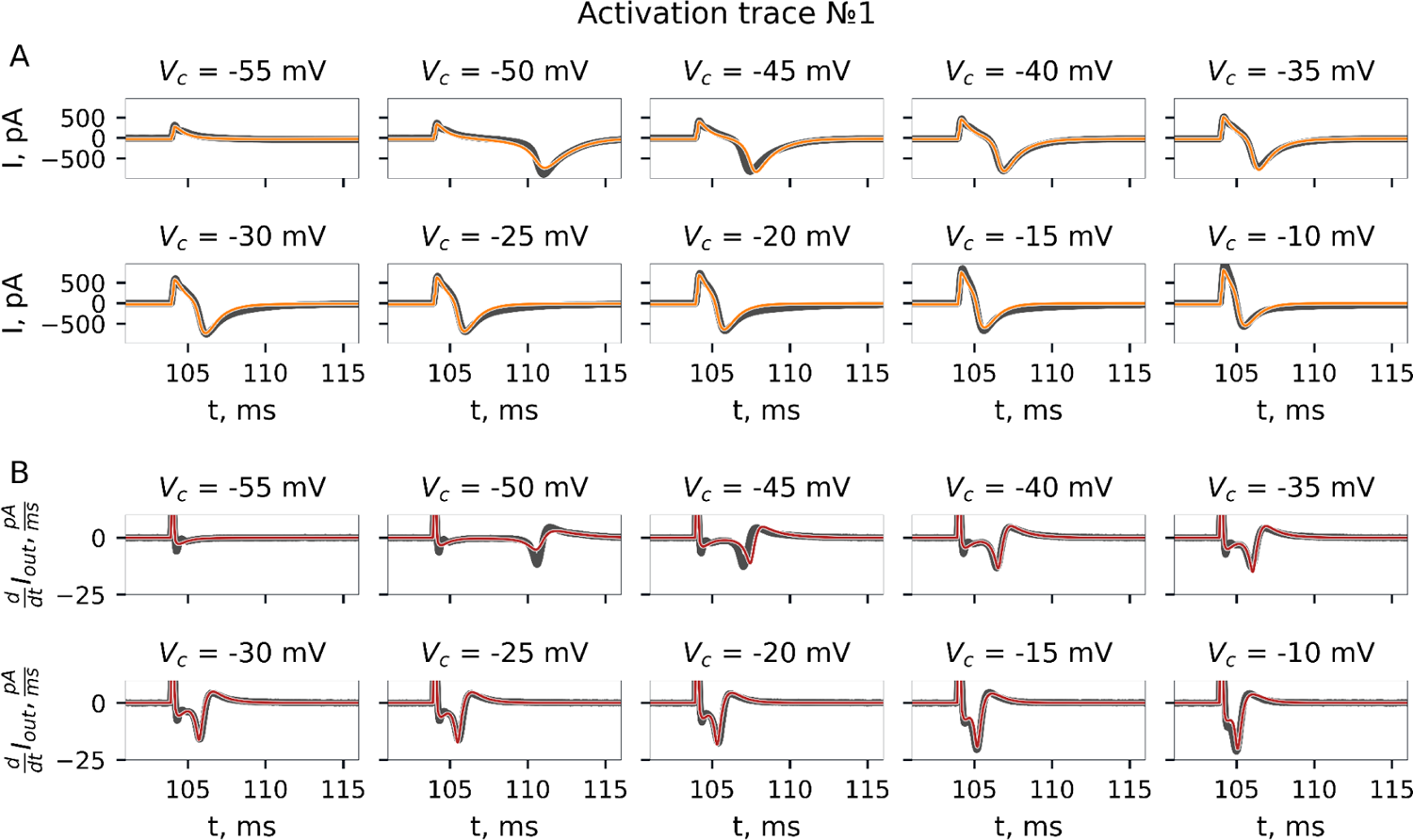
Activation protocol. Recorded current trace and its fit by GA. Experiment №1.

**Fig S8.**
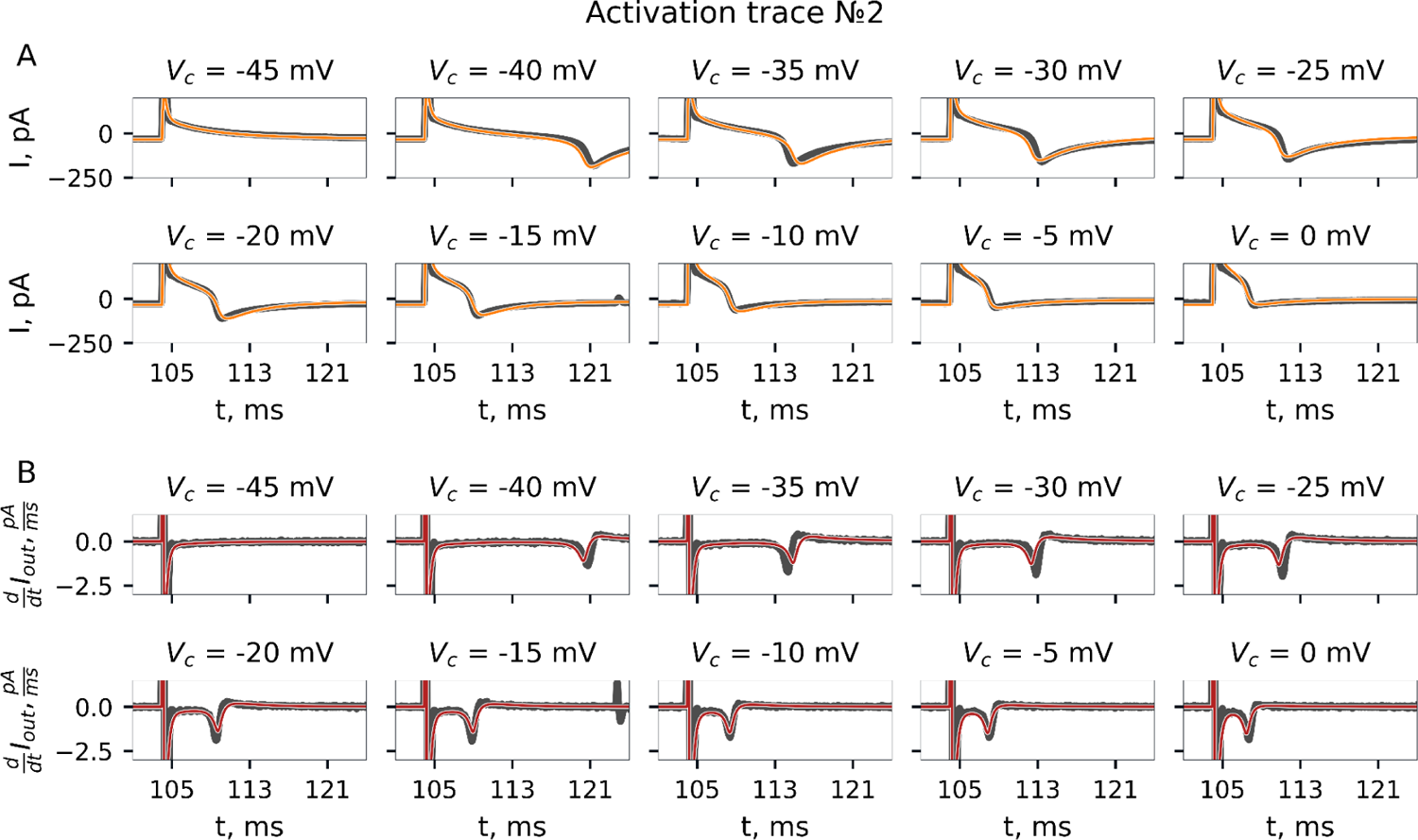
Activation protocol. Recorded current trace and its fit by GA. Experiment №2.

**Fig S9.**
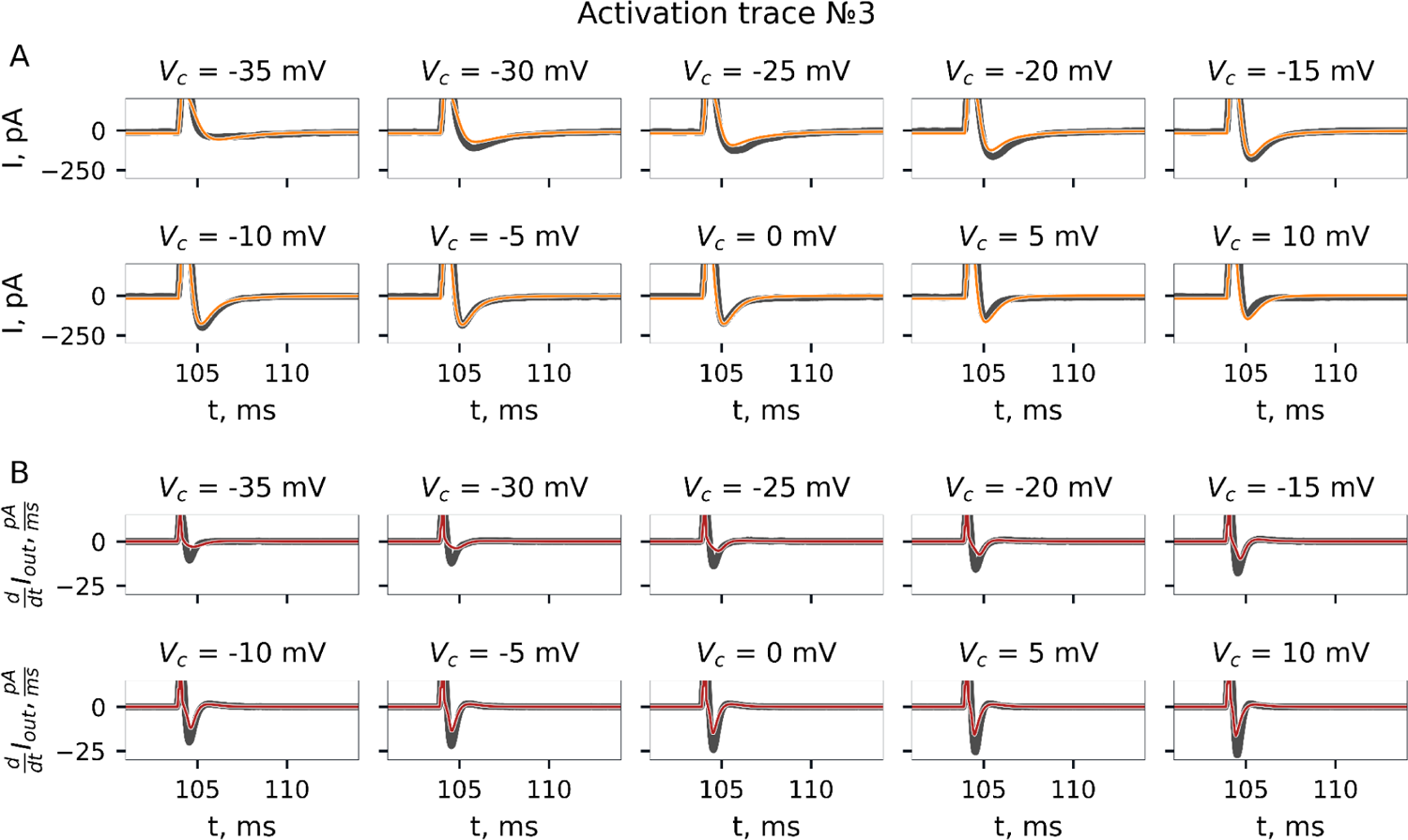
Activation protocol. Recorded current trace and its fit by GA. Experiment №3.

**Fig S10.**
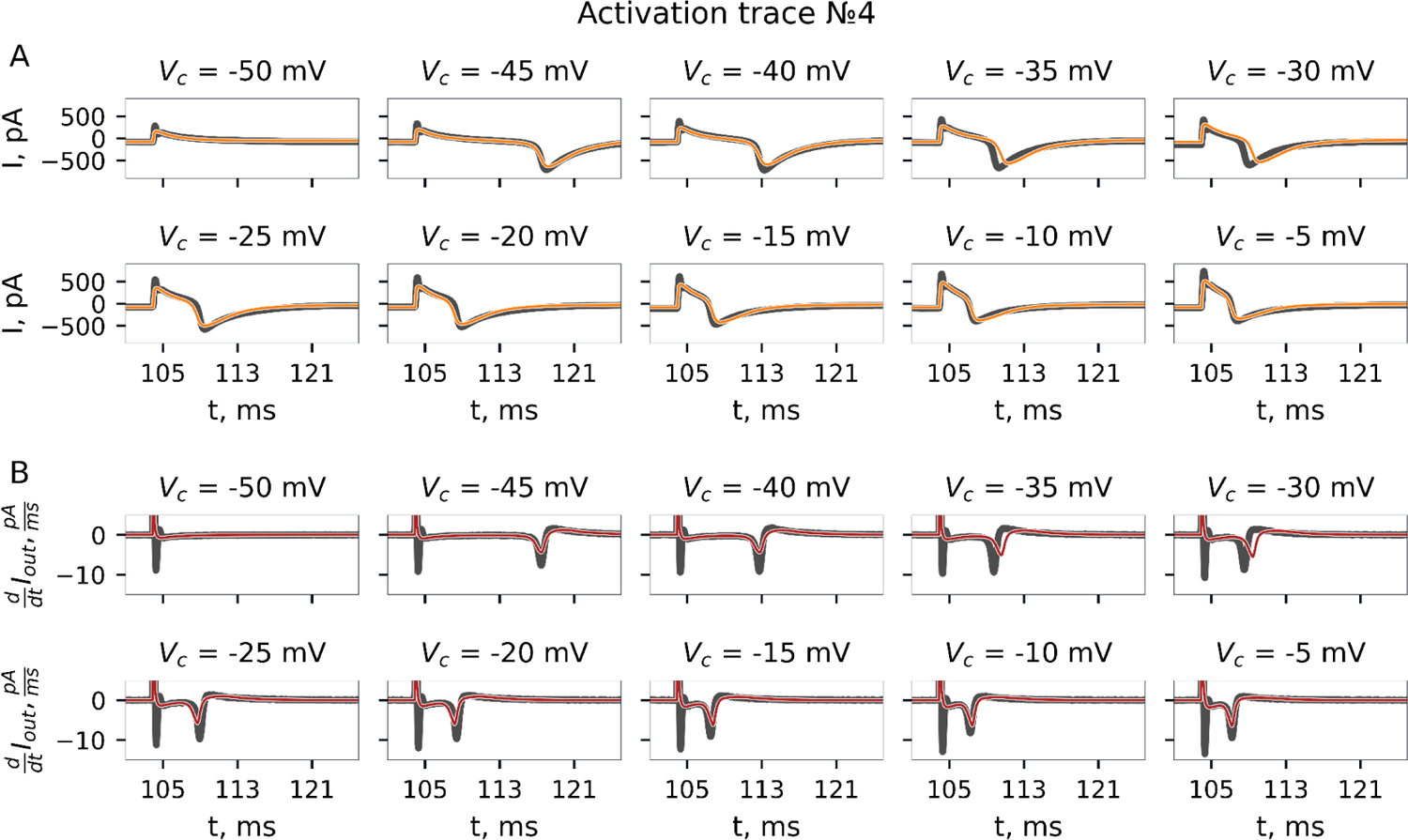
Activation protocol. Recorded current trace and its fit by GA. Experiment №4.

**Fig S11.**
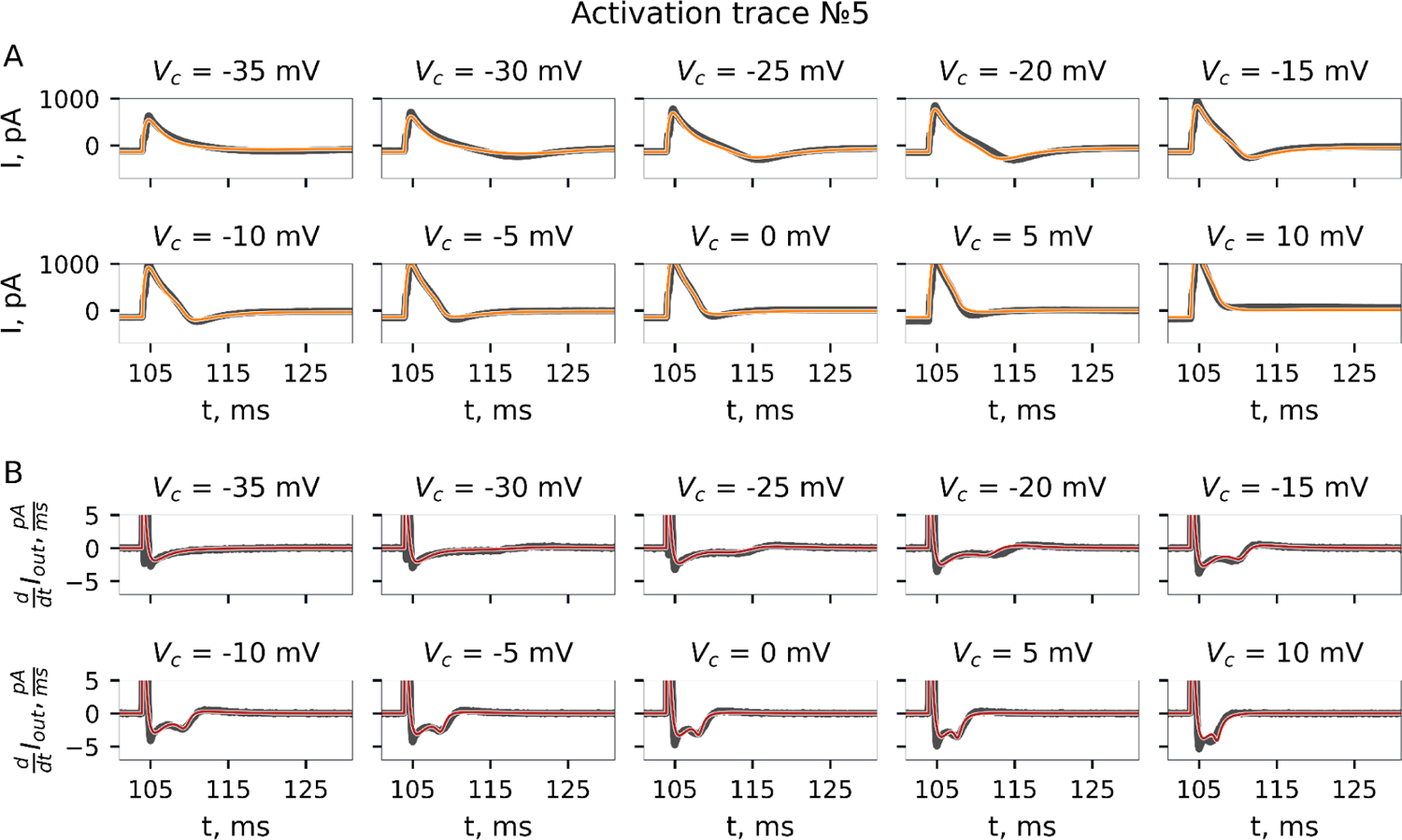
Activation protocol. Recorded current trace and its fit by GA. Experiment №5.

**Fig S12.**
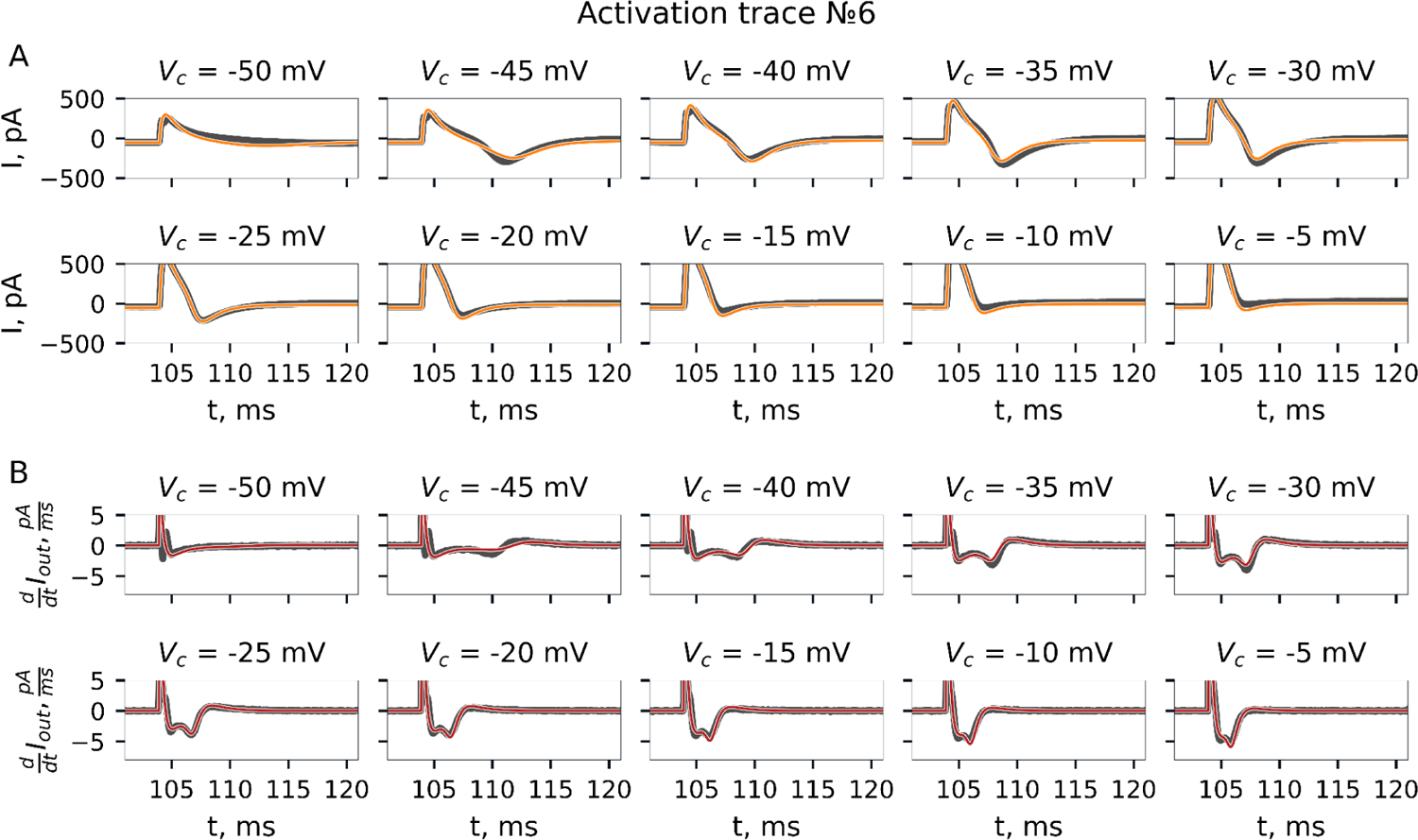
Activation protocol. Recorded current trace and its fit by GA. Experiment №6.

**Fig S13.**
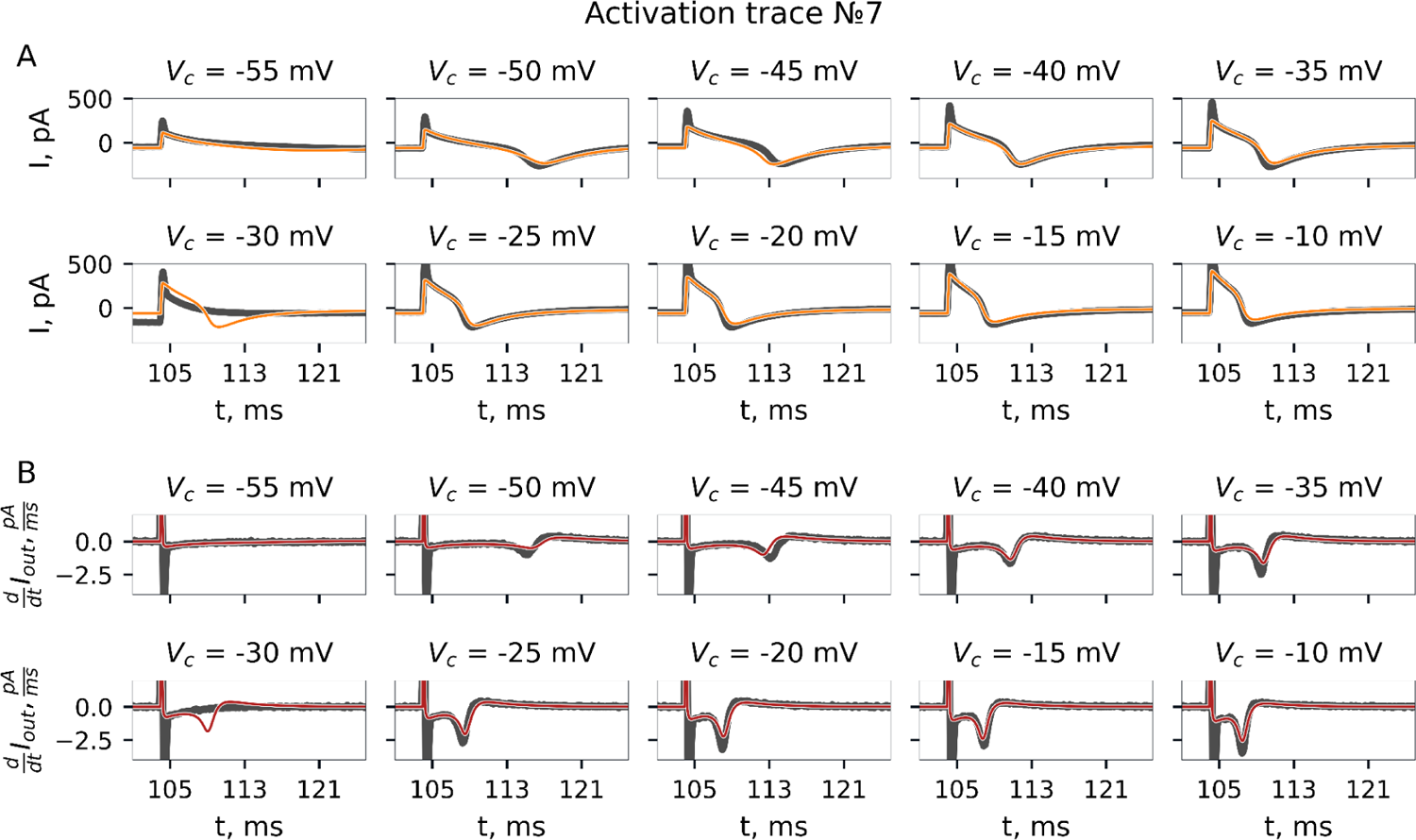
Activation protocol. Recorded current trace and its fit by GA. Experiment №7.

**Fig S14.**
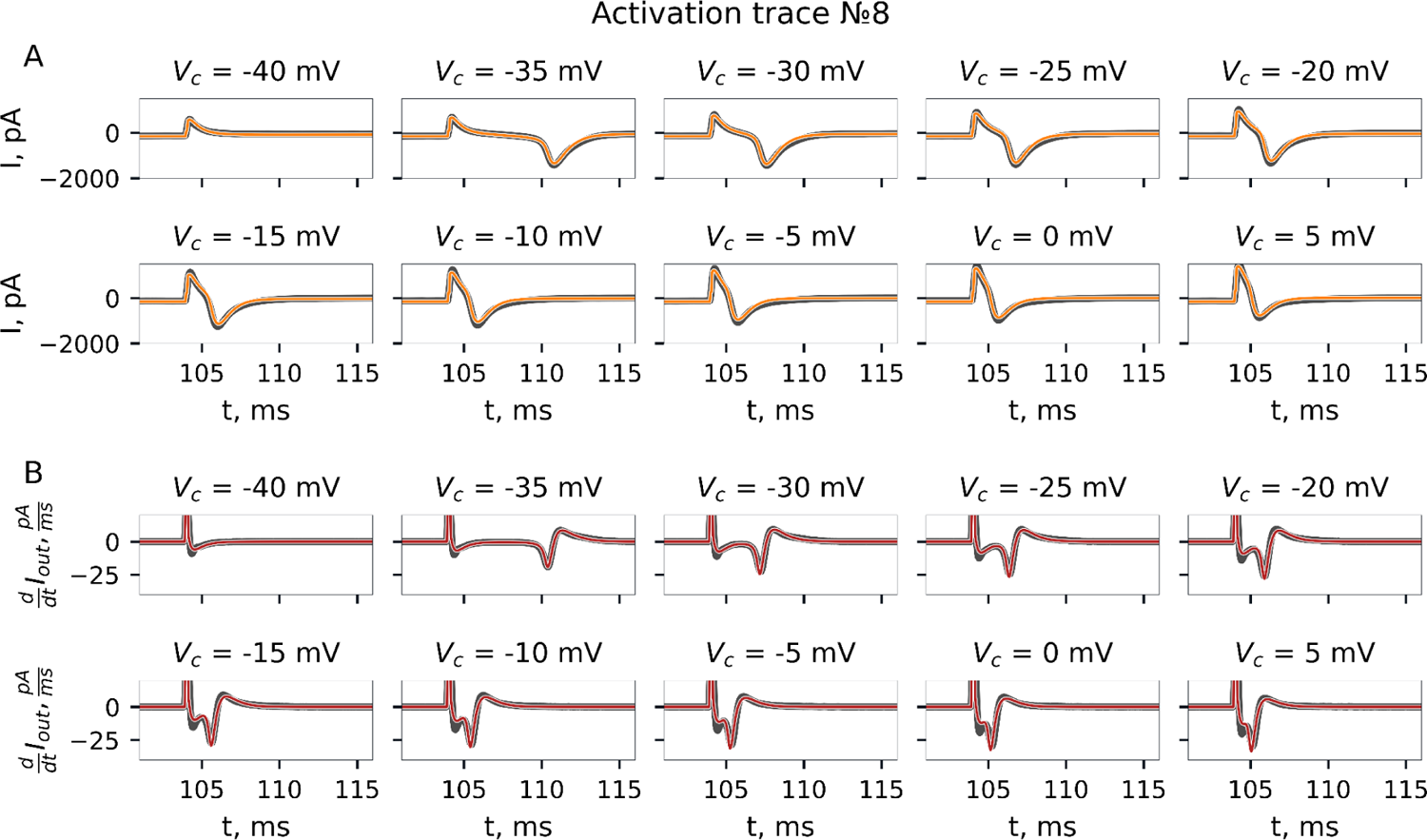
Activation protocol. Recorded current trace and its fit by GA. Experiment №8.

**Fig S15.**
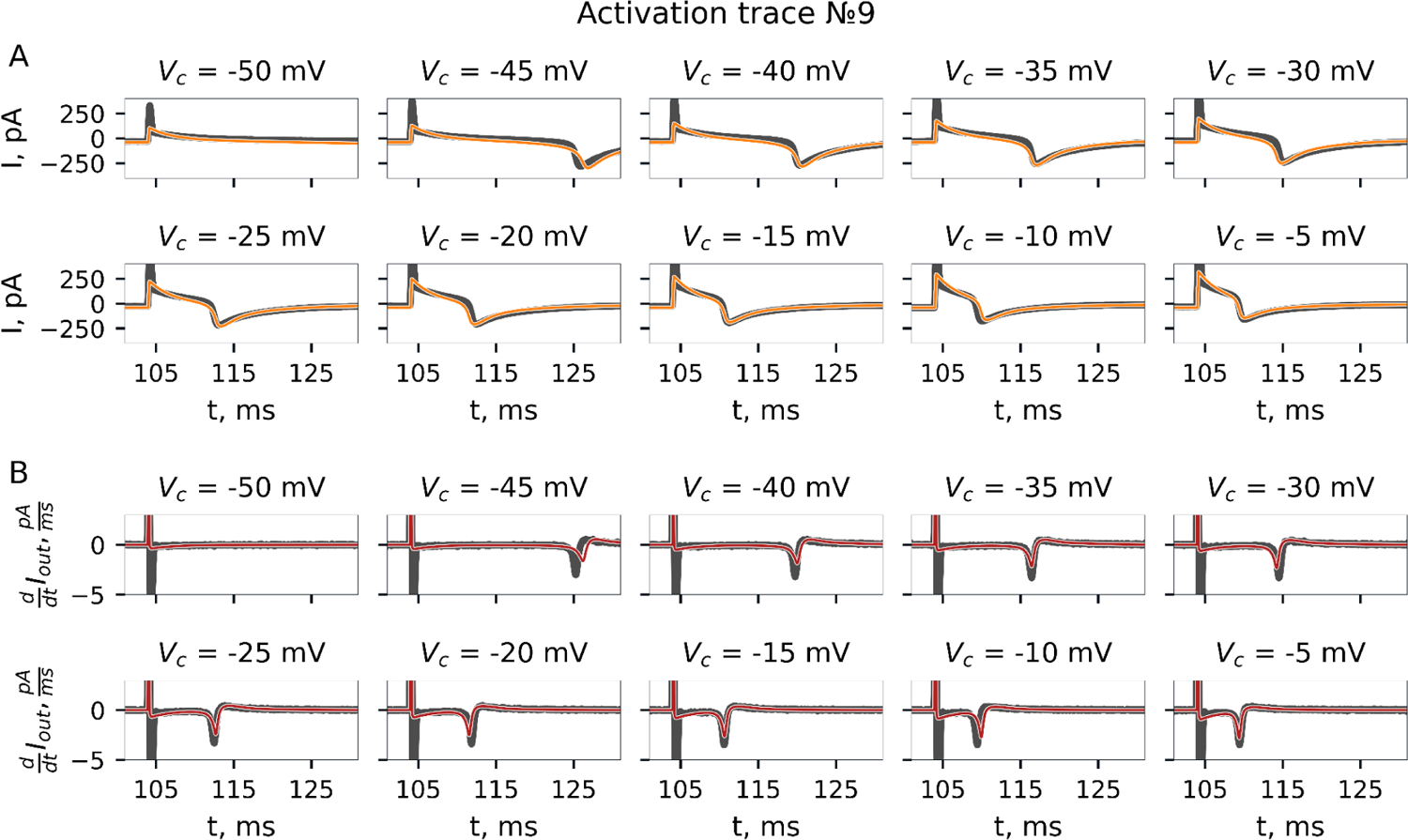
Activation protocol. Recorded current trace and its fit by GA. Experiment №9.

**Fig S16.**
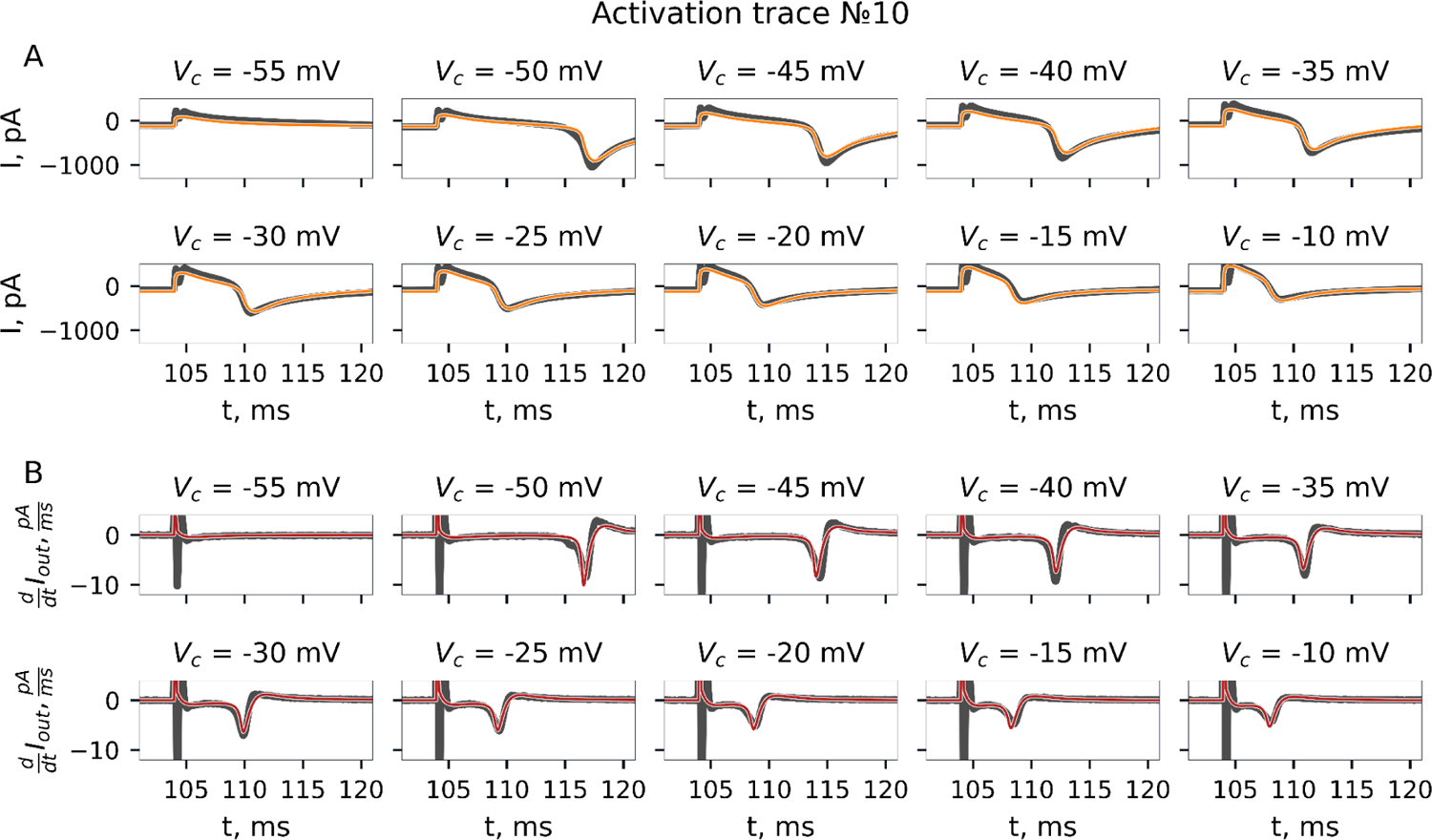
Activation protocol. Recorded current trace and its fit by GA. Experiment №10.

**Fig S17.**
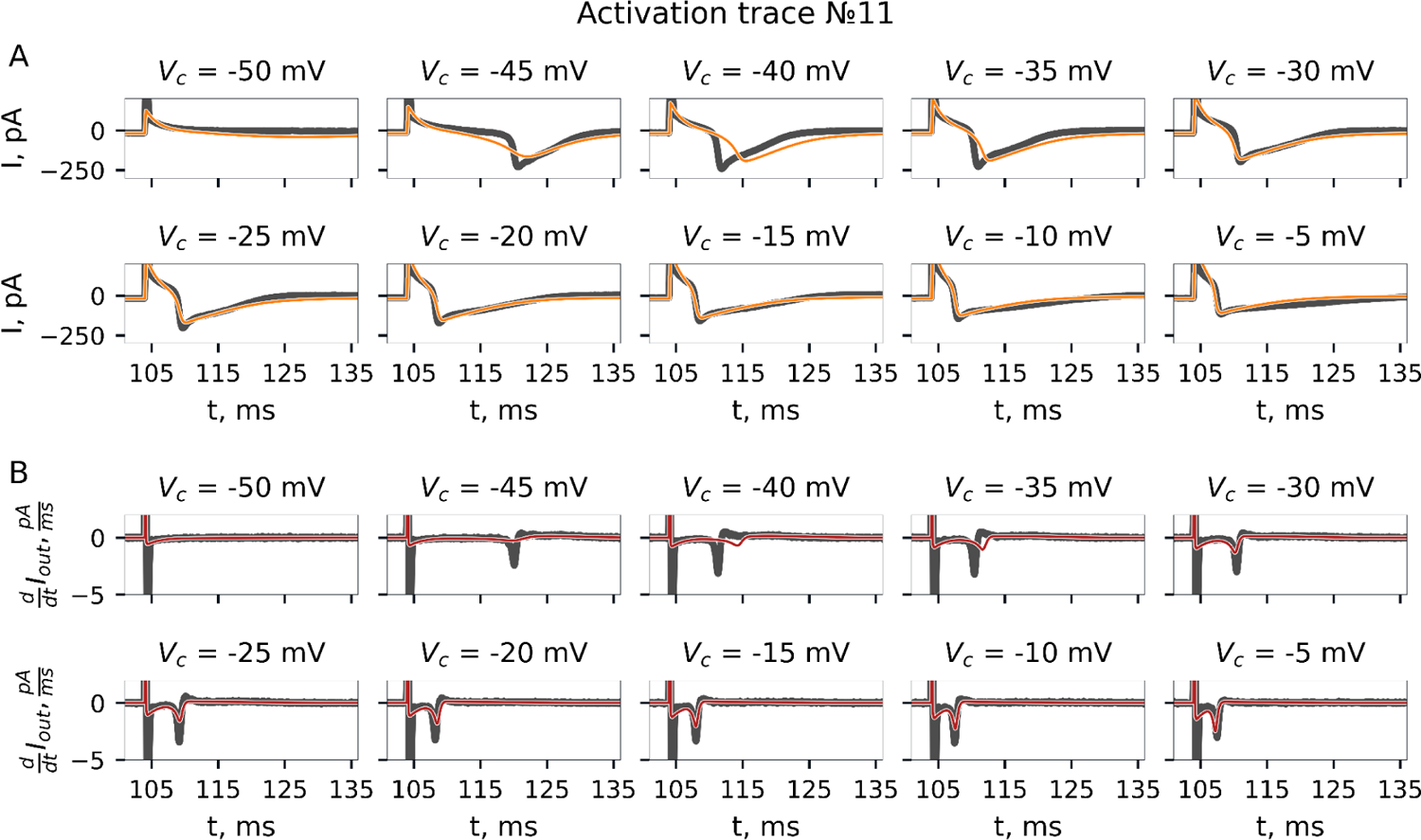
Activation protocol. Recorded current trace and its fit by GA. Experiment №11.

**Fig S18.**
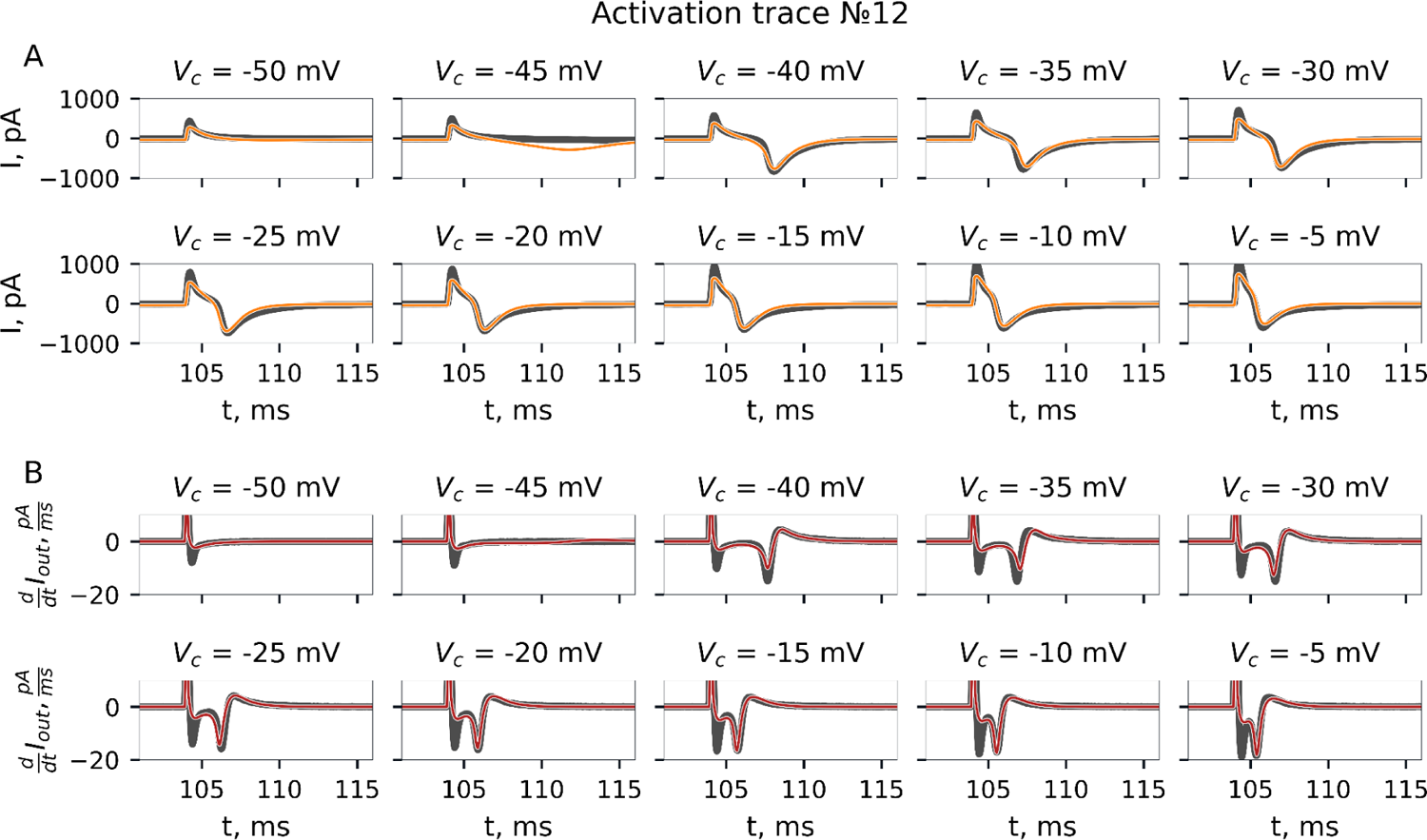
Activation protocol. Recorded current trace and its fit by GA. Experiment №12.

**Fig S19.**
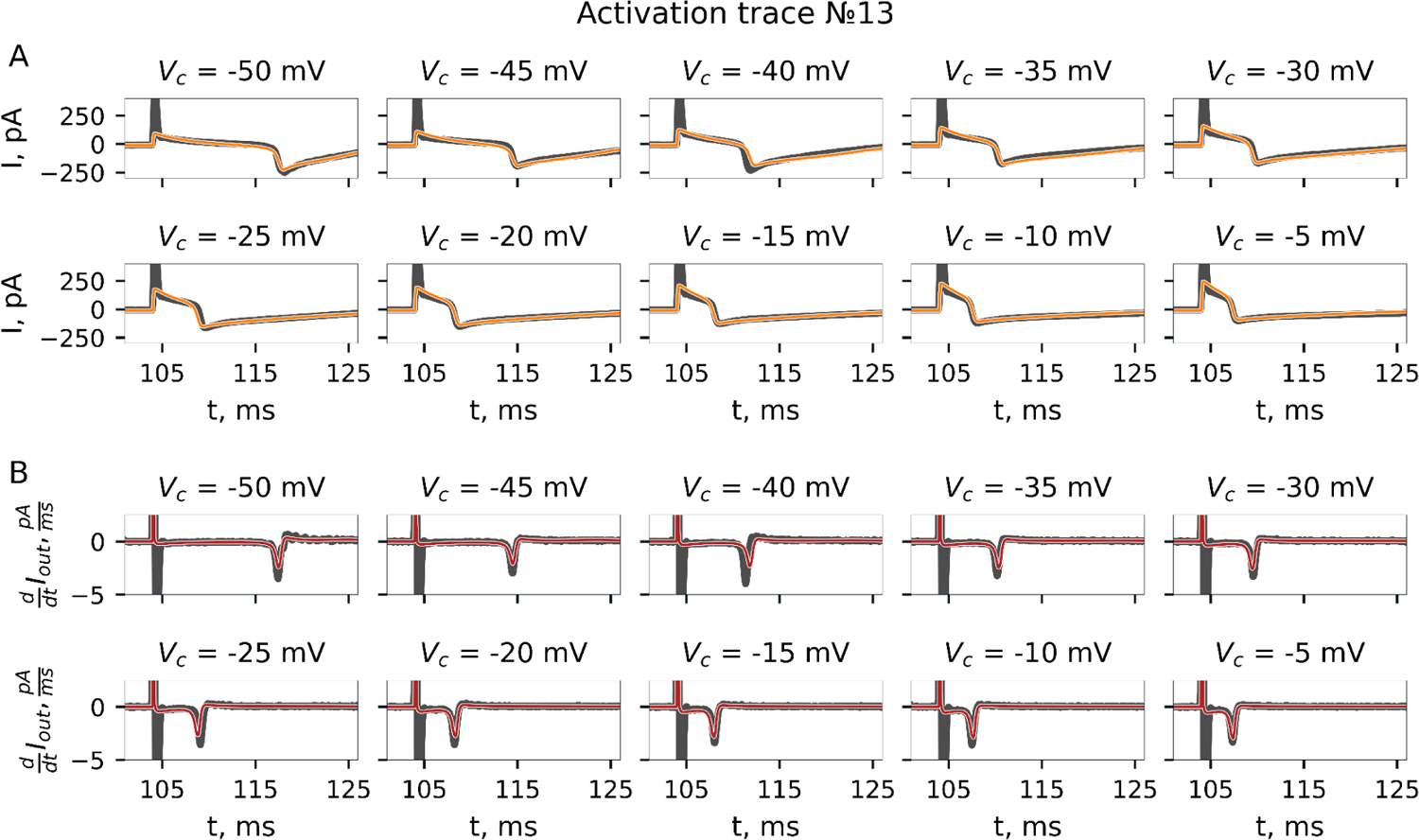
Activation protocol. Recorded current trace and its fit by GA. Experiment №13.

**Fig S20.**
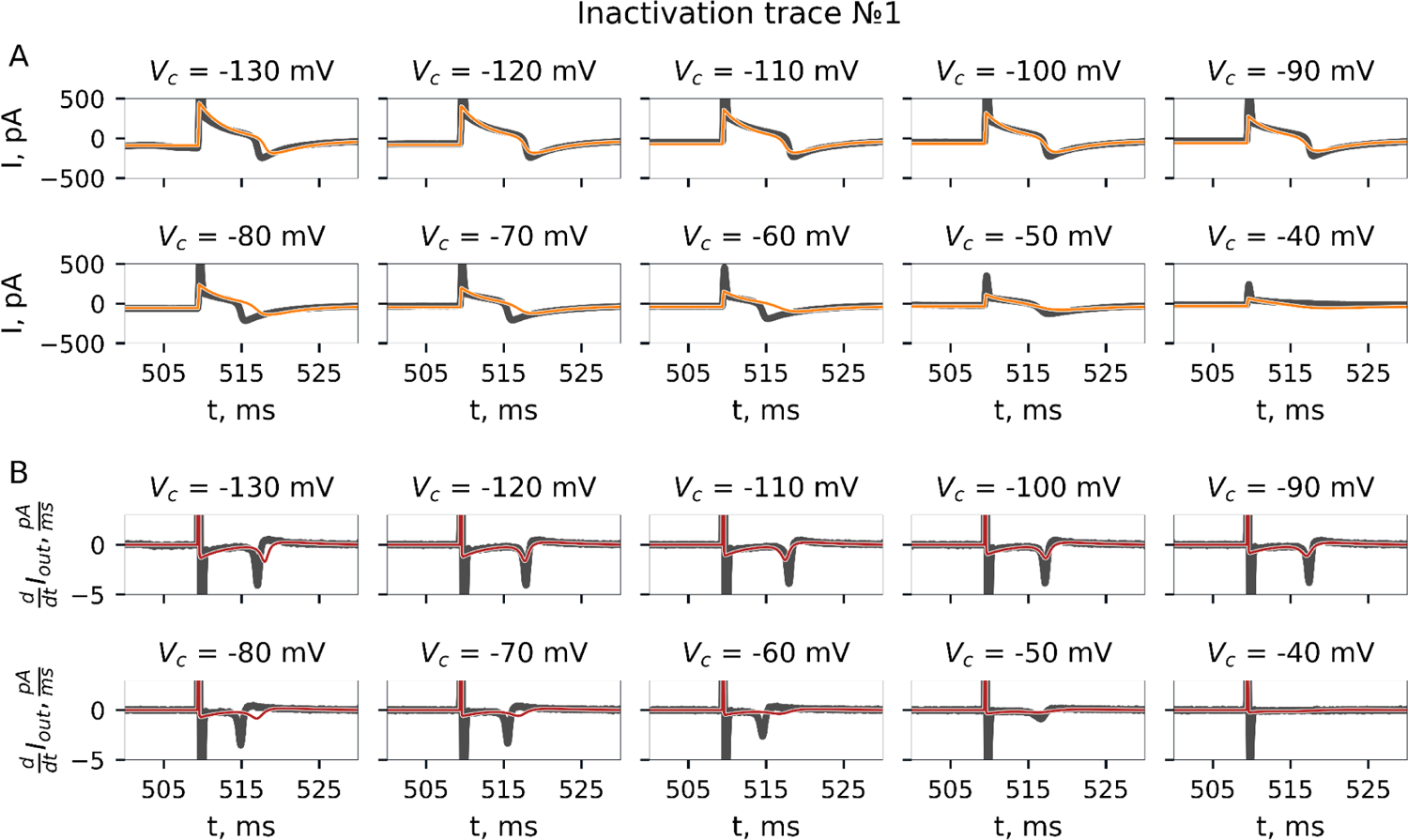
Inactivation protocol. Recorded current trace and its fit by GA. Experiment №1.

**Fig S21.**
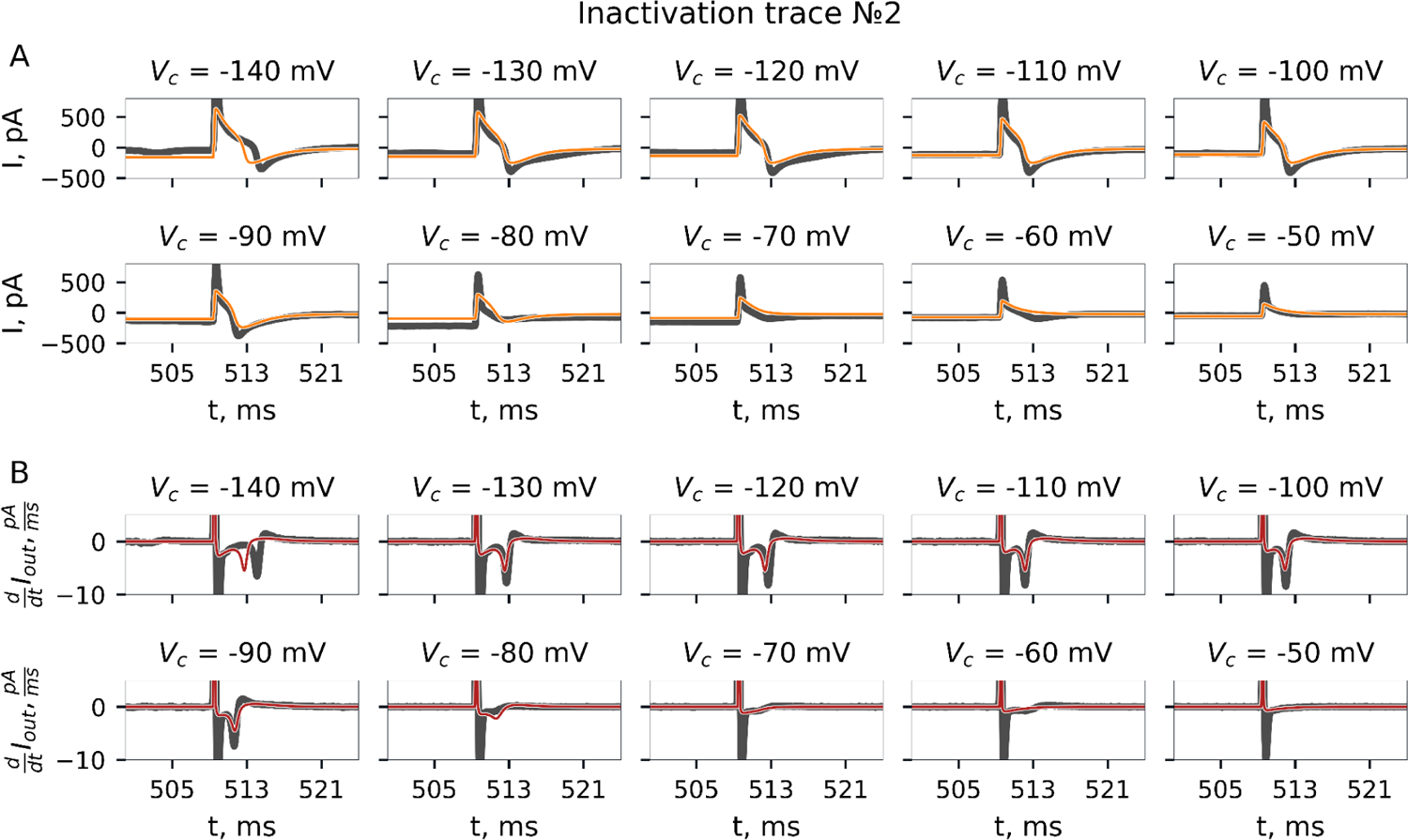
Inactivation protocol. Recorded current trace and its fit by GA. Experiment №2.

**Fig S22.**
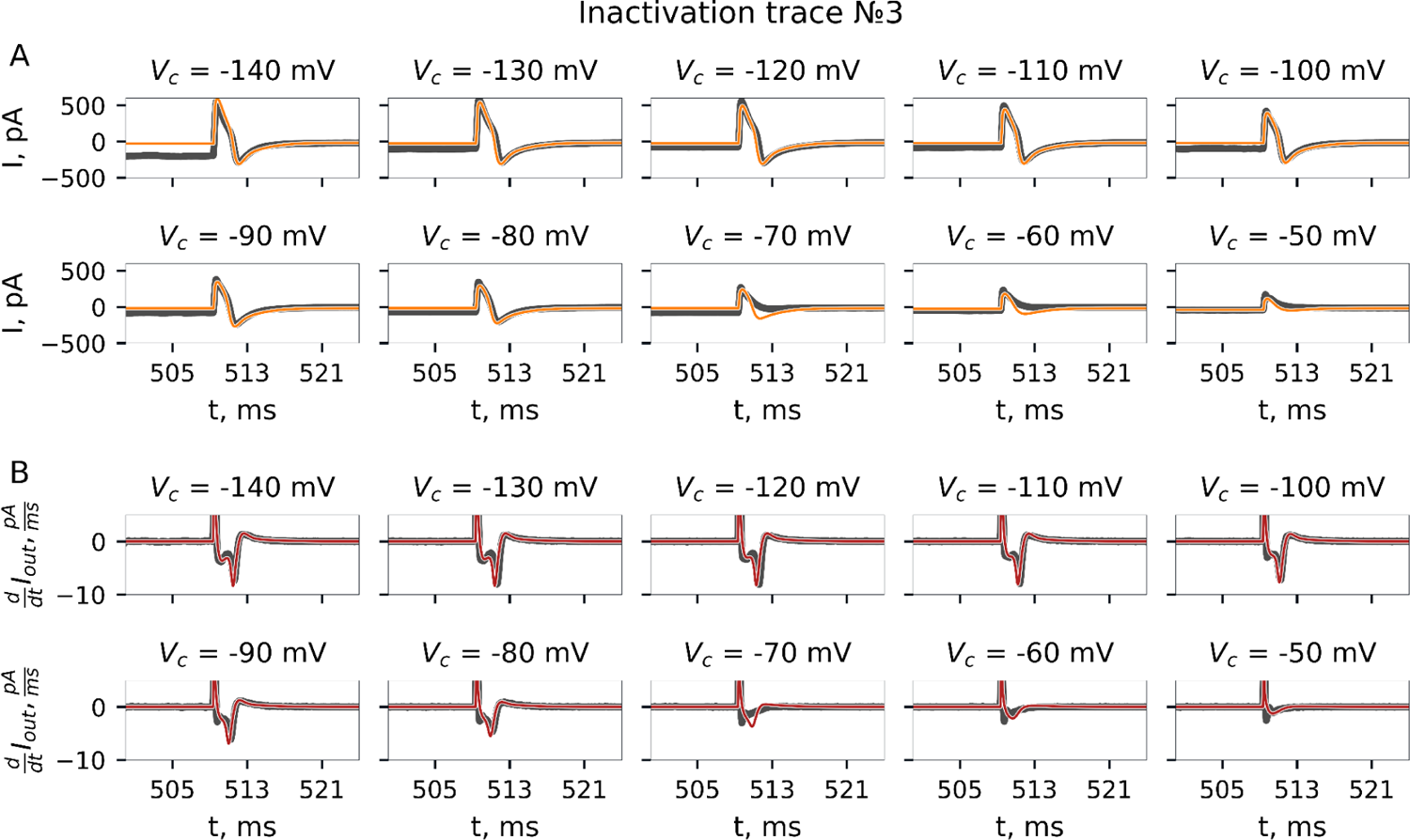
Inactivation protocol. Recorded current trace and its fit by GA. Experiment №3.

**Fig S23.**
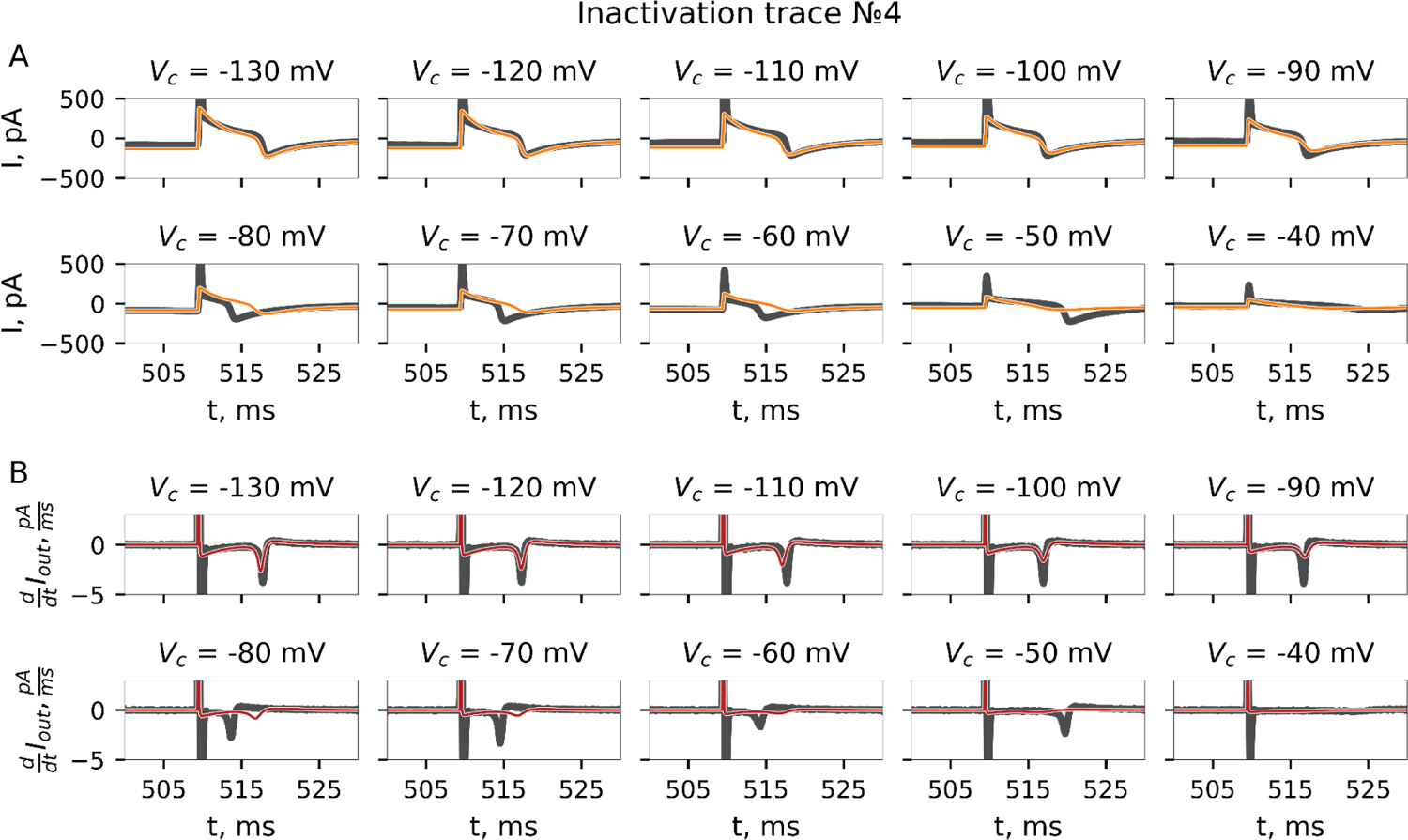
Inactivation protocol. Recorded current trace and its fit by GA. Experiment №4.

**Fig S24.**
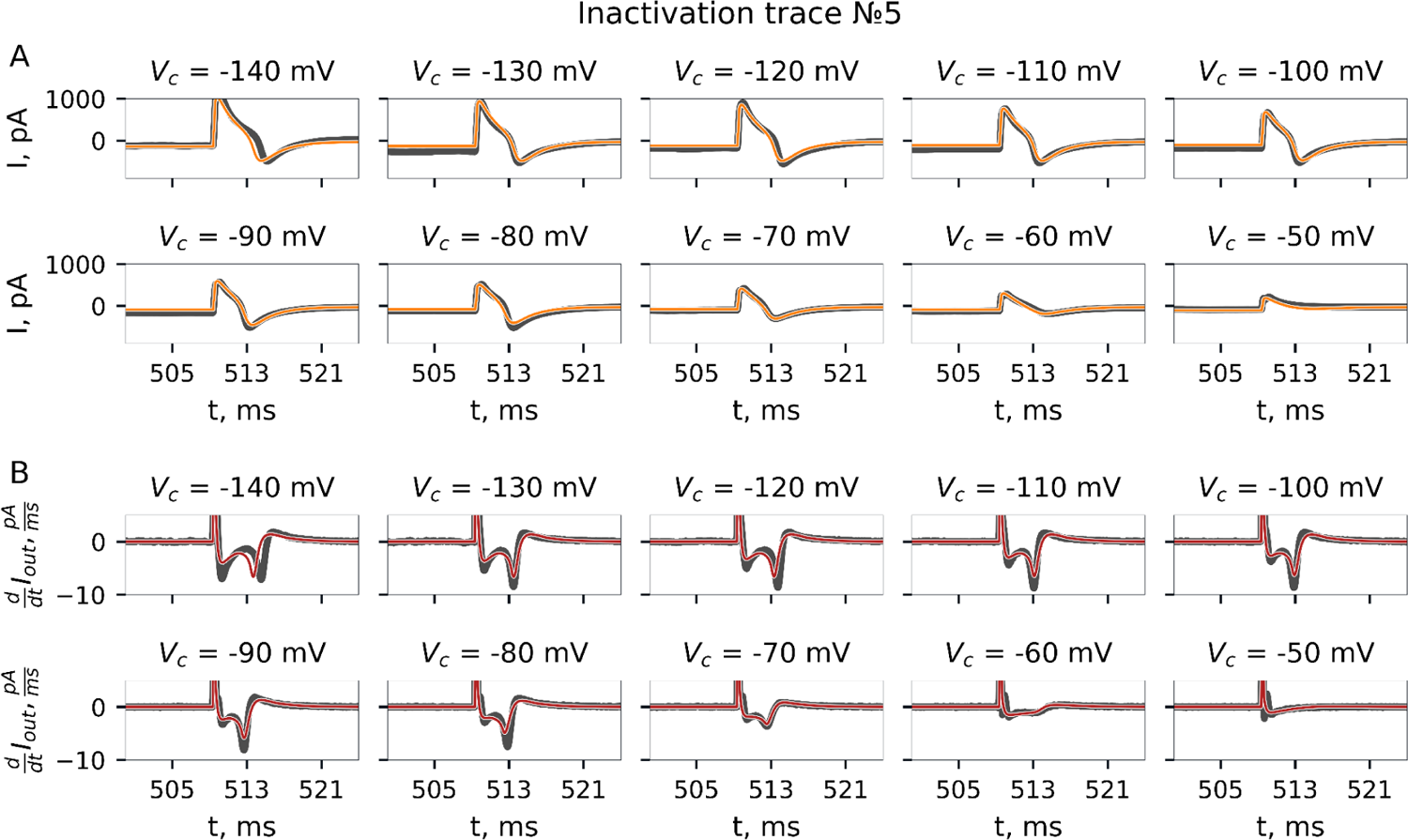
Inactivation protocol. Recorded current trace and its fit by GA. Experiment №5.

**Fig S25.**
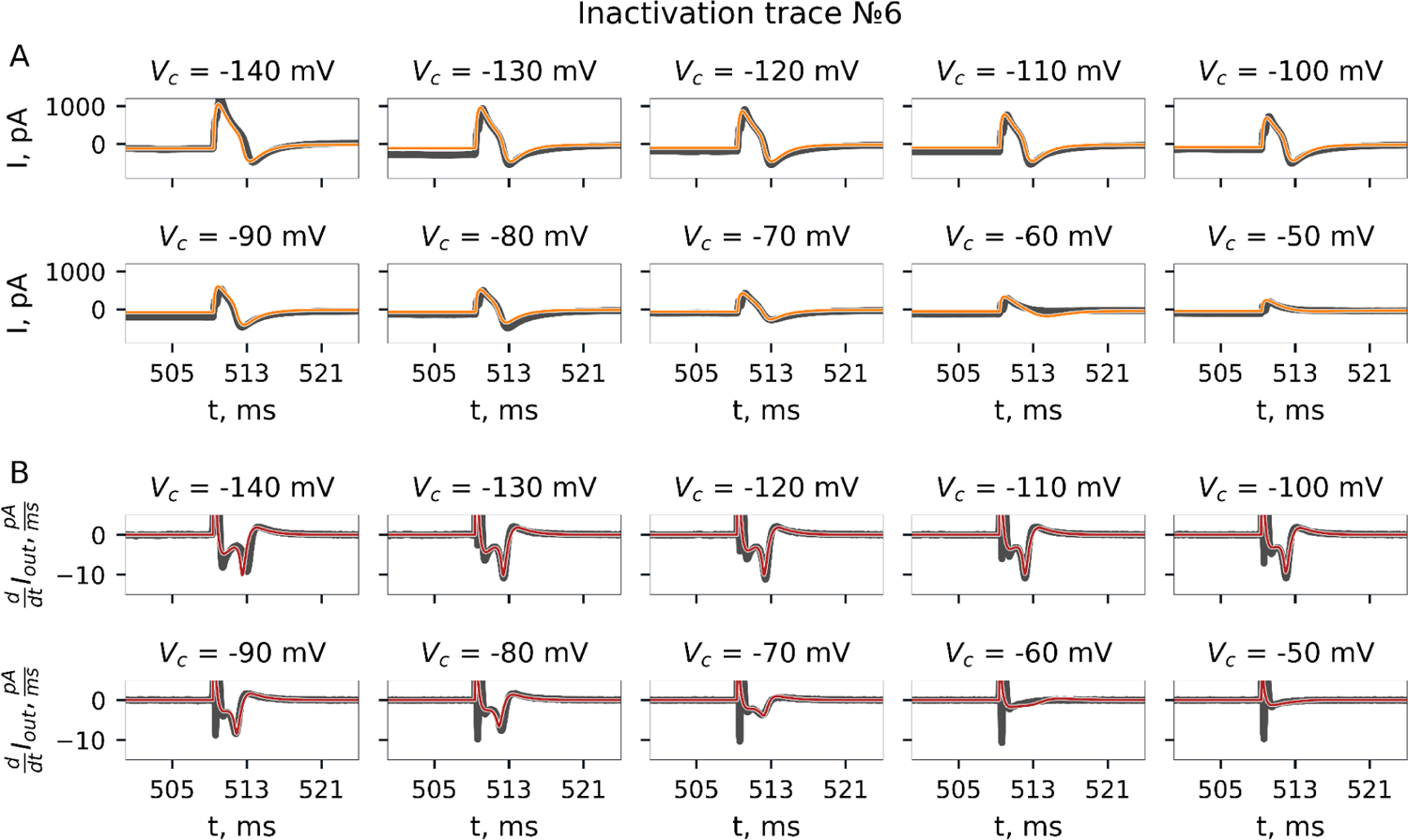
Inactivation protocol. Recorded current trace and its fit by GA. Experiment №6.

**Fig S26.**
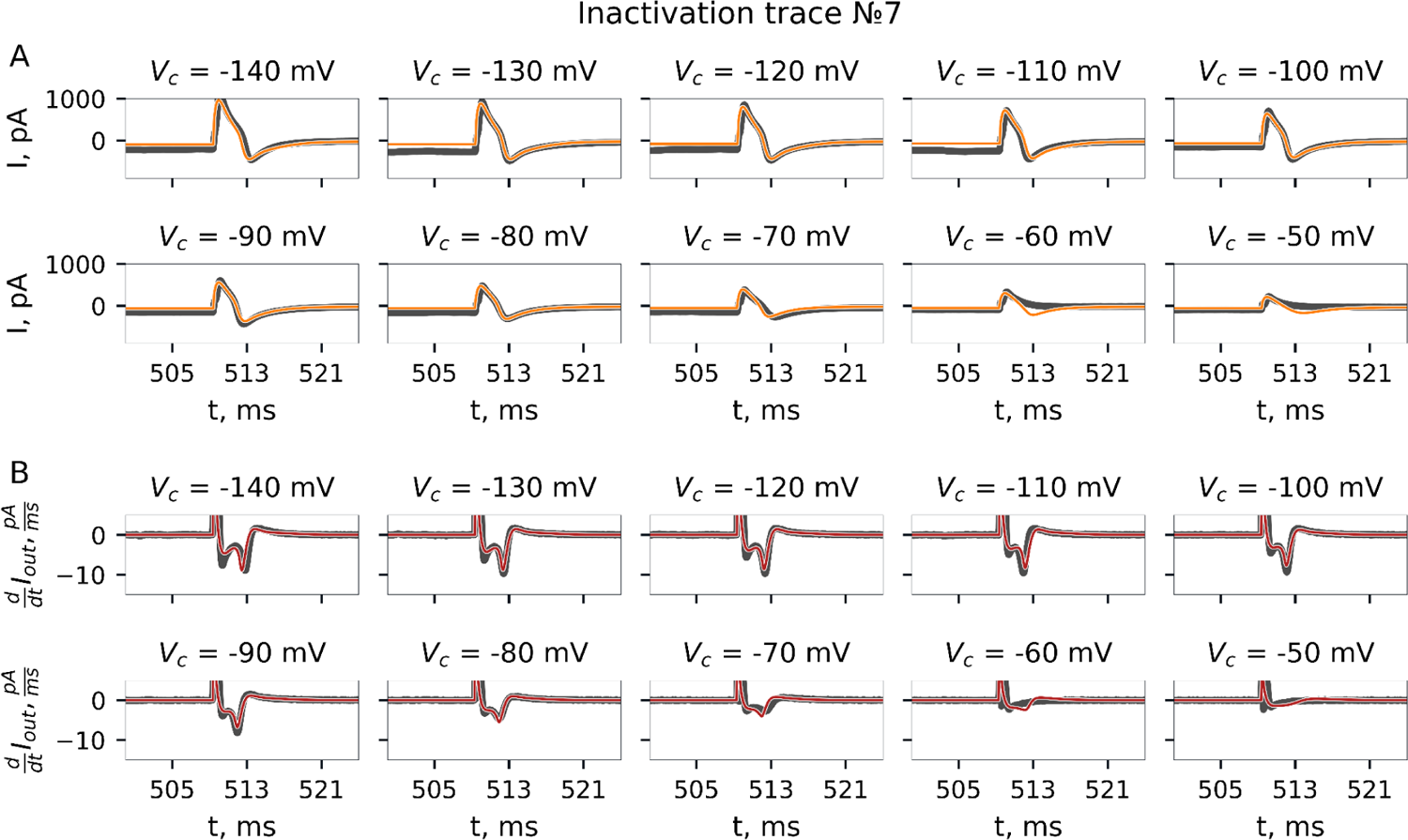
Inactivation protocol. Recorded current trace and its fit by GA. Experiment №7.

**Fig S27.**
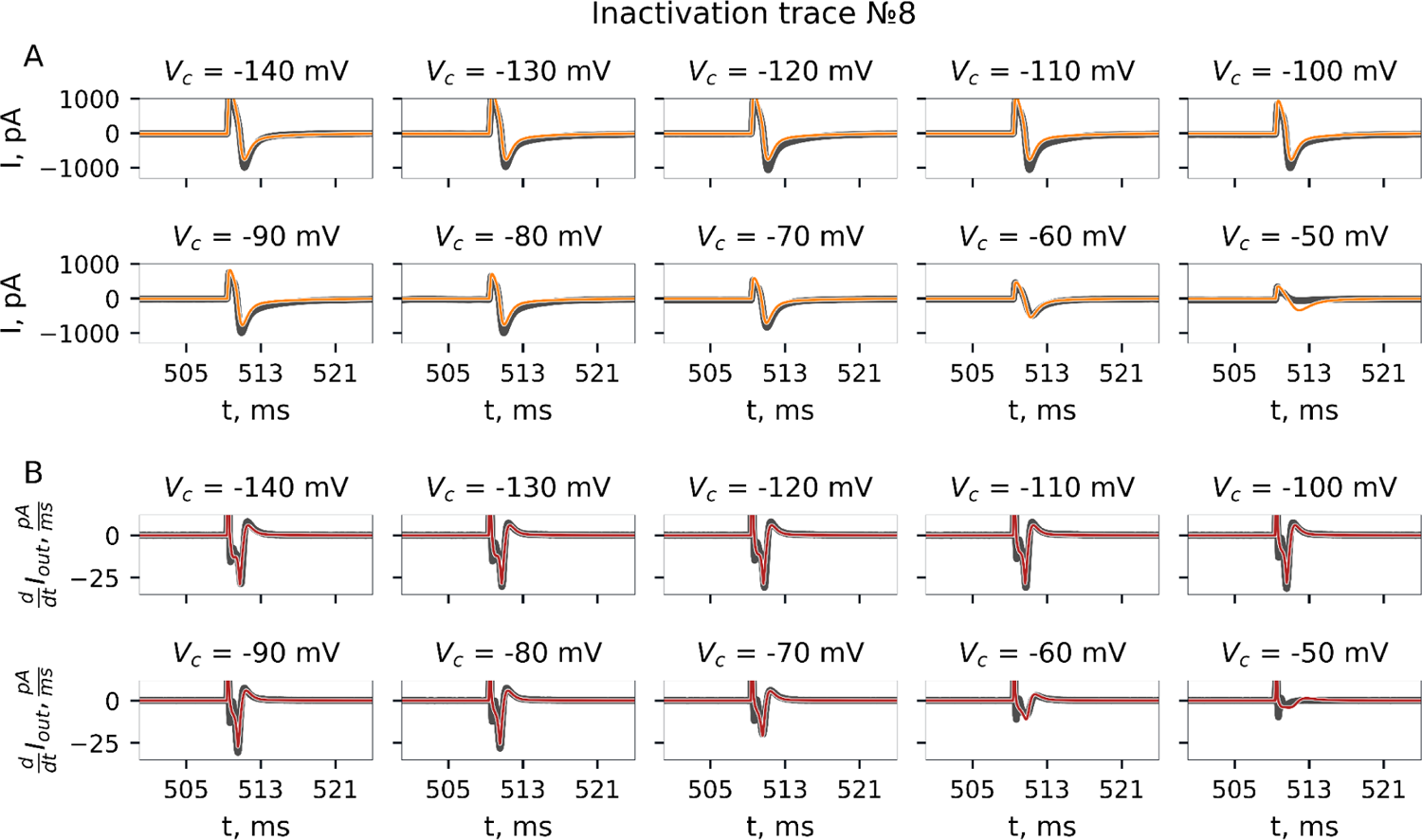
Inactivation protocol. Recorded current trace and its fit by GA. Experiment №8.

**Fig S28.**
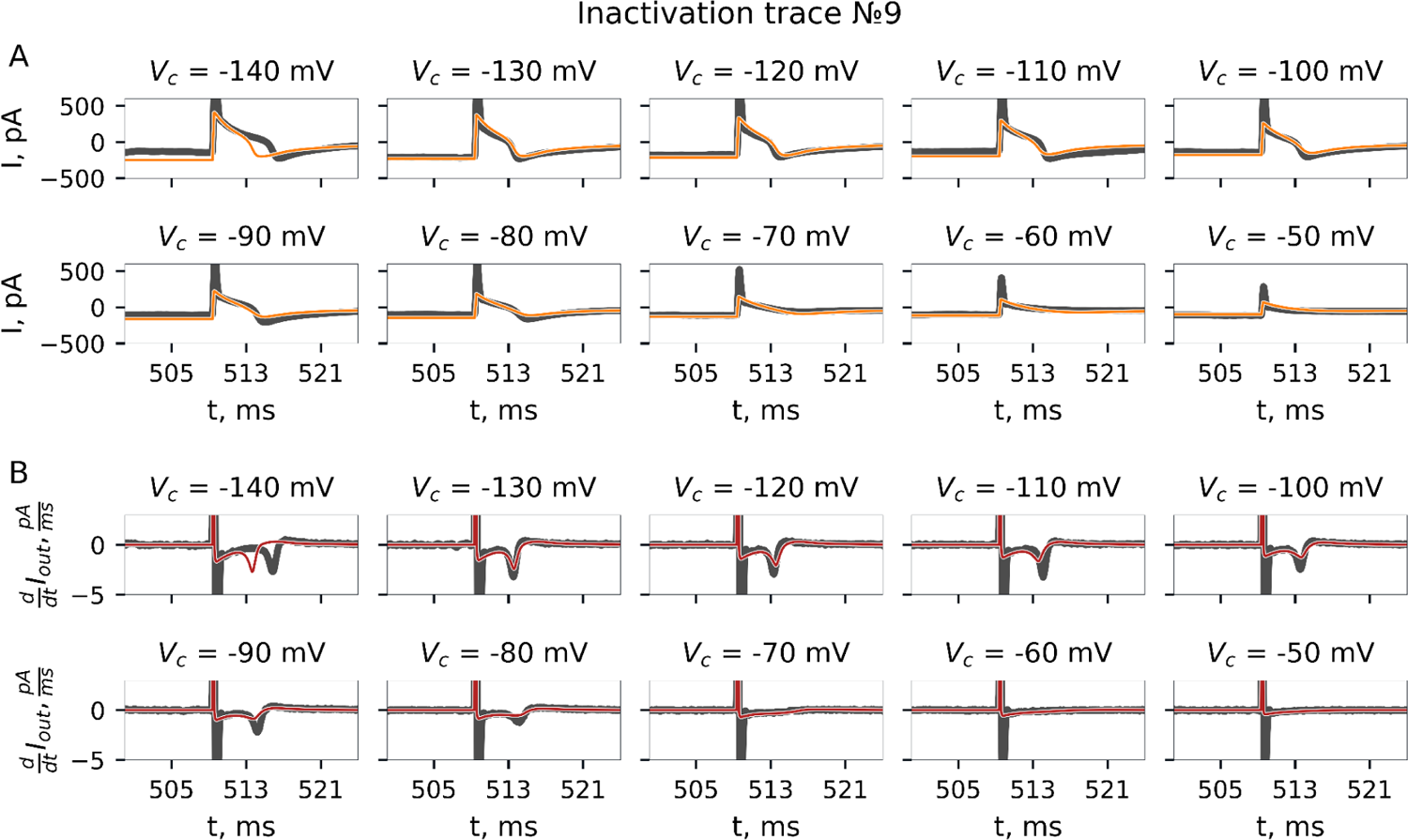
Inactivation protocol. Recorded current trace and its fit by GA. Experiment №9.

**Fig S29.**
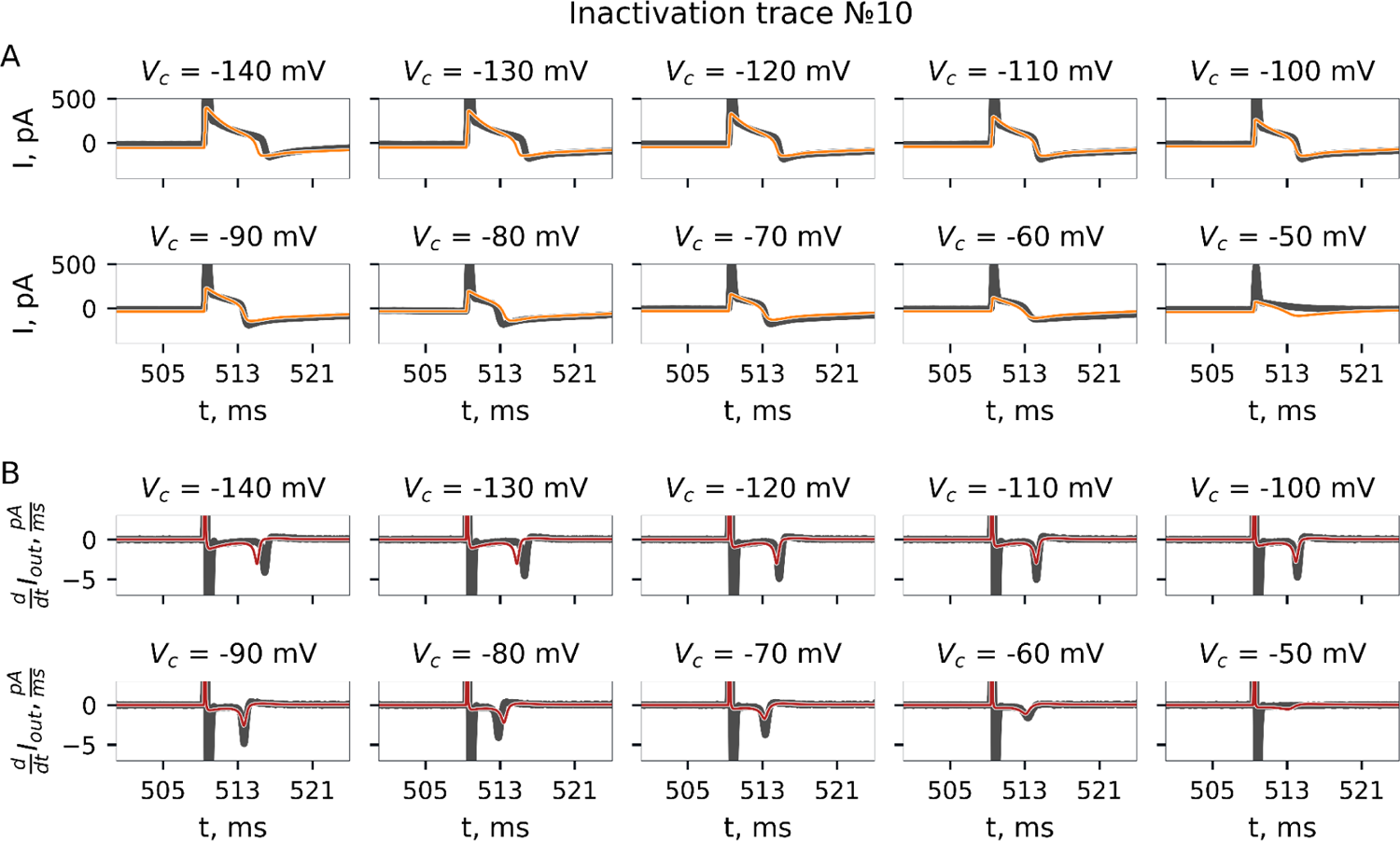
Inactivation protocol. Recorded current trace and its fit by GA. Experiment №10.

**Fig S30.**
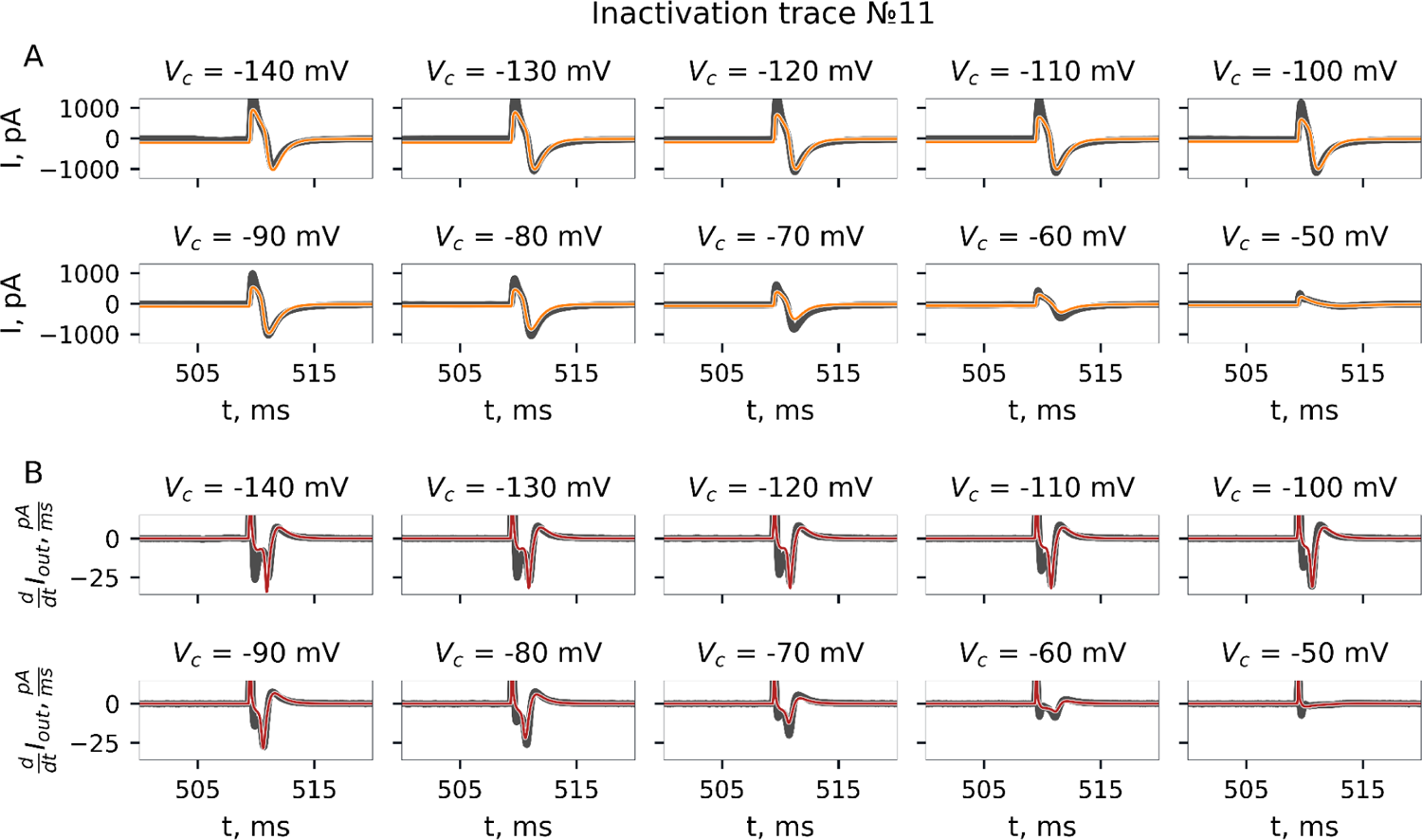
Inactivation protocol. Recorded current trace and its fit by GA. Experiment №11.

**Fig S31.**
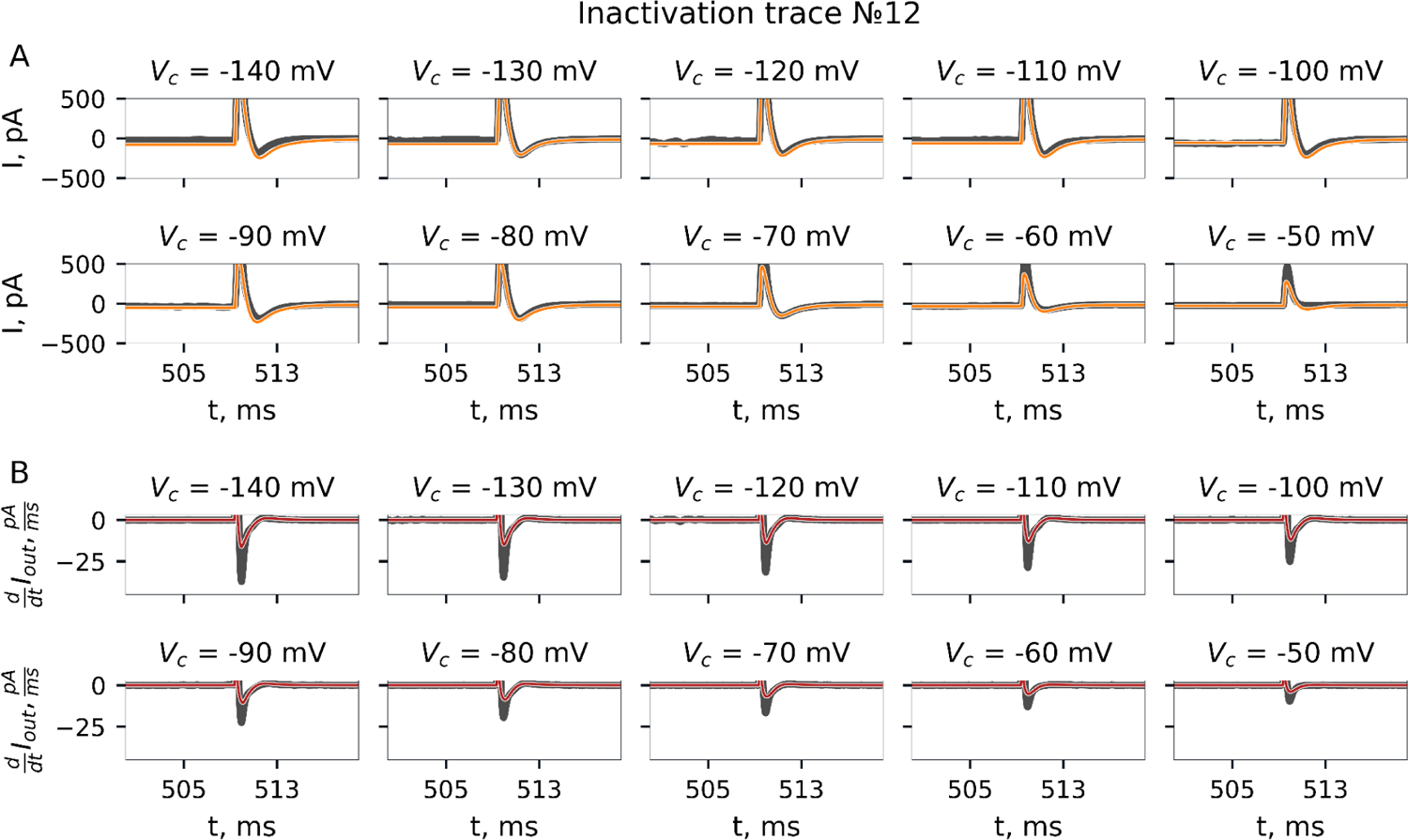
Inactivation protocol. Recorded current trace and its fit by GA. Experiment №12.

**Fig S32.**
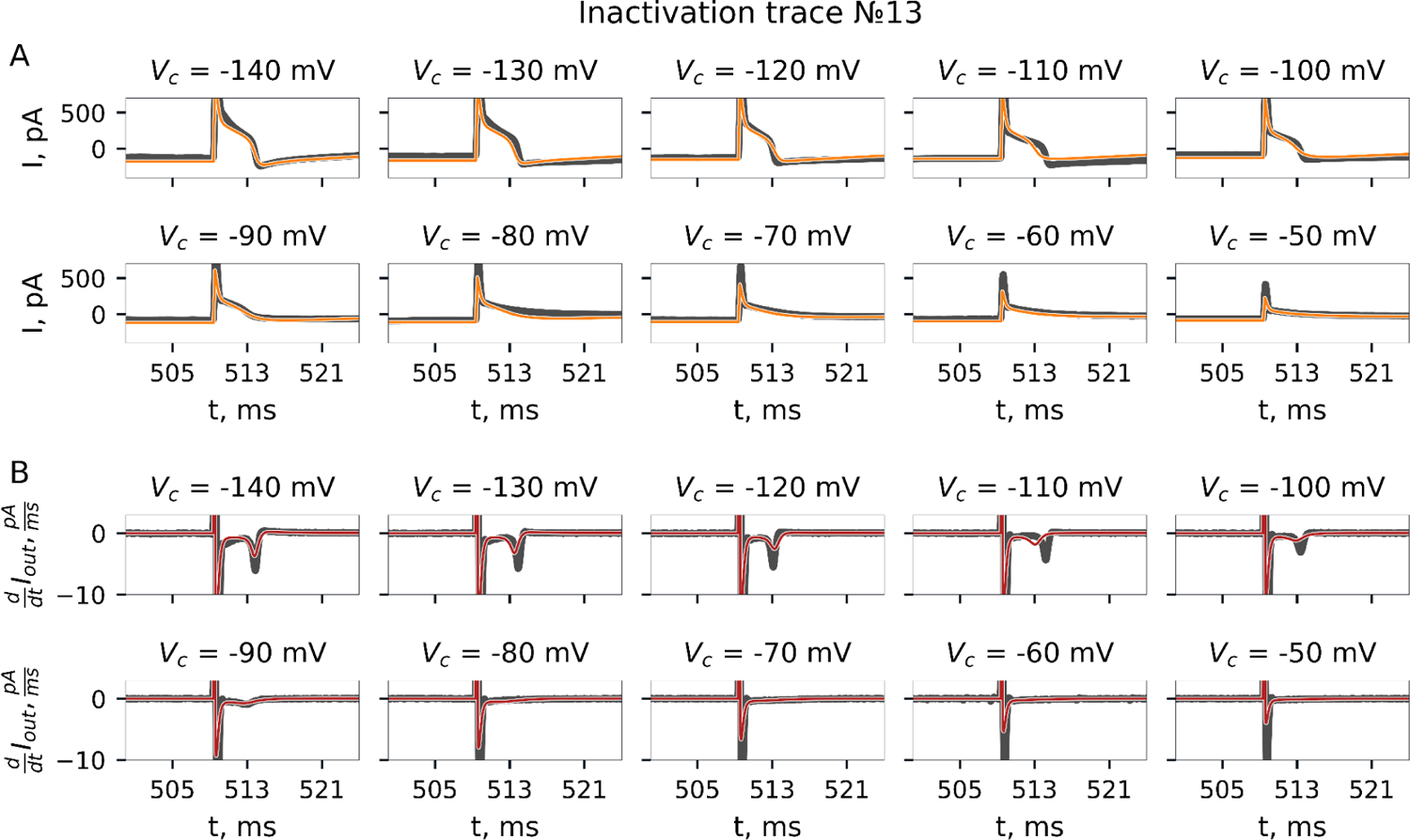
Inactivation protocol. Recorded current trace and its fit by GA. Experiment №13.

## References

Aras KK, Faye NR, Cathey B & Efimov IR (2018). Critical Volume of Human Myocardium Necessary to Maintain Ventricular Fibrillation. Circ Arrhythm Electrophysiol 11, e006692.

Argenziano M, Lambers E, Hong L, Sridhar A, Zhang M, Chalazan B, Menon A, Savio-Galimberti E, Wu JC, Rehman J & Darbar D (2018). Electrophysiologic Characterization of Calcium Handling in Human Induced Pluripotent Stem Cell-Derived Atrial Cardiomyocytes. Stem Cell Reports 10, 1867–1878.

Armstrong CM & Chow RH (1987). Supercharging: a method for improving patch-clamp performance. Biophys J 52, 133–136.

Attwell D, Cohen I, Eisner D, Ohba M & Ojeda C (1979). The steady state TTX-sensitive (“window”) sodium current in cardiac Purkinje fibres. Pflugers Arch 379, 137–142.

Balser JR (1999). Structure and function of the cardiac sodium channels. Cardiovasc Res 42, 327–338.

Banyasz T, Jian Z, Horvath B, Khabbaz S, Izu LT & Chen-Izu Y (2014). Beta-adrenergic stimulation reverses the I Kr-I Ks dominant pattern during cardiac action potential. Pflugers Arch 466, 2067–2076.

Barajas-Martínez H, Hu D, Goodrow RJ Jr, Joyce F & Antzelevitch C (2013). Electrophysiologic characteristics and pharmacologic response of human cardiomyocytes isolated from a patient with hypertrophic cardiomyopathy. Pacing Clin Electrophysiol 36, 1512–1515.

Bartolucci C, Passini E, Hyttinen J, Paci M & Severi S (2020). Simulation of the Effects of Extracellular Calcium Changes Leads to a Novel Computational Model of Human Ventricular Action Potential With a Revised Calcium Handling. Front Physiol 11, 314.

Beaumont J, Roberge FA & Leon LJ (1993). On the interpretation of voltage-clamp data using the Hodgkin-Huxley model. Math Biosci 115, 65–101.

Boyle PM, Zghaib T, Zahid S, Ali RL, Deng D, Franceschi WH, Hakim JB, Murphy MJ, Prakosa A, Zimmerman SL, Ashikaga H, Marine JE, Kolandaivelu A, Nazarian S, Spragg DD, Calkins H & Trayanova NA (2019). Computationally guided personalized targeted ablation of persistent atrial fibrillation. Nat Biomed Eng 3, 870–879.

Burnett SD, Blanchette AD, Chiu WA & Rusyn I (2021). Human induced pluripotent stem cell (iPSC)-derived cardiomyocytes as an in vitro model in toxicology: strengths and weaknesses for hazard identification and risk characterization. Expert Opin Drug Metab Toxicol 17, 887–902.

Burridge PW, Matsa E, Shukla P, Lin ZC, Churko JM, Ebert AD, Lan F, Diecke S, Huber B, Mordwinkin NM, Plews JR, Abilez OJ, Cui B, Gold JD & Wu JC (2014). Chemically defined generation of human cardiomyocytes. Nat Methods 11, 855–860.

Chen Z, Xian W, Bellin M, Dorn T, Tian Q, Goedel A, Dreizehnter L, Schneider CM, Ward-van Oostwaard D, Ng JKM, Hinkel R, Pane LS, Mummery CL, Lipp P, Moretti A, Laugwitz K-L & Sinnecker D (2017). Subtype-specific promoter-driven action potential imaging for precise disease modelling and drug testing in hiPSC-derived cardiomyocytes. Eur Heart J 38, 292–301.

Clancy CE & Kass RS (2005). Inherited and acquired vulnerability to ventricular arrhythmias: cardiac Na+ and K+ channels. Physiol Rev 85, 33–47.

Clerx M, Beattie KA, Gavaghan DJ & Mirams GR (2019). Four Ways to Fit an Ion Channel Model. Biophys J 117, 2420–2437.

Davis RP, Casini S, van den Berg CW, Hoekstra M, Remme CA, Dambrot C, Salvatori D, Oostwaard DW, Wilde AAM, Bezzina CR, Verkerk AO, Freund C & Mummery CL (2012). Cardiomyocytes derived from pluripotent stem cells recapitulate electrophysiological characteristics of an overlap syndrome of cardiac sodium channel disease. Circulation 125, 3079–3091.

Dementyeva EV, Medvedev SP, Kovalenko VR, Vyatkin YV, Kretov EI, Slotvitsky MM, Shtokalo DN, Pokushalov EA & Zakian SM (2019). Applying Patient-Specific Induced Pluripotent Stem Cells to Create a Model of Hypertrophic Cardiomyopathy. Biochemistry 84, 291–298.

Drouin E, Charpentier F, Gauthier C, Laurent K & Le Marec H (1995). Electrophysiologic characteristics of cells spanning the left ventricular wall of human heart: evidence for presence of M cells. J Am Coll Cardiol 26, 185–192.

Dutta S, Chang KC, Beattie KA, Sheng J, Tran PN, Wu WW, Wu M, Strauss DG, Colatsky T & Li Z (2017a). Optimization of an In silico Cardiac Cell Model for Proarrhythmia Risk Assessment. Front Physiol 8, 616.

Dutta S, Mincholé A, Quinn TA & Rodriguez B (2017b). Electrophysiological properties of computational human ventricular cell action potential models under acute ischemic conditions. Prog Biophys Mol Biol 129, 40–52.

Fassina D, M. Costa C, Bishop M, Plank G, Whitaker J, Harding SE & Niederer SA (2023). Assessing the arrhythmogenic risk of engineered heart tissue patches through in silico application on infarcted ventricle models. Comput Biol Med 154, 106550.

Feaster TK, Cadar AG, Wang L, Williams CH, Chun YW, Hempel JE, Bloodworth N, Merryman WD, Lim CC, Wu JC, Knollmann BC & Hong CC (2015). Matrigel Mattress: A Method for the Generation of Single Contracting Human-Induced Pluripotent Stem Cell-Derived Cardiomyocytes. Circ Res 117, 995–1000.

Gardner DJ, Reynolds DR, Woodward CS & Balos CJ (2022). Enabling New Flexibility in the SUNDIALS Suite of Nonlinear and Differential/Algebraic Equation Solvers. ACM Trans Math Softw 48, 1–24.

George AL Jr (2005). Inherited disorders of voltage-gated sodium channels. J Clin Invest 115, 1990–1999.

Glukhov AV, Fedorov VV, Kalish PW, Ravikumar VK, Lou Q, Janks D, Schuessler RB, Moazami N & Efimov IR (2012). Conduction remodeling in human end-stage nonischemic left ventricular cardiomyopathy. Circulation 125, 1835–1847.

Goversen B, van der Heyden MAG, van Veen TAB & de Boer TP (2018). The immature electrophysiological phenotype of iPSC-CMs still hampers in vitro drug screening: Special focus on IK1. Pharmacol Ther 183, 127–136.

Grandi E, Pasqualini FS & Bers DM (2010). A novel computational model of the human ventricular action potential and Ca transient. J Mol Cell Cardiol 48, 112–121.

Groenendaal W, Ortega FA, Kherlopian AR, Zygmunt AC, Krogh-Madsen T & Christini DJ (2015). Cell-specific cardiac electrophysiology models. PLoS Comput Biol 11, e1004242.

Haario H, Laine M, Mira A & Saksman E (2006). DRAM: Efficient adaptive MCMC. Stat Comput 16, 339–354.

Hanck DA & Sheets MF (1992). Time-dependent changes in kinetics of Na+ current in single canine cardiac Purkinje cells. Am J Physiol 262, H1197–H1207.

Herron TJ, Rocha AMD, Campbell KF, Ponce-Balbuena D, Willis BC, Guerrero-Serna G, Liu Q, Klos M, Musa H, Zarzoso M, Bizy A, Furness J, Anumonwo J, Mironov S & Jalife J (2016). Extracellular Matrix–Mediated Maturation of Human Pluripotent Stem Cell–Derived Cardiac Monolayer Structure and Electrophysiological Function. Circ Arrhythm Electrophysiol 9, e003638.

Hespanhol L, Vallio CS, Costa LM & Saragiotto BT (2019). Understanding and interpreting confidence and credible intervals around effect estimates. Braz J Phys Ther 23, 290–301.

Hindmarsh AC, Brown PN, Grant KE, Lee SL, Serban R, Shumaker DE & Woodward CS (2005). SUNDIALS: Suite of nonlinear and differential/algebraic equation solvers. ACM Trans Math Softw 31, 363–396.

Hodgkin AL & Huxley AF (1952). A quantitative description of membrane current and its application to conduction and excitation in nerve. J Physiol 117, 500–544.

Holzem KM, Gomez JF, Glukhov AV, Madden EJ, Koppel AC, Ewald GA, Trenor B & Efimov IR (2016). Reduced response to IKr blockade and altered hERG1a/1b stoichiometry in human heart failure. J Mol Cell Cardiol 96, 82–92.

Horváth A, Lemoine MD, Löser A, Mannhardt I, Flenner F, Uzun AU, Neuber C, Breckwoldt K, Hansen A, Girdauskas E, Reichenspurner H, Willems S, Jost N, Wettwer E, Eschenhagen T & Christ T (2018). Low Resting Membrane Potential and Low Inward Rectifier Potassium Currents Are Not Inherent Features of hiPSC-Derived Cardiomyocytes. Stem Cell Reports 10, 822–833.

Johnstone RH, Chang ETY, Bardenet R, de Boer TP, Gavaghan DJ, Pathmanathan P, Clayton RH & Mirams GR (2016). Uncertainty and variability in models of the cardiac action potential: Can we build trustworthy models? J Mol Cell Cardiol 96, 49–62.

Jonsson MKB, Vos MA, Mirams GR, Duker G, Sartipy P, de Boer TP & van Veen TAB (2012). Application of human stem cell-derived cardiomyocytes in safety pharmacology requires caution beyond hERG. Journal of Molecular and Cellular Cardiology 52, 998–1008. Available at: http://dx.doi.org/10.1016/j.yjmcc.2012.02.002.

Jost N, Virág L, Bitay M, Takács J, Lengyel C, Biliczki P, Nagy Z, Bogáts G, Lathrop DA, Papp JG & Varró A (2005). Restricting excessive cardiac action potential and QT prolongation: a vital role for IKs in human ventricular muscle. Circulation 112, 1392–1399.

Kang C, Badiceanu A, Brennan JA, Gloschat C, Qiao Y, Trayanova NA & Efimov IR (2017). β-adrenergic stimulation augments transmural dispersion of repolarization via modulation of delayed rectifier currents IKs and IKr in the human ventricle. Sci Rep 7, 15922.

Karakikes I, Ameen M, Termglinchan V & Wu JC (2015). Human induced pluripotent stem cell–derived cardiomyocytes. Circ Res 117, 80–88.

Kaufmann SG, Westenbroek RE, Maass AH, Lange V, Renner A, Wischmeyer E, Bonz A, Muck J, Ertl G, Catterall WA, Scheuer T & Maier SKG (2013). Distribution and function of sodium channel subtypes in human atrial myocardium. J Mol Cell Cardiol 61, 133–141.

Kim JJ, Yang L, Lin B, Zhu X, Sun B, Kaplan AD, Bett GCL, Rasmusson RL, London B & Salama G (2015). Mechanism of automaticity in cardiomyocytes derived from human induced pluripotent stem cells. J Mol Cell Cardiol 81, 81–93.

King DR, Entz M 2nd, Blair GA, Crandell I, Hanlon AL, Lin J, Hoeker GS & Poelzing S (2021). The conduction velocity-potassium relationship in the heart is modulated by sodium and calcium. Pflugers Arch 473, 557–571.

King JH, Huang CL-H & Fraser JA (2013). Determinants of myocardial conduction velocity: implications for arrhythmogenesis. Front Physiol 4, 154.

Koivumäki JT, Naumenko N, Tuomainen T, Takalo J, Oksanen M, Puttonen KA, Lehtonen Š, Kuusisto J, Laakso M, Koistinaho J & Tavi P (2018). Structural Immaturity of Human iPSC-Derived Cardiomyocytes: In Silico Investigation of Effects on Function and Disease Modeling. Front Physiol 9, 80.

Kyrozis A & Reichling DB (1995). Perforated-patch recording with gramicidin avoids artifactual changes in intracellular chloride concentration. J Neurosci Methods 57, 27–35.

Lee J, Smaill B & Smith N (2006). Hodgkin–Huxley type ion channel characterization: An improved method of voltage clamp experiment parameter estimation. J Theor Biol 242, 123–134.

Lei CL, Clerx M, Gavaghan DJ & Mirams GR (2022). Model-driven optimal experimental design for calibrating cardiac electrophysiology models. *bioRxiv*2022.11.01.514669. Available at: https://www.biorxiv.org/content/biorxiv/early/2022/11/02/2022.11.01.514669 [Accessed December 28, 2022].

Lei CL, Clerx M, Whittaker DG, Gavaghan DJ, de Boer TP & Mirams GR (2020). Accounting for variability in ion current recordings using a mathematical model of artefacts in voltage-clamp experiments. Philos Trans A Math Phys Eng Sci 378, 20190348.

Lei CL, Wang K, Clerx M, Johnstone RH, Hortigon-Vinagre MP, Zamora V, Allan A, Smith GL, Gavaghan DJ, Mirams GR & Polonchuk L (2017). Tailoring Mathematical Models to Stem-Cell Derived Cardiomyocyte Lines Can Improve Predictions of Drug-Induced Changes to Their Electrophysiology. Front Physiol 8, 986.

Lemoine MD, Mannhardt I, Breckwoldt K, Prondzynski M, Flenner F, Ulmer B, Hirt MN, Neuber C, Horváth A, Kloth B, Reichenspurner H, Willems S, Hansen A, Eschenhagen T & Christ T (2017). Human iPSC-derived cardiomyocytes cultured in 3D engineered heart tissue show physiological upstroke velocity and sodium current density. Sci Rep 7, 5464.

Lian X, Zhang J, Azarin SM, Zhu K, Hazeltine LB, Bao X, Hsiao C, Kamp TJ & Palecek SP (2013). Directed cardiomyocyte differentiation from human pluripotent stem cells by modulating Wnt/β-catenin signaling under fully defined conditions. Nat Protoc 8, 162–175.

Ma J, Guo L, Fiene SJ, Anson BD, Thomson JA, Kamp TJ, Kolaja KL, Swanson BJ & January CT (2011). High purity human-induced pluripotent stem cell-derived cardiomyocytes: electrophysiological properties of action potentials and ionic currents. Am J Physiol Heart Circ Physiol 301, H2006–H2017.

Martinez-Moreno R, Selga E, Riuró H, Carreras D, Parnes M, Srinivasan C, Wangler MF, Pérez GJ, Scornik FS & Brugada R (2020). An SCN1B Variant Affects Both Cardiac-Type (NaV1.5) and Brain-Type (NaV1.1) Sodium Currents and Contributes to Complex Concomitant Brain and Cardiac Disorders. Front Cell Dev Biol 8, 528742.

Millonas MM & Hanck DA (1998). Nonequilibrium response spectroscopy of voltage-sensitive ion channel gating. Biophys J 74, 210–229.

Montnach J, Lorenzini M, Lesage A, Simon I, Nicolas S, Moreau E, Marionneau C, Baró I, De Waard M & Loussouarn G (2021). Computer modeling of whole-cell voltage-clamp analyses to delineate guidelines for good practice of manual and automated patch-clamp. Sci Rep 11, 3282.

Moore JW, Hines M & Harris EM (1984). Compensation for resistance in series with excitable membranes. Biophys J 46, 507–514.

Nagatomo T, Fan Z, Ye B, Tonkovich GS, January CT, Kyle JW & Makielski JC (1998). Temperature dependence of early and late currents in human cardiac wild-type and long Q-T DeltaKPQ Na+ channels. Am J Physiol 275, H2016–H2024.

Nguyen HX, Wu T, Needs D, Zhang H, Perelli RM, DeLuca S, Yang R, Tian M, Landstrom AP, Henriquez C & Bursac N (2022). Engineered bacterial voltage-gated sodium channel platform for cardiac gene therapy. Nat Commun 13, 620.

Nguyen TP, Wang DW, Rhodes TH & George AL Jr (2008). Divergent biophysical defects caused by mutant sodium channels in dilated cardiomyopathy with arrhythmia. Circ Res 102, 364–371.

O’Hara T, Virág L, Varró A & Rudy Y (2011). Simulation of the Undiseased Human Cardiac Ventricular Action Potential: Model Formulation and Experimental Validation. PLoS Computational Biology 7, e1002061. Available at: http://dx.doi.org/10.1371/journal.pcbi.1002061.

Paci M, Passini E, Klimas A, Severi S, Hyttinen J, Rodriguez B & Entcheva E (2020). All-Optical Electrophysiology Refines Populations of In Silico Human iPSC-CMs for Drug Evaluation. Biophys J 118, 2596–2611.

Plank G, Loewe A, Neic A, Augustin C, Huang Y-L, Gsell MAF, Karabelas E, Nothstein M, Prassl AJ, Sánchez J, Seemann G & Vigmond EJ (2021). The openCARP simulation environment for cardiac electrophysiology. Comput Methods Programs Biomed 208, 106223.

Podgurskaya AD, Tsvelaya VA, Slotvitsky MM, Dementyeva EV, Valetdinova KR & Agladze KI (2019). The Use of iPSC-Derived Cardiomyocytes and Optical Mapping for Erythromycin Arrhythmogenicity Testing. Cardiovasc Toxicol 19, 518–528.

Prè D, Nestor MW, Sproul AA, Jacob S, Koppensteiner P, Chinchalongporn V, Zimmer M, Yamamoto A, Noggle SA & Arancio O (2014). A time course analysis of the electrophysiological properties of neurons differentiated from human induced pluripotent stem cells (iPSCs). PLoS One 9, e103418.

Quan W & Rudy Y (1990). Unidirectional block and reentry of cardiac excitation: a model study. Circ Res 66, 367–382.

Rush S & Larsen H (1978). A practical algorithm for solving dynamic membrane equations. IEEE Trans Biomed Eng 25, 389–392.

Sakakibara Y, Furukawa T, Singer DH, Jia H, Backer CL, Arentzen CE & Wasserstrom JA (1993a). Sodium current in isolated human ventricular myocytes. Am J Physiol 265, H1301–H1309.

Sakakibara Y, Furukawa T, Singer DH, Jia H, Backer CL, Arentzen CE & Wasserstrom JA (1993b). Sodium current in isolated human ventricular myocytes. American Journal of Physiology-Heart and Circulatory Physiology 265, H1301–H1309. Available at: http://dx.doi.org/10.1152/ajpheart.1993.265.4.h1301.

Sakakibara Y, Wasserstrom JA, Furukawa T, Jia H, Arentzen CE, Hartz RS & Singer DH (1992). Characterization of the sodium current in single human atrial myocytes. Circulation Research 71, 535–546. Available at: http://dx.doi.org/10.1161/01.res.71.3.535.

Salvatier J, Wiecki TV & Fonnesbeck C (2016). Probabilistic programming in Python using PyMC3. PeerJ Comput Sci 2, e55.

Satin J, Kehat I, Caspi O, Huber I, Arbel G, Itzhaki I, Magyar J, Schroder EA, Perlman I & Gepstein L (2004). Mechanism of spontaneous excitability in human embryonic stem cell derived cardiomyocytes. The Journal of Physiology 559, 479–496. Available at: http://dx.doi.org/10.1113/jphysiol.2004.068213.

Seibertz F, Rapedius M, Fakuade FE, Tomsits P, Liutkute A, Cyganek L, Becker N, Majumder R, Clauß S, Fertig N & Voigt N (2022). A modern automated patch-clamp approach for high throughput electrophysiology recordings in native cardiomyocytes. Commun Biol 5, 969.

Selga E, Sendfeld F, Martinez-Moreno R, Medine CN, Tura-Ceide O, Wilmut SI, Pérez GJ, Scornik FS, Brugada R & Mills NL (2018). Sodium channel current loss of function in induced pluripotent stem cell-derived cardiomyocytes from a Brugada syndrome patient. J Mol Cell Cardiol 114, 10–19.

Shaheen N, Shiti A, Huber I, Shinnawi R, Arbel G, Gepstein A, Setter N, Goldfracht I, Gruber A, Chorna SV & Gepstein L (2018). Human Induced Pluripotent Stem Cell-Derived Cardiac Cell Sheets Expressing Genetically Encoded Voltage Indicator for Pharmacological and Arrhythmia Studies. Stem Cell Reports 10, 1879–1894.

Sheets MF & Ten Eick RE (1994). Whole-Cell Voltage Clamp of Cardiac Sodium Current. Methods in Neurosciences169–188. Available at: http://dx.doi.org/10.1016/b978-0-12-185287-0.50015-6.

Sherman AJ, Shrier A & Cooper E (1999). Series resistance compensation for whole-cell patch-clamp studies using a membrane state estimator. Biophys J 77, 2590–2601.

Slotvitsky M, Berezhnoy A, Scherbina S, Rimskaya B, Tsvelaya V, Balashov V, Efimov AE, Agapov I & Agladze K (2022). Polymer Kernels as Compact Carriers for Suspended Cardiomyocytes. Micromachines (Basel); DOI: 10.3390/mi14010051.

Slotvitsky MM, Tsvelaya VA, Podgurskaya AD & Agladze KI (2020). Formation of an electrical coupling between differentiating cardiomyocytes. Sci Rep 10, 7774.

Smith RC (2013). Uncertainty Quantification: Theory, Implementation, and Applications. SIAM.

Starmer C (2003). What happens when cardiac Na channel function is compromised? 2. Numerical studies of the vulnerable period in tissue altered by drugs. Cardiovascular Research 57, 1062–1071. Available at: http://dx.doi.org/10.1016/s0008-6363(02)00727-7.

Strickholm A (1995a). A single electrode voltage, current- and patch-clamp amplifier with complete stable series resistance compensation. Journal of Neuroscience Methods 61, 53–66. Available at: http://dx.doi.org/10.1016/0165-0270(95)00021-l.

Strickholm A (1995b). A supercharger for single electrode voltage and current clamping. J Neurosci Methods 61, 47–52.

Szentandrássy N, Farkas V, Bárándi L, Hegyi B, Ruzsnavszky F, Horváth B, Bányász T, Magyar J, Márton I & Nánási PP (2012). Role of action potential configuration and the contribution of C^2+^a and K^+^ currents to isoprenaline-induced changes in canine ventricular cells. Br J Pharmacol 167, 599–611.

Tan SH & Ye L (2018). Maturation of Pluripotent Stem Cell-Derived Cardiomyocytes: a Critical Step for Drug Development and Cell Therapy. J Cardiovasc Transl Res 11, 375–392.

Terrenoire C, Wang K, Chan Tung KW, Chung WK, Pass RH, Lu JT, Jean J-C, Omari A, Sampson KJ, Kotton DN, Keller G & Kass RS (2013). Induced pluripotent stem cells used to reveal drug actions in a long QT syndrome family with complex genetics. Journal of General Physiology 141, 61–72. Available at: http://dx.doi.org/10.1085/jgp.201210899.

Tomek J, Bueno-Orovio A, Passini E, Zhou X, Minchole A, Britton O, Bartolucci C, Severi S, Shrier A, Virag L, Varro A & Rodriguez B (2019). Development, calibration, and validation of a novel human ventricular myocyte model in health, disease, and drug block. Elife; DOI: 10.7554/eLife.48890.

Tse G & Yeo JM (2015). Conduction abnormalities and ventricular arrhythmogenesis: The roles of sodium channels and gap junctions. Int J Cardiol Heart Vasc 9, 75–82.

Tusscher KHWJT, ten Tusscher KHWJ, Noble D, Noble PJ & Panfilov AV (2004). A model for human ventricular tissue. American Journal of Physiology-Heart and Circulatory Physiology 286, H1573–H1589. Available at: http://dx.doi.org/10.1152/ajpheart.00794.2003.

Valdivia CR, Chu WW, Pu J, Foell JD, Haworth RA, Wolff MR, Kamp TJ & Makielski JC (2005). Increased late sodium current in myocytes from a canine heart failure model and from failing human heart. Journal of Molecular and Cellular Cardiology 38, 475–483. Available at: http://dx.doi.org/10.1016/j.yjmcc.2004.12.012.

Veerman CC, Wilde AAM & Lodder EM (2015). The cardiac sodium channel gene SCN5A and its gene product NaV1.5: Role in physiology and pathophysiology. Gene 573, 177–187.

Volders PGA, Stengl M, van Opstal JM, Gerlach U, Spätjens RLHMG, Beekman JDM, Sipido KR & Vos MA (2003). Probing the contribution of IKs to canine ventricular repolarization: key role for beta-adrenergic receptor stimulation. Circulation 107, 2753–2760.

Wagner S, Dybkova N, Rasenack ECL, Jacobshagen C, Fabritz L, Kirchhof P, Maier SKG, Zhang T, Hasenfuss G, Brown JH, Bers DM & Maier LS (2006). Ca2+/calmodulin-dependent protein kinase II regulates cardiac Na+ channels. J Clin Invest 116, 3127–3138.

Wand MP & Jones MC (1994). Kernel Smoothing. CRC Press.

Wang DW, Viswanathan PC, Balser JR, George AL Jr & Benson DW (2002). Clinical, genetic, and biophysical characterization of SCN5A mutations associated with atrioventricular conduction block. Circulation 105, 341–346.

Weiss JN, Qu Z & Shivkumar K (2017). Electrophysiology of Hypokalemia and Hyperkalemia. Circ Arrhythm Electrophysiol; DOI: 10.1161/CIRCEP.116.004667.

William H. Press, Teukolsky SA, Vetterling WT & Flannery BP (2007). Numerical Recipes 3rd Edition: The Art of Scientific Computing. Cambridge University Press.

Willms AR, Baro DJ, Harris-Warrick RM & Guckenheimer J (1999). An improved parameter estimation method for Hodgkin-Huxley models. J Comput Neurosci 6, 145–168.

Wilson JR, Clark RB, Banderali U & Giles WR (2011). Measurement of the membrane potential in small cells using patch clamp methods. Channels 5, 530–537.

Windisch H, Ahammer H, Schaffer P, Müller W & Platzer D (1995). Optical multisite monitoring of cell excitation phenomena in isolated cardiomyocytes. Pflugers Arch 430, 508–518.

Zhu H, Scharnhorst KS, Stieg AZ, Gimzewski JK, Minami I, Nakatsuji N, Nakano H & Nakano A (2017). Two dimensional electrophysiological characterization of human pluripotent stem cell-derived cardiomyocyte system. Sci Rep 7, 43210.

## References

Axon Instruments Inc., “Axopatch 200B patch clamp theory and operation.” https://www.autom8.com/wp-content/uploads/2016/07/Axopatch-200B.pdf, 1997–1999. Accessed: 2023-05-04.

